# Human senataxin uses distinct peripheral elements to unwind R-loops and act on RNA polymerase II

**DOI:** 10.64898/2026.11.01.763451

**Authors:** Matyas Lanicek, Katerina Sedova, Zdenka Hasanova, Odil Porrua, Marek Sebesta

## Abstract

Persistent R-loops and paused or stalled RNA polymerase II (RNAPII) elongation complexes can threaten genome stability. Senataxin (SETX) contributes to R-loop resolution and RNAPII regulation, but their intimate connection during transcription makes the individual contributions of these activities difficult to distinguish in cells. To determine whether these activities rely on the same molecular features, we investigated the structure and biochemical properties of the SETX helicase domain (SETX-HD). Cryo-EM defines the conserved helicase core and RNA-binding channel of SETX-HD, while biochemical analyses identify distinct roles for two peripheral elements: a previously uncharacterised C-terminal RNA-binding domain (CT-RBD) and the prong subdomain. In reconstituted mammalian transcription assays, SETX-HD promotes passage beyond a defined RNAPII pause and elongation-complex dissociation on the same template. The CT-RBD stabilises RNA engagement and is required for efficient R-loop unwinding, whereas its deletion preserves both RNAPII-associated activities. Conversely, replacing the native SETX prong with its fungal Sen1 counterpart preserves R-loop unwinding but impairs both RNAPII outcomes. These reciprocal separation-of-function effects establish that a common SETX-HD RNA-translocation core depends on distinct peripheral elements for R-loop unwinding and action on mammalian RNAPII. This separation provides a direct route to determine how these activities contribute to SETX function in cells.

## INTRODUCTION

RNA polymerase II (RNAPII) pausing and backtracking are integral to transcriptional regulation.^1–2^ When prolonged, however, these states can obstruct advancing replisomes and, in some contexts, favour R-loop formation.^3–14^ R-loops comprise an RNA-DNA hybrid and displaced single-stranded DNA. Although R-loops participate in physiological processes, their inappropriate persistence can impede replication, alter chromatin organisation and gene expression, and promote genome instability.^15–27^ When an R-loop accompanies a stalled or arrested RNAPII elongation complex, removing the RNA-DNA hybrid does not necessarily restore productive transcription. The polymerase may resume elongation, undergo termination, or be removed through surveillance pathways.^2,28–35^ R-loop removal and resolution of RNAPII stalling or arrest are therefore related but mechanistically distinct tasks.^28,36^

Human senataxin (SETX) is a predominantly nuclear protein with a large N-terminal non-catalytic region and a conserved C-terminal helicase domain.^36^ Biallelic loss-of-function variants in SETX cause ataxia with oculomotor apraxia type 2, whereas dominant variants are associated with juvenile amyotrophic lateral sclerosis type 4.^37,38^ SETX contributes to both R-loop resolution^39–42^ and RNAPII regulation.^43^ SETX depletion causes R-loop accumulation at transcriptional pause and termination regions alongside defective RNAPII termination^39^ and increased RNAPII pausing and backtracking.^44^ These coupled effects make the respective contributions of R-loop resolution and RNAPII regulation to SETX function in cells difficult to distinguish.^28,36,44^ At the biochemical level, both full-length SETX and its isolated helicase domain (SETX-HD) act as RNA-stimulated 5′-to-3′ translocases that preferentially unwind 5′- overhang R-loop (5′-oh R-loop) substrates.^28,45^ SETX-HD also acts on mammalian RNAPII, promoting dissociation of stalled or arrested elongation complexes in a species-specific manner^28^ and opposing RNAPII backtracking at sequence-induced pauses to promote forward transcription.^44^ Whether R-loop unwinding and RNAPII action depend on the same features of SETX-HD, and what determines the distinct RNAPII outcomes of SETX activity, remain unclear.

The budding yeast orthologue of SETX, Sen1, provides a mechanistic reference for SETX. In the budding yeast *Saccharomyces cerevisiae*, Sen1 (*Sc*Sen1) acts in transcription termination^46–52^ and limits transcription-associated R-loops.^53–56^ SETX and Sen1 are superfamily 1B (SF1B) helicases with a conserved core composed of two RecA-like subdomains.^57,58^ The core translocates 5′-to-3′ along a single-stranded RNA that passes through the helicase channel and is termed the tracking strand.^59,60^ In the alternating-grip model, ATP-dependent rearrangements of the two subdomains redistribute contacts with the tracking strand to produce directional movement.^61,62^ Peripheral elements extend this conserved core, including the β-barrel and prong subdomains in Sen1.^59,63^ In a structurally characterised *Sc*Sen1-RNAPII pre-termination complex, the helicase core engages nascent RNA while the β-barrel contacts RNAPII directly.^64^ Yet, human SETX and fungal Sen1 differ in their broader domain architectures and recruitment pathways.^65^ No experimental structure of SETX-HD has been reported, leaving unclear how its nucleic-acid-binding channel relates to the tracking-strand architecture established in fungal Sen1 structures.^59^ The contributions of SETX-HD peripheral elements to efficient R-loop unwinding and RNAPII-associated activities have likewise remained undefined.

Here, cryo-EM and comparative sequence analysis define the architecture of human SETX-HD, revealing a conserved SF1B core with differences at positions corresponding to RNA contacts in fungal Sen1. Using reconstituted mammalian transcription assays, we show that SETX-HD can promote both passage beyond a defined RNAPII pause and elongation-complex dissociation on the same template. We find that efficient R-loop unwinding depends on a previously uncharacterised C-terminal RNA-binding domain (CT-RBD), yet SETX-HD variants lacking this domain retain both RNAPII activities. Conversely, replacing the native SETX prong with the corresponding *Sc*Sen1 prong preserves R-loop unwinding but impairs SETX-HD action on RNAPII. These reciprocal separation-of-function effects establish that distinct peripheral elements specialise a common SETX-HD RNA-translocation core for R-loop unwinding and RNAPII-associated activities.

## RESULTS

### Cryo-EM defines the nucleotide-bound architecture of human SETX-HD

To define the architecture of human SETX-HD, we analysed the recombinant helicase domain (residues 1674-2677; Fig. 1a) reconstituted with a synthetic 5′-overhang R-loop (5′-oh R-loop; Supplementary Fig. 1) by cryo-EM. Refinement and three-dimensional classification resolved two principal populations: one with additional nucleic-acid-associated density and a second without resolved nucleic acid but with ligand-like density in the interlobe cleft (Supplementary Fig. 2 and Supplementary Table S1). The latter population yielded a 3.83 Å consensus map. Focused refinement improved the density for the prong subdomain, and combination of the focused and consensus reconstructions produced a 4.0 Å composite map that supported an atomic model spanning residues 1695-2465 (Fig. 1b,c and Supplementary Fig. 3c-g).

**Fig. 1.**
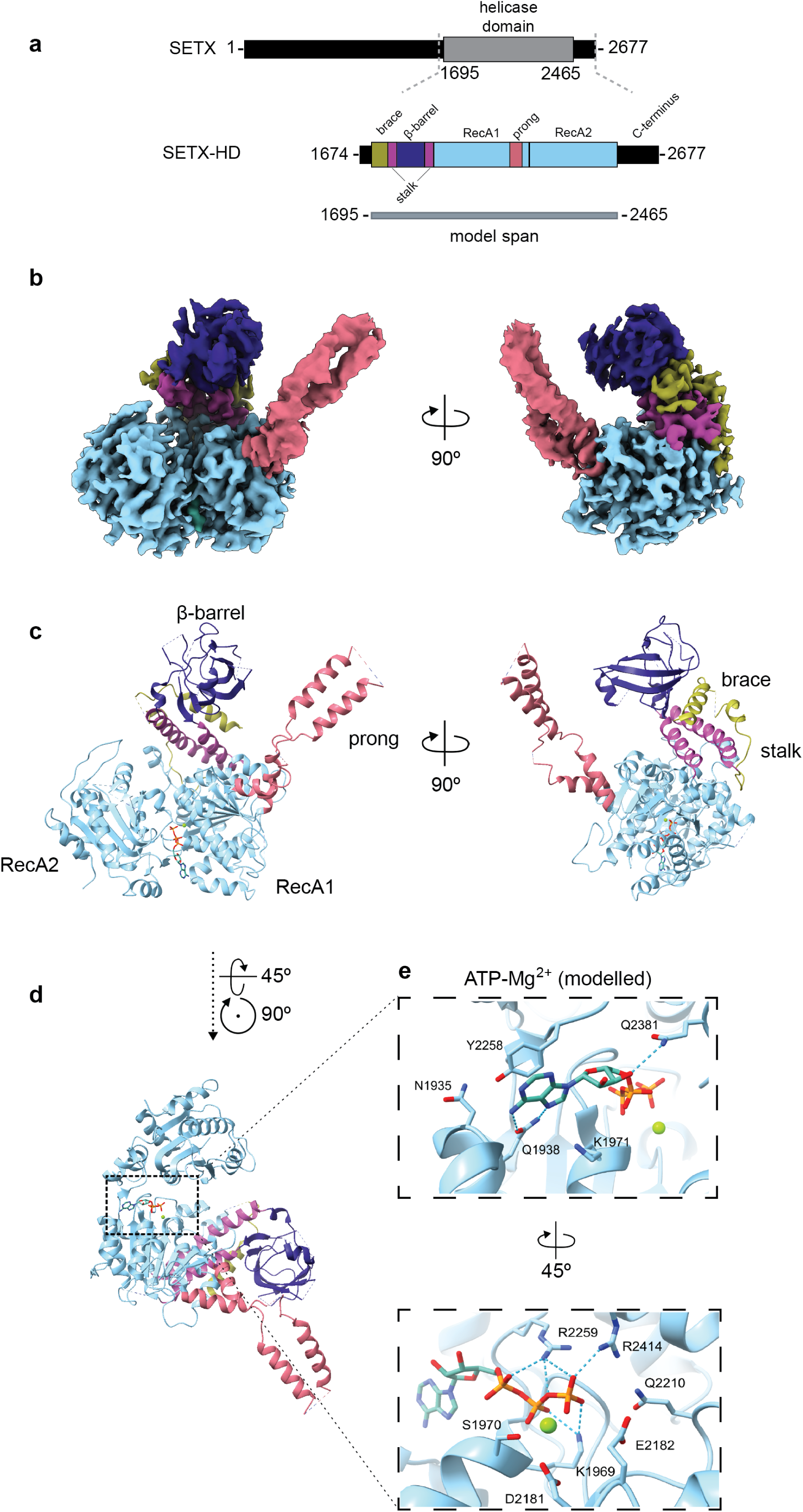
Cryo-electron microscopy defines the nucleotide-bound architecture of human SETX-HD. **a,** Domain organisation of full-length human senataxin (SETX), indicating the SETX-HD construct purified for cryo-EM (residues 1674-2677), the annotated helicase domain (residues 1695-2465), and the final atomic-model span (residues 1695-2465). The brace, β-barrel, stalk, RecA1, prong, RecA2, and C-terminal extension are indicated. **b,** The 4.0 Å composite cryo-electron microscopy (cryo-EM) map of the ATP-compatible nucleotide-bound SETX-HD state obtained in the presence of ATP and Mg²⁺, shown in two views related by 90° and coloured by structural element. **c,** Atomic model of SETX-HD (PDB 9TBG) shown in the same orientations as in b. Structural elements are coloured consistently across b-e and labelled in c. **d,** Atomic model with the interlobe nucleotide-binding cleft highlighted; the indicated rotations define the orientations used for the close views in e. **e,** Close views of the fitted ATP-Mg²⁺ model and surrounding active-site residues. The putative Mg²⁺ ion is shown as a green sphere and selected polar contacts as dashed lines. The ligand was modelled as ATP-Mg²⁺ on the basis of the sample conditions and compatible density.

The SETX-HD model adopts the characteristic SF1B organisation, with RecA1 and RecA2 forming the nucleotide-binding cleft and the β-barrel, stalk, brace, and prong arranged around the helicase core (Fig. 1a-c).^59,62,63^ The β-barrel is connected to RecA1 by two stalk helices and supported by the N-terminal brace, whereas the approximately 90-residue prong subdomain forms an insertion within RecA1 and is less well resolved than the helicase core. Structural alignment on RecA1 revealed that the SETX-HD β-barrel-stalk-brace assembly is rotated outwards by approximately 9° relative to *Sc*Sen1 and 12° relative to *Chaetomium thermophilum* Sen1 (*Ct*Sen1), while the prong subdomain also occupies a distinct position (Supplementary Fig. 5a,b).

Density in the interlobe cleft was compatible with a nucleotide and putative Mg²⁺ ion and was modelled as ATP-Mg²⁺ on the basis of the sample composition (Fig. 1d,e and Supplementary Fig. 6). In the fitted model, Y2258, Q1938, and Q2381 are positioned around the adenine and ribose, whereas the Walker A and Walker B motifs and adjacent conserved residues surround the phosphate and putative metal-binding region. Together, the density and sample composition identify this population as an ATP-compatible nucleotide-bound state obtained in the presence of ATP and Mg²⁺ (SETX-HD^ATP-Mg²⁺^).

The resulting map and model define the nucleotide-bound architecture of human SETX-HD and provide a structural reference for interpreting the nucleic-acid-associated particle population.

### SETX-HD engages nucleic acid via a conserved SF1B channel

The particle population containing additional nucleic-acid density yielded a 6.48 Å reconstruction (SETX-HD^R-loop^; Fig. 2a and Supplementary Fig. 3a,b). Positioning the experimentally determined SETX-HD model within this reconstruction located the additional density relative to the helicase architecture and nucleic-acid-binding channel (Fig. 2b).

**Fig. 2.**
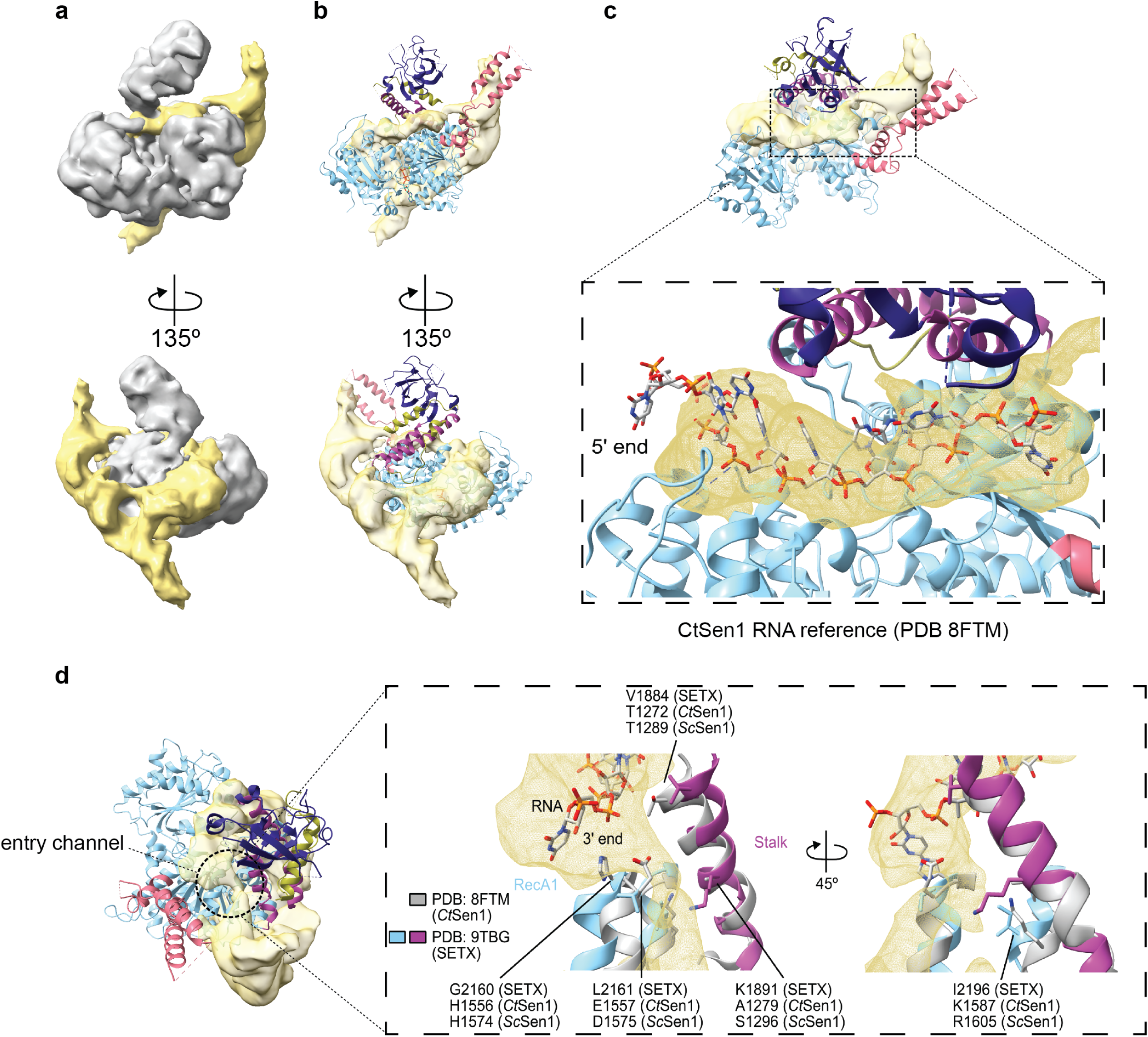
An R-loop-associated SETX-HD reconstruction suggests a candidate nucleic-acid path. **a,** The 6.48 Å 5′-oh R-loop-associated SETX-HD reconstruction shown in two views related by 135°. Protein density is shown in grey and additional density associated with the reconstituted nucleic-acid substrate in yellow. **b,** Atomic model of nucleotide-bound SETX- HD (PDB 9TBG) positioned in the 5′-oh R-loop-associated map and shown in the same orientations as in a. **c,** Close view of the additional density traversing the conserved nucleic-acid-binding channel. The RNA strand from RNA-bound *Chaetomium thermophilum* Sen1 (*Ct*Sen1; PDB 8FTM)^59^ was positioned by structural superposition and is shown as a reference for the tracking-strand path; its 5′ end is indicated. **d,** Entry-channel comparison of SETX-HD (PDB 9TBG) with RNA-bound *Ct*Sen1 (PDB 8FTM). SETX-HD is coloured by structural element and *Ct*Sen1 is shown in grey. Selected SETX residues and the corresponding positions in *Ct*Sen1 and *Saccharomyces cerevisiae* Sen1 (*Sc*Sen1) are labelled. The 3′ end of the *Ct*Sen1-bound RNA is indicated; the right-hand close view is related to the central view by a 45° rotation. The *Ct*Sen1-bound RNA is shown as a structural reference for comparing the tracking- strand path and entry-channel positions.

Superposition of RNA-bound *Ct*Sen1 on SETX-HD placed the central portion of the additional density along the conserved SF1B single-stranded nucleic-acid-binding channel (Fig. 2c).^59^ The additional density broadly followed the RNA path observed in *Ct*Sen1 through the central channel, while extending away from this trajectory towards the entry and exit sides. Outside the entry side, the density continued for several helical turns. Given the composition of the reconstituted sample and the continuity of this density with the channel-associated region, we use an RNA-DNA hybrid as the working interpretation of this region, while alternative duplex models remain indistinguishable in this reconstruction (Supplementary Fig. 7).

To compare the tracking-strand interaction network across the Sen1-SETX family, we compiled 30 distinct RNA-contact positions reported for RNA-bound *Ct*Sen1 and *Sc*Sen1. Twenty corresponding positions are represented in the SETX-HD model and could therefore be compared across SETX-HD, *Ct*Sen1, and *Sc*Sen1 using structural alignment (Supplementary Figs. 4 and 5c).^59,64^ Fifteen of these 20 positions retain the same residue in all three proteins, indicating broad residue-level conservation at positions corresponding to fungal tracking-strand contacts (Supplementary Fig. 5c). Among the five non-identical positions are two *Ct*Sen1 tracking-strand contacts with proposed mechanistic roles. R1427 contacts the tracking strand and has been proposed to act as a ratchet pawl that limits backward slippage during translocation, whereas H1556 contributes a polar contact within the distributed RecA1-side grip on the 3′ tracking strand.^59^ The corresponding positions in human SETX are occupied by P2048 and G2160, respectively (Supplementary Figs. 5c and 11a-d). Beyond these two positions, comparison of the entry-channel surfaces revealed further differences in local chemistry: several positions occupied by polar or charged residues in fungal Sen1 correspond to hydrophobic residues in SETX (V1884, L2161, and I2196), whereas an electropositive region containing K1891 is retained (Fig. 2d). Thus, SETX retains most residues at positions corresponding to fungal tracking-strand contacts while presenting a distinct local physicochemical environment at the entry side of the channel.

To test whether basic residues lining the SETX-HD channel contribute to RNA-stimulated ATP turnover, we analysed nine lysine- and arginine-to-alanine variants spanning the entry, hybrid-facing, and exit surfaces. All nine variants exhibited reduced RNA-stimulated ATPase activity despite near-wild-type bulk ssRNA binding (Supplementary Figs. 9 and 10). These mutational data, together with the structural comparison, define a largely conserved SETX-HD nucleic-acid channel with locally distinct entry chemistry and basic surfaces that support RNA-stimulated ATP turnover.

### A C-terminal RNA-binding domain stabilises SETX-HD on RNA and supports efficient R-loop unwinding

Although SETX retains most residues at positions corresponding to fungal tracking-strand contacts, selected substitutions and differences in entry-channel chemistry raised the question of whether additional SETX features contribute to productive RNA engagement. The SETX-HD^R-loop^ reconstruction also contained unassigned peripheral density near the C-terminal end of the model, providing an independent structural rationale for examining the unresolved C-terminal region (Fig. 3a).

**Fig. 3.**
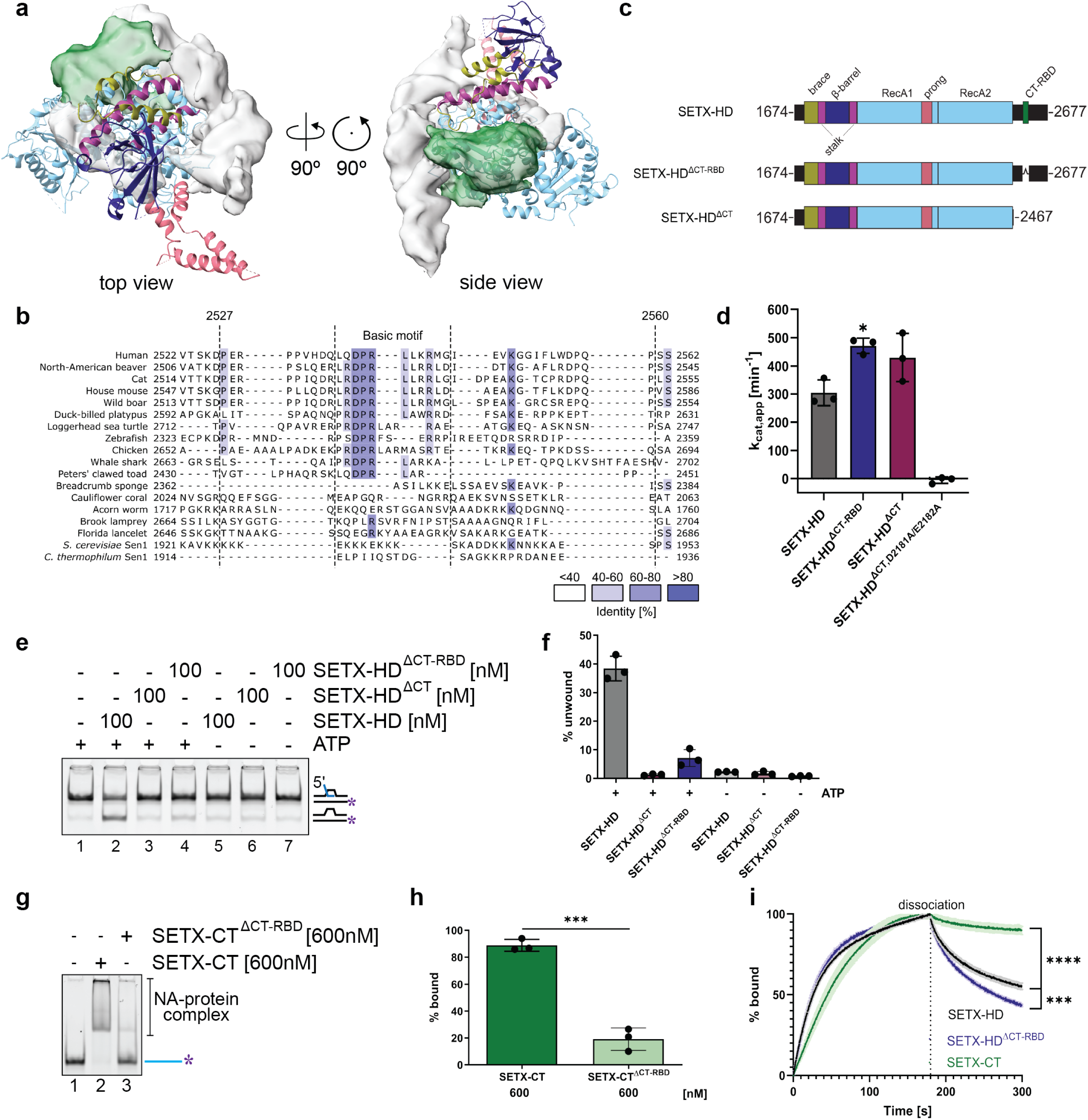
A C-terminal RNA-binding domain (CT-RBD) stabilises SETX-HD on RNA and supports efficient R-loop unwinding. **a,** Nucleotide-bound SETX-HD model positioned in the 5′-oh R-loop-associated cryo-EM map. Unassigned peripheral density near the expected emergence point of the unresolved C-terminal extension is coloured green; orthogonal top and side views are shown. **b,** Alignment of C-terminal sequences from representative SETX/Sen1 proteins. Human SETX residues 2527 and 2560 delimit the internal deletion, and the jawed-vertebrate-associated basic motif is indicated. Residues are shaded according to percentage identity. **c,** SETX-HD constructs used to test the C-terminal extension: SETX-HD (residues 1674-2677), SETX-HD^ΔCT-RBD^ (deletion of residues 2527-2560), and SETX-HD^ΔCT^ (residues 1674-2467). **d,** Apparent turnover numbers (*k*_cat,app_) for RNA-stimulated ATP turnover by the indicated proteins and the catalytically inactive SETX-HD^ΔCT,D2181A/E2182A^ control. **e,** Representative native polyacrylamide gel electrophoresis (PAGE) assay of 5′-oh R-loop unwinding by SETX-HD, SETX-HD^ΔCT^, and SETX-HD^ΔCT-RBD^ in the presence or absence of ATP. **f,** Quantification of 5′-oh R-loop unwinding in e. **g,** Representative electrophoretic mobility shift assay (EMSA) comparing binding of the isolated SETX C-terminal fragment (SETX-CT; residues 2455-2677) and SETX-CT^ΔCT-RBD^ to single-stranded RNA (ssRNA) at 600 nM protein. **h,** Quantification of ssRNA binding in g. **i,** Normalised biolayer interferometry (BLI) sensorgrams comparing association with and dissociation from immobilised 41-nt ssRNA for SETX-HD, SETX-HD^ΔCT-RBD^, and SETX-CT. SETX-HD and SETX-HD^ΔCT-RBD^ were analysed at 50 nM and SETX-CT at 2 μM. Data in d, f, and h are mean ± s.d. from three independent experiments. BLI measurements in i were performed in three independent experiments. Statistical comparisons were performed using two-sided Welch’s *t*-tests in d and two-sided unpaired Student’s *t*-tests in h and i. The *P* values for the indicated comparisons are: d, SETX-HD versus SETX-HD^ΔCT-RBD^, *P* = 0.0103; h, SETX-CT versus SETX-CT^ΔCT-RBD^ at 600 nM, *P* < 0.001; i, SETX-CT versus SETX-HD, *P* < 0.0001, and SETX-HD versus SETX-HD^ΔCT-RBD^, *P* < 0.001. BLI comparisons use the final recorded dissociation timepoint (299.803 s). Asterisks in the panels indicate these comparisons.

Comparative sequence analysis identified a basic motif within the distal C-terminal extension of SETX that was conserved across the sampled jawed vertebrates but was not recognisable in apparently full-length lamprey SETX proteins or in fungal Sen1 orthologues (Fig. 3b and Supplementary Figs. 12 and 13). Because incomplete or uncertain sampling in other earlier-branching lineages prevents more precise placement of its evolutionary origin, we describe the motif as a jawed-vertebrate-associated feature.

To test the contribution of this region to SETX-HD activity, we compared wild-type SETX-HD with deletion of the complete C-terminal extension (SETX-HD^ΔCT^; residues 1674-2467), deletion of the conserved basic segment (SETX-HD^Δ2527-2560^), and a catalytically inactive ΔCT control carrying the D2181A/E2182A substitutions (SETX-HD^ΔCT,D2181A/E2182A^; Fig. 3c and Supplementary Fig. 14a). SETX-HD^ΔCT^ retained RNA-stimulated ATP turnover comparable to wild-type SETX-HD, whereas SETX-HD^Δ2527-2560^ exhibited increased turnover. The catalytically inactive control lacked detectable ATP hydrolysis (Fig. 3d). By contrast, both C-terminal deletion variants exhibited near-background 5′-oh R-loop unwinding activity, whereas wild-type SETX-HD unwound approximately 40% of the substrate under the same conditions (Fig. 3e,f and Supplementary Fig. 14b-e). Negative-stain electron microscopy of SETX-HD^ΔCT^ yielded a reconstruction whose overall envelope resembled that of the low-pass-filtered nucleotide-bound SETX-HD cryo-EM map (Supplementary Fig. 15), arguing against gross misfolding. Thus, deletion of either the complete C-terminal extension or its basic segment preserved or increased RNA-stimulated ATP turnover while severely impairing 5′-oh R-loop unwinding.

We next characterised the nucleic-acid-binding properties of the C-terminal extension. In electrophoretic mobility-shift assays, the isolated C-terminal fragment SETX-CT (residues 2455-2677) bound ssRNAs of several lengths, as well as ssDNA and the 5′-oh R-loop (Supplementary Fig. 16a-h). Under these assay conditions, ssRNA binding by SETX-CT^Δ2527-2560^ was approximately fourfold lower than that of the wild-type fragment (Fig. 3g,h). Extending the deletion by a further 30 residues caused no additional decrease (Supplementary Fig. 16i-k), indicating that residues 2527-2560 account for most of the measurable ssRNA-binding contribution of the isolated C-terminal extension.

To assess how this region contributes to RNA engagement by SETX-HD, we examined RNA-binding kinetics by biolayer interferometry (BLI). Isolated SETX-CT exhibited slow apparent dissociation from ssRNA, whereas deletion of residues 2527-2560 from SETX-HD accelerated both apparent association with and dissociation from ssRNA (Fig. 3i and Supplementary Fig. 16l-n). By contrast, wild-type SETX-HD and SETX-HD^Δ2527-2560^ exhibited comparable association and dissociation kinetics with ssDNA (Supplementary Fig. 16o-v). Consistently, deletion of the basic segment from SETX-HD reduced binding to ssRNA and the 5′-oh R-loop, but not detectably to ssDNA, in electrophoretic mobility-shift assays (Supplementary Fig. 14h-l). Together, these measurements indicate that the C-terminal basic segment preferentially stabilises RNA engagement.

Purified SETX-CT did not restore SETX-HD^ΔCT^ unwinding *in trans* across the concentrations tested (Supplementary Figs. 14f,g and 16a), indicating that the RNA-binding activity of the isolated fragment is insufficient to restore efficient unwinding when supplied separately. These results identify residues 2527-2560 as a previously uncharacterised C-terminal RNA-binding domain (CT-RBD) that stabilises RNA engagement and supports efficient R-loop unwinding.

### SETX-HD promotes both pause escape and elongation-complex dissociation

We previously showed that SETX-HD promotes dissociation of mammalian RNAPII stalled at a G-rich tract by omission of GTP.^28^ At sequence-induced pauses, including a T-tract that strongly pauses RNAPII, Han *et al.*^44^ showed that SETX-HD opposes backtracking and promotes forward transcription. These observations raised the question of whether SETX promotes progression in one transcriptional context and dissociation in another, or whether both are alternative outcomes of SETX action on paused RNAPII. Because the assay used by Han *et al.* followed transcriptional progression rather than elongation-complex release, it did not distinguish between these possibilities. We therefore measured both pause escape and elongation-complex dissociation on the same T-tract-containing template.

We assembled mammalian RNAPII elongation complexes carrying fluorescent nascent RNA on a biotinylated DNA template containing a T-tract that produces a strong pause centred at T_3_-T_4_ and immobilised the template on streptavidin beads (Fig. 4a,b). RNA length reports how far RNAPII has transcribed, whereas separation of bead-associated and supernatant RNA distinguishes elongation complexes retained on the immobilised template from those that dissociate. Increasing SETX-HD concentrations progressively reduced the T_3_-T_4_ pause population and increased downstream transcripts, indicating that SETX-HD promotes passage beyond the T_3_-T_4_ pause (Supplementary Fig. 17a-c). If SETX-HD acted only by favouring continued elongation, increased passage beyond the pause would be expected to increase the run-off population. Run-off RNA did not increase detectably, however, while shorter upstream products accumulated (Supplementary Fig. 17c), raising the possibility that some elongation complexes instead dissociate before reaching the end of the templates. We therefore tested elongation-complex release directly by analysing bead-associated and supernatant RNA separately.

**Fig. 4.**
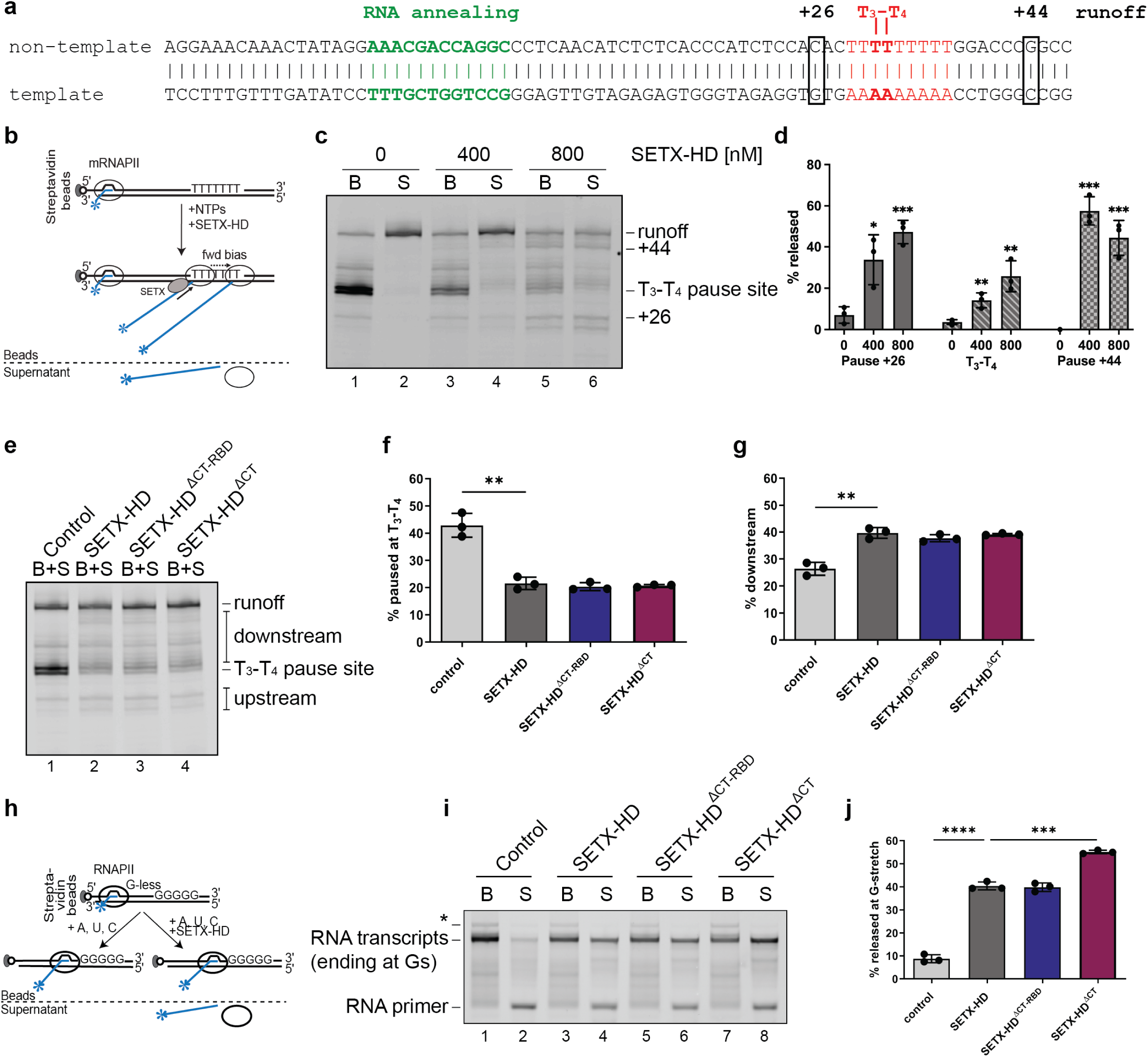
The C-terminal RNA-binding domain (CT-RBD) is not required for SETX-HD activity on RNAPII. **a,** Sequence and organisation of the immobilised elongation-complex substrate containing a defined T-rich pause. The RNA-annealing region, +26 position, principal T_3_-T_4_ pause, +44 position, and run-off position are indicated. **b,** Schematic of the immobilised T-rich pause assay used to resolve bead-associated and released RNA after addition of nucleoside triphosphates (NTPs) and SETX-HD. The fluorescent label on the nascent RNA is indicated by a blue asterisk. **c,** Representative denaturing PAGE analysis of bead-associated (B) and released supernatant (S) RNA after incubation with 0, 400, or 800 nM SETX-HD. Transcript positions corresponding to +26, the T_3_-T_4_ pause, +44, and run-off are indicated. **d,** Position-resolved quantification of RNA released into the supernatant for the +26, T_3_-T_4_, and +44 product classes in c. **e,** Representative denaturing PAGE analysis of combined bead and supernatant (B+S) RNA comparing buffer control, SETX-HD, SETX-HD^ΔCT-RBD^ (deletion of residues 2527-2560), and SETX-HD^ΔCT^. Upstream, T_3_-T_4_ pause, downstream, and run-off products are indicated. **f,g,** Quantification of the fraction of RNA at the T_3_-T_4_ pause (f) and downstream of the pause (g) in e. **h,** Schematic of the independent immobilised elongation-complex assay containing a G-less cassette followed by a G-rich tract. Transcription in the absence of GTP stalls RNAPII at the G-rich tract, and released labelled RNA is recovered in the supernatant. **i,** Representative denaturing PAGE analysis of bead-associated (B) and supernatant (S) RNA from the G-less/G-rich assay comparing buffer control, SETX-HD, SETX-HD^ΔCT-RBD^, and SETX-HD^ΔCT^. The asterisk marks transcripts produced by low-level read-through of the G-rich tract. **j,** Quantification of transcript release from the G-rich tract in i. Quantitative data in d, f, g, and j are mean ± s.d. from three independent experiments. Statistical comparisons used two-sided unpaired Student’s *t*-tests without adjustment for multiple comparisons. For comparisons marked by asterisks: d, relative to 0 nM SETX-HD, at 400 and 800 nM, respectively, +26, *P* = 0.0217 and *P* < 0.001; T_3_-T_4_, *P* = 0.00843 and *P* = 0.00725; +44, *P* < 0.001 at both concentrations; f, buffer versus SETX-HD, *P* = 0.00173; g, buffer versus SETX-HD, *P* = 0.00179; j, buffer versus SETX-HD, *P* < 0.0001, and SETX-HD versus SETX-HD^ΔCT^, *P* < 0.001. The T-rich pause assay reports endpoint RNA populations, whereas the G-less/G-rich assay directly reports transcript release from immobilised elongation complexes.

SETX-HD produced a concentration-dependent increase in RNA recovered in the supernatant, directly demonstrating elongation-complex dissociation (Fig. 4c,d). Release was not uniform across the template: the principal RNAPII population paused at T_3_-T_4_ was relatively resistant, whereas RNAPIIs paused at several upstream and downstream sequences were released more efficiently.

In summary, on the same pause-containing template, SETX-HD promotes both passage beyond the T_3_-T_4_ pause and elongation-complex dissociation, while susceptibility to release varies markedly with pausing sequence context.

### The CT-RBD is not required for SETX-HD activity on RNAPII

We next asked whether the CT-RBD that stabilises SETX-HD on RNA and supports efficient 5′-oh R-loop unwinding is likewise required for SETX-HD action on RNAPII. On the same T-tract-containing template, wild-type SETX-HD, SETX-HD^ΔCT-RBD^, and SETX-HD^ΔCT^ each reduced accumulation at the T_3_-T_4_ pause and increased downstream transcripts to similar extents (Fig. 4e-g), indicating that the C-terminal region is not required for SETX-HD-mediated RNAPII pause escape. The levels of run-off transcripts and the shorter upstream products were also similar among the SETX-HD variants (Supplementary Fig. 17d,e), suggesting no effect on RNAPII release.

To monitor the role of the CT-RBD on SETX-dependent elongation-complex dissociation more directly and accurately, we employed the G-less/G-rich template-based system used in our earlier work^28^, in which omission of GTP stalls RNAPII at a defined G-rich tract and the only measurable outcome of SETX action is elongation complex dissociation (Fig. 4h). Wild-type SETX-HD, SETX-HD^ΔCT-RBD^, and SETX-HD^ΔCT^ each promoted efficient transcript release (Fig. 4i,j), with SETX-HD^ΔCT^ producing a modest but significant increase in elongation complex dissociation.

Collectively, these results establish that, while the CT-RBD is critical for R-loop unwinding, it is dispensable for SETX-HD-mediated pause escape or dissociation of stalled RNAPII.

### Replacing the native SETX prong impairs SETX-HD action on RNAPII while preserving R-loop unwinding

The retention of both RNAPII-associated activities after removal of the CT-RBD led us to ask whether another divergent SETX-HD feature contributes to its action on mammalian RNAPII. Our previous cross-species reconstitutions established that human SETX-HD and fungal Sen1 act on RNAPII in a species-specific manner.^28^ Among the peripheral elements of SETX and Sen1, the prong is one of the least conserved (Supplementary Fig. 18).

Direct deletion of the SETX prong (residues 2061-2147; SETX-HD^Δprong^) produced insoluble protein (Supplementary Fig. 19a). We therefore tested prong interchangeability by replacing the native SETX prong with the corresponding *Sc*Sen1 prong, generating SETX-HD^Sen1prong^, which could be purified (Fig. 5a and Supplementary Fig. 19b). SETX-HD^Sen1prong^ retained 5′- oh R-loop binding and ATP-dependent unwinding (Fig. 5b-d and Supplementary Fig. 19c-e), indicating that the *Sc*Sen1 prong can functionally replace the native prong for R-loop unwinding.

**Fig. 5.**
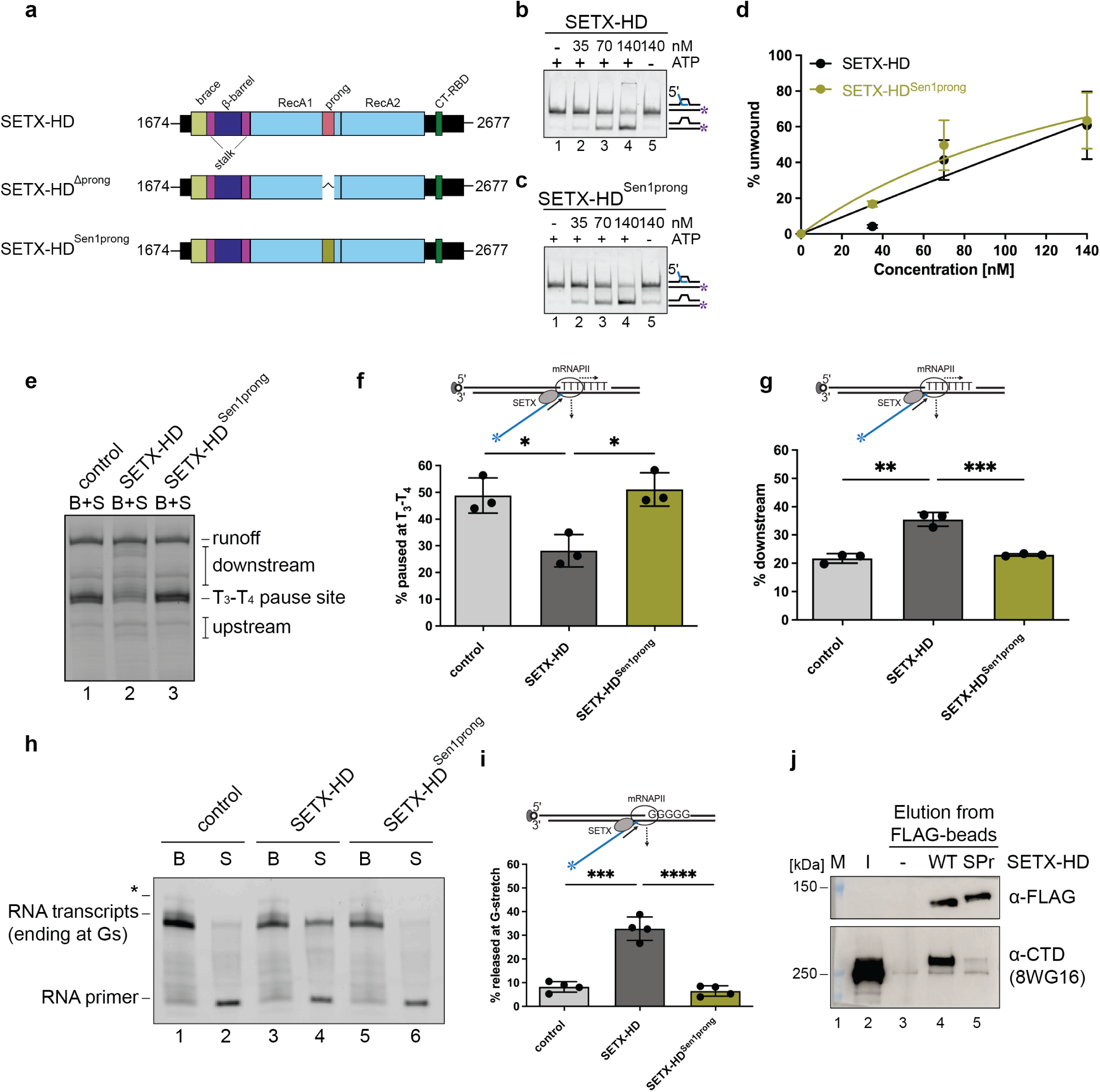
Replacing the native SETX prong impairs SETX-HD action on RNAPII while preserving R-loop unwinding. **a,** SETX-HD constructs used t test the prong: wild-type SETX-HD, SETX-HD^Δprong^ (deletion of residues 2061-2147), and SETX-HD^Sen1prong^, in which the native SETX prong is replaced by the corresponding *Sc*Sen1 prong. **b,c,** Representative native PAGE assays of 5′-oh R-loop unwinding by increasing concentrations of SETX-HD (b) or SETX-HD^Sen1prong^ (c) in the presence of ATP, with a 140 nM no-ATP control. **d,** Quantification of 5′-oh R-loop unwinding in b and c. **e,** Representative denaturing PAGE analysis of combined bead and supernatant (B+S) RNA from the T-rich pause assay comparing buffer control, SETX-HD, and SETX-HD^Sen1prong^. **f,g,** Quantification of the fraction of RNA at the T_3_-T_4_ pause (f) and downstream of the pause (g) in e. **h,** Representative denaturing PAGE analysis of bead-associated (B) and released supernatant (S) RNA from the independently stalled G-less/G-rich elongation complex comparing buffer control, SETX-HD, and SETX-HD^Sen1prong^. The asterisk marks transcripts produced by low-level read-through of the G-rich tract. **i,** Quantification of transcript release from the G-rich tract in h. **j,** Pull-down of RNAPII from clarified HEK293T cell lysate using FLAG-tagged SETX-HD or SETX-HD^Sen1prong^, followed by capture on anti- FLAG resin. Wild-type SETX-HD pulled down RNAPII, whereas SETX-HD^Sen1prong^ did not pull down detectable RNAPII above the no-protein background. M, molecular-mass marker; I, input; -, no-protein control; WT, SETX-HD; SPr, SETX-HD^Sen1prong^. SETX-HD proteins were detected with anti-FLAG antibody and RNAPII with antibody 8WG16 against the RPB1 C-terminal domain. Data in d, f, and g are mean ± s.d. from three independent experiments; data in i are mean ± s.d. from four independent experiments. Statistical comparisons used two-sided unpaired Student’s *t*-tests without adjustment for multiple comparisons. The *P* values for the indicated comparisons are: f, buffer versus SETX-HD, *P* = 0.0160, and SETX-HD versus SETX-HD^Sen1prong^, *P* = 0.0101; g, buffer versus SETX-HD, *P* = 0.00134, and SETX-HD versus SETX- HD^Sen1prong^, *P* < 0.001; i, buffer versus SETX-HD, *P* < 0.001, and SETX-HD versus SETX- HD^Sen1prong^, *P* < 0.0001.

Prong replacement, however, abolished SETX-HD activities on mammalian RNAPII. On the T-tract-containing template, SETX-HD^Sen1prong^ failed to promote pause escape, as the T_3_-T_4_ pause and downstream transcript populations remained indistinguishable from the no-protein control (Fig. 5e-g and Supplementary Fig. 19f,g). In the G-less/G-rich assay, SETX-HD^Sen1prong^ was likewise unable to induce elongation-complex dissociation (Fig. 5h,i). The *Sc*Sen1 prong can therefore support R-loop unwinding within SETX-HD but does not efficiently substitute for the native SETX prong when SETX-HD acts on mammalian RNAPII.

We hypothesised that the native SETX prong contributes to SETX-HD association with mammalian RNAPII and tested this possibility using pull-down assays with clarified HEK293T cell extracts. Wild-type SETX-HD recovered RNAPII, whereas SETX-HD^Sen1prong^ did not recover detectable RNAPII above the no-protein background (Fig. 5j). The loss of detectable RNAPII association provides a possible molecular explanation for the impaired activity of the prong chimera on reconstituted elongation complexes.

The contrasting effects of CT-RBD deletion and prong replacement establish distinct peripheral requirements for R-loop unwinding and SETX-HD action on mammalian RNAPII (Fig. 6).

**Fig. 6.**
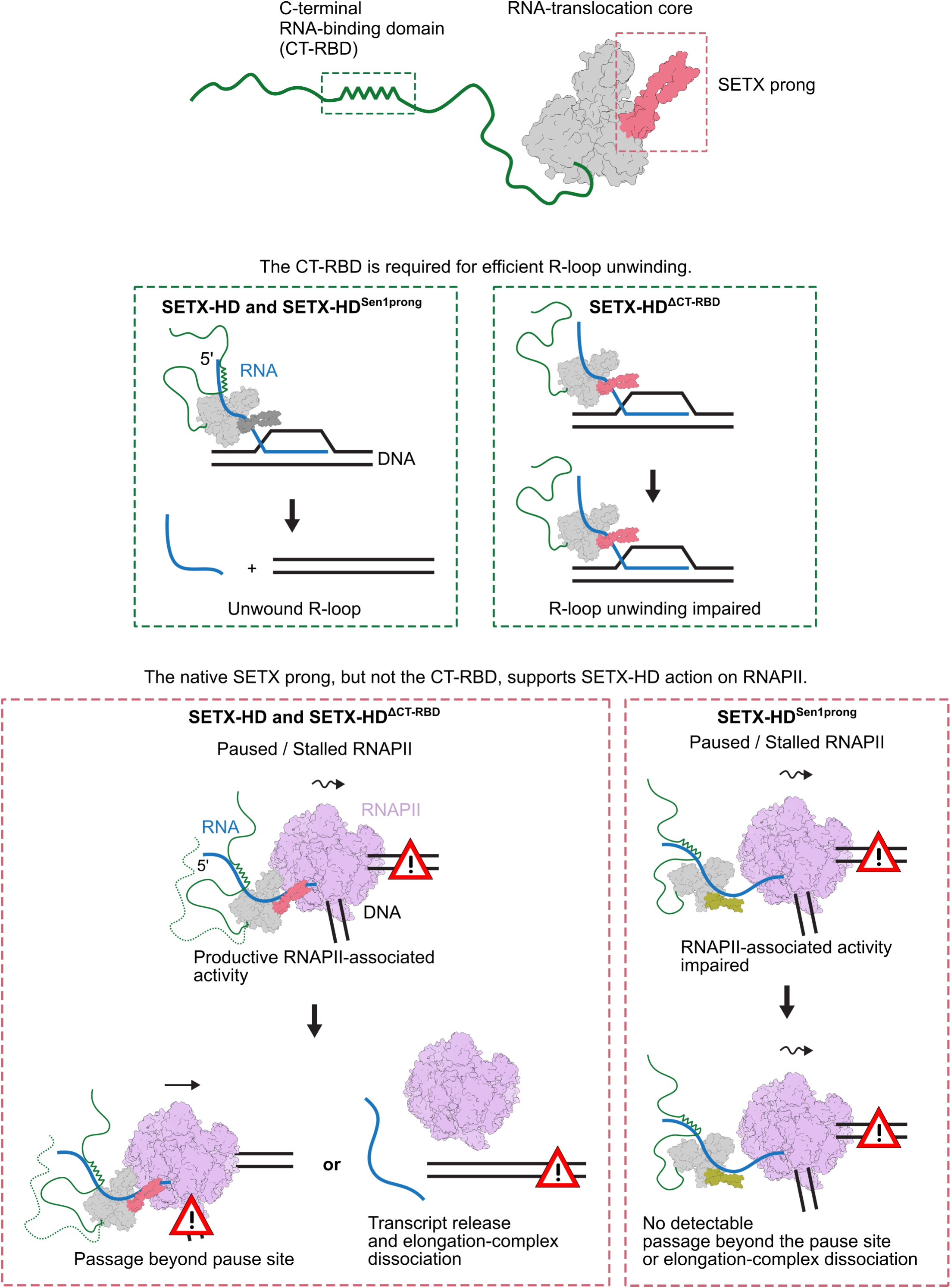
Distinct peripheral elements support R-loop unwinding and SETX-HD action on RNAPII. Schematic summary of the reciprocal separation-of-function effects of the C-terminal RNA-binding domain (CT-RBD) and native SETX prong. The upper schematic depicts the RNA-translocation core and its peripheral elements. The C-terminal extension, including the CT- RBD, is represented schematically. RNA is shown in blue, DNA in black, RNAPII in purple, the native SETX prong in pink, the substituted ScSen1 prong in ochre, and the C-terminal region containing the CT-RBD in green. In the RNAPII panel, solid and dotted green lines schematically distinguish the intact C-terminal extension of SETX-HD from the corresponding CT-RBD-deleted extension of SETX-HD^ΔCT-RBD^, respectively. Red warning triangles indicate positions on the DNA template at which RNAPII may pause or stall. In the R-loop context, SETX-HD and SETX-HD^Sen1prong^ support efficient R-loop unwinding, whereas SETX-HD^ΔCT-RBD^ is strongly impaired, identifying the CT-RBD as a requirement for efficient R-loop unwinding. In the RNAPII context, SETX-HD and SETX-HD^ΔCT-RBD^ retain passage beyond a defined pause and elongation-complex dissociation, whereas SETX-HD^Sen1prong^ is strongly impaired in both activities while retaining R-loop unwinding. These reciprocal effects separate R-loop unwinding from SETX-HD action on mammalian RNAPII and assign different peripheral requirements to the two functional contexts.

## DISCUSSION

Our findings establish that R-loop unwinding and action on mammalian RNAPII are separable within SETX-HD. The C-terminal RNA-binding domain (CT-RBD) stabilises RNA engagement required for efficient R-loop unwinding, whereas the native SETX prong subdomain is crucial for SETX-HD action on mammalian RNAPII. A common SETX-HD RNA-translocation core can therefore support both activities while relying on different peripheral elements.

Directional translocation by SF1 helicases couples ATP-dependent rearrangements of the two RecA-like lobes to an alternating grip on the single-stranded tracking strand.^61,62^ Nucleotide binding and hydrolysis change the relative positions of RecA1 and RecA2, while their interactions with the tracking strand alternate between tighter and weaker states, converting reciprocal lobe movements into forward translocation.^61–62^ SETX preserves most residues at positions corresponding to fungal tracking-strand contacts but differs at positions corresponding to *Ct*Sen1 R1427 and H1556, implicated in suppressing backward slippage and maintaining the RecA1-side grip, respectively (Supplementary Fig. 5c).^59^ Our SETX-HD structure, obtained in the presence of ATP and Mg²⁺, captures a relatively closed RecA-lobe arrangement and provides the first structural view of a Sen1-SETX family member under ATP/Mg²⁺ conditions. Together with structurally characterised fungal Sen1 states, it extends the available conformational series across the open and closed RecA-lobe arrangements expected during ATP-dependent alternating-grip translocation (Supplementary Fig. 20).^59,61–64^ These observations support a model in which SETX-HD retains the Sen1-like mechanism of directional RNA translocation while redistributing tracking-strand grip across the channel. We therefore hypothesised that this redistribution could increase dependence on additional RNA-binding elements outside the conserved ATPase core that stabilise the tracking strand during translocation.

Within SETX-HD, the CT-RBD provides such an additional RNA-binding element. Deletion of the C-terminal basic segment destabilises RNA engagement and severely impairs R-loop unwinding despite retained RNA-stimulated ATP turnover, indicating that ATP hydrolysis alone is insufficient for efficient strand displacement when this additional grip is lost. Its inferred proximity to the exit side of the candidate tracking-strand path suggests how the CT-RBD could support this coupling. We propose that the CT-RBD contributes an exit-side grip on the 5′ RNA after it has passed through the nucleic-acid-binding channel, extending tracking-strand engagement beyond the conserved helicase core. Such an additional contact could reduce disengagement or backward slippage as the RecA lobes alternate their grip during successive ATPase cycles, thereby coupling ATP-dependent RNA translocation to efficient R-loop unwinding. The failure of isolated SETX-CT to restore SETX-HD^ΔCT^ unwinding *in trans* further suggests that this function depends on the spatial relationship between the C-terminal grip and the channel rather than on RNA binding alone.

The evolutionary distribution of these features suggests substantial plasticity in how the Sen1-SETX family maintains tracking-strand engagement. In the sampled lamprey SETX proteins, the positions corresponding to *Ct*Sen1 R1427 and H1556 are occupied by different residues, whereas the C-terminal basic motif associated with the CT-RBD in jawed vertebrates is not recognisable (Supplementary Figs. 11e,f and 13). Divergence at these prominent fungal tracking-strand positions can therefore coexist with the absence of the CT-RBD-associated motif, arguing against a simple model in which this peripheral RNA-binding feature directly replaces those individual contacts. We instead propose that the evolution of SETX involved a redistribution of tracking-strand engagement between a broadly conserved central interaction network and lineage-specific contributions from peripheral elements. In jawed vertebrates, the CT-RBD may represent one such solution, providing the proposed exit-side grip and reducing dependence on the precise distribution of contacts within the central channel. The motif-lacking lamprey configuration raises the possibility that productive RNA engagement can be maintained through a different balance of the remaining channel contacts or through other, as yet unidentified, RNA-binding elements.

SETX-HD variants lacking the CT-RBD remain active on RNAPII, suggesting that the elongation complex provides a different context for stable engagement of the helicase. On an isolated 5′-oh R-loop, the CT-RBD stabilises SETX-HD on the tracking RNA and supports efficient unwinding. During transcription, by contrast, the nascent RNA is already constrained by RNAPII, and SETX can additionally associate with the polymerase. Previous evidence for SETX-RNAPII association^43^, together with the *Sc*Sen1-RNAPII structure, in which the Sen1 β-barrel contacts RNAPII while its prong contributes to nascent RNA engagement,^64^ supports such a mode of interaction. Our earlier finding that human SETX-HD and fungal Sen1 act on RNAPII in a species-specific manner^28^ further indicates that the polymerase context imposes species-specific requirements on helicase action. Differences at the β-barrel-RPB3 interface have been proposed to contribute to this specificity.^64^ Our present results extend that model: replacing the native SETX prong with its *Sc*Sen1 counterpart compromises RNAPII association and activity despite retention of the human SETX β-barrel. We propose that the native SETX prong contributes to productive engagement with the mammalian elongation complex through interactions with RNAPII, nascent RNA, or both. Such prong-dependent engagement could help explain how peripheral elements confer species-specific requirements on a conserved RNA-translocation motor.

SETX-HD promotes both passage beyond a defined RNAPII pause and elongation-complex dissociation on the same transcription template, reconciling our earlier finding of SETX-mediated release with subsequent evidence that SETX opposes backtracking and promotes forward transcription.^28,44^ SETX-HD behaviour parallels that of fungal Sen1, which can promote both transcriptional progression and dissociation of RNAPII and RNAPIII elongation complexes,^60,66^ suggesting that SETX has retained a conserved Sen1-like repertoire of transcription-associated activities. In our system, susceptibility to SETX-HD-mediated dissociation varies markedly with RNAPII position: complexes at the principal T_3_-T_4_ pause are comparatively resistant, whereas those paused at upstream and downstream positions are released more efficiently. This pattern implicates local sequence context and associated elongation-complex properties, potentially including RNA-DNA hybrid stability and backtracking propensity, rather than transcript length alone or a simple distinction between paused and stalled RNAPII. In the fungal Sen1-RNAPII system, ATP-dependent translocation along nascent RNA can promote forward movement of the polymerase and is proposed to drive a termination-prone hypertranslocated state.^60,64^ We propose that SETX retains this mechanistic principle, with local elongation-complex properties influencing whether helicase-driven RNAPII movement favours continued transcription or elongation-complex dissociation.

Across helicase and nucleic-acid-remodelling families, conserved ATPase cores are commonly specialised by accessory elements that influence substrate recognition, regulation, and ATP-dependent processing.^57,67–71^ The complementary separation-of-function relationships identified here extend this principle: distinct peripheral elements can enable a common RNA-translocation core to perform distinct biochemical tasks. SETX therefore illustrates a form of modularity in which peripheral specialisation acts not only at the level of substrate recognition, but also in determining how ATP-dependent RNA translocation is converted into a particular molecular outcome.

The activities defined with SETX-HD operate within a much larger protein whose additional elements and interaction partners can influence recruitment to R-loops and RNAPII and the efficiency with which these substrates are processed. Full-length *Ct*Sen1 provides a structural precedent for such regulation, because its N-terminal region can restrict access to the nucleic-acid-binding cleft of the helicase domain.^59^ Human SETX appears to use its surrounding regions differently: purified full-length SETX can unwind R-loop substrates that isolated SETX-HD unwinds poorly and remains responsive to stimulation by BRCA1-BARD1.^28,45^ Full-length SETX can also reach RNAPII and R-loop-rich transcription regions through interactions outside SETX-HD, including pathways involving SMN and BRCA1-BARD1.^72–74^ Because SETX is recruited to both RNAPII and R-loop-rich transcriptional regions, cellular SETX perturbation produces coupled defects in R-loop homeostasis and RNAPII behaviour, making the specific contribution of SETX to R-loop resolution and RNAPII regulation difficult to distinguish.^28,36,39,44^ The separation-of-function variants identified here provide a direct experimental route to disentangle these activities in cells and determine how each contributes to genome stability during transcription, and when these functions intersect.

## METHODS

### Plasmid construction

Initial plasmid assembly was performed using ligation-independent cloning (LIC)^75^; point mutants and internal-deletion constructs were generated as described below.

Codon-optimised DNA fragments encoding the helicase domain of wild-type human SETX (SETX-HD; residues 1674-2677) or the truncated SETX-HD^ΔCT^ variant (residues 1674-2467) were used to generate tagged constructs. SETX-HD was first inserted into plasmid 2L-T (pET Flag TEV LIC cloning vector; Addgene #29715), producing an N-terminally FLAG-tagged construct. This construct was subcloned into 2Bc-T (pET His_6_ LIC cloning vector; Addgene #37236) to generate FLAG-SETX-HD-(His)_6_ and then into plasmid 438-A (pFastBac cloning vector with BioBrick PolyPromoter LIC Subcloning; Addgene #55218) for insect-cell expression. SETX-HD^ΔCT^ was inserted into 2Bc-T to generate C-terminally tagged SETX-HD^ΔCT^-(His)_6_ and then subcloned into 438-A for insect-cell expression. The D2181A/E2182A, R1899A, R1904A, K2131A, K2129A/K2131A, R2136A, K2139A, R2168A, R2198A, and R2235A substitutions were introduced by site-directed mutagenesis. SETX-HD^Δ2061-2147^ (SETX-HD^Δprong^) and SETX-HD^Δ2527-2560^ (SETX-HD^ΔCT-RBD^) were generated by divergent PCR using 438-A-FLAG-SETX-HD-(His)_6_ as the template. To generate SETX-HD^Sen1prong^, the SETX prong-encoding sequence was deleted by divergent PCR and replaced by ligation of a PCR fragment encoding the *S. cerevisiae* Sen1 prong subdomain (residues 1461-1554).

For bacterial expression of the SETX C-terminal extension (SETX-CT; residues 2455-2677), the corresponding coding sequence was inserted into plasmid 2B-T (pET His_6_ TEV LIC cloning vector; Addgene #29666), producing an N-terminally (His)_6_-tagged construct. The SETX-CT^Δ2527-2560^ (SETX-CT^ΔCT-RBD^) and SETX-CT^Δ2527-2590^ internal-deletion variants were generated by divergent PCR.

Oligonucleotides used in this study are listed in Supplementary Table S2. Plasmids and constructs used in this study are listed in Supplementary Tables S3 and S4, respectively. The sequence integrity of all constructs was verified by sequencing, and the identities of the purified proteins were confirmed by mass spectrometry.

### Protein expression in insect cells

Recombinant bacmids were generated by transposing the coding and regulatory sequences from the constructs described above in *E. coli* DH10Bac cells (Thermo Fisher Scientific). *Sf*9 cells were transfected with the recombinant bacmids using FuGENE Transfection Reagent (Promega), and the resulting virus was amplified to generate P1 stocks. For protein expression, 600 mL cultures of High Five cells (Thermo Fisher Scientific) at 1 × 10^6^ cells/mL were infected with the corresponding P1 virus at a multiplicity of infection (MOI) >1. Cells were harvested 48 h after infection, washed with 1× PBS, and stored at -80 °C.

### Protein purification

#### Purification of SETX-HD and its variants used for in vitro analyses

Thawed insect-cell pellets expressing SETX-HD variants were resuspended in lysis buffer containing 50 mM Tris-Cl, pH 8, 500 mM NaCl, 10% (v/v) glycerol, 1 mM DTT, 0.4% (v/v) Triton X-100, 10 mM imidazole, and 2 mM MgCl₂, supplemented with protease inhibitors (0.66 μg/mL pepstatin, 5 μg/mL benzamidine, 4.75 μg/mL leupeptin, and 2 μg/mL aprotinin) and Benzonase (25 U per mL of lysate). The clarified lysate was applied to 2 mL Ni-NTA resin (Qiagen) pre-equilibrated in wash buffer (50 mM Tris-Cl, pH 8, 500 mM NaCl, 10 mM imidazole, 2 mM MgCl₂, and 1 mM DTT). The beads were subsequently washed extensively with the wash buffer. Bound SETX-HD was eluted in 50 mM Tris-Cl, pH 8, 350 mM NaCl, 2 mM MgCl₂, 1 mM DTT, and 600 mM imidazole. SETX-HD-containing fractions were diluted twofold with 50 mM Tris-Cl, pH 8 and 1 mM DTT and applied to a 1 mL HiTrap Heparin HP column (Cytiva). Protein was eluted in 50 mM Tris-Cl, pH 7.5, 2 mM MgCl₂, and 1 mM DTT using a 35-column-volume linear gradient from 200 to 650 mM NaCl. Fractions containing homogeneous SETX-HD were pooled and concentrated. Glycerol was adjusted to 10% (v/v), and samples were snap-frozen in liquid nitrogen and stored at -80 °C.

#### Purification of SETX-HD for structural analysis by cryogenic electron microscopy (cryo-EM) and its reconstitution with a 5′-oh R-loop

SETX-HD was purified from thawed insect-cell pellets through the Ni-NTA elution step as described above. SETX-HD-containing fractions were pooled and incubated with TEV protease to remove the C-terminal His_6_ and N-terminal FLAG tags. The sample was concentrated and loaded onto a 24 mL Superdex 200 Increase 10/300 (Cytiva) size-exclusion column equilibrated in 25 mM Tris-Cl, pH 7.5, 200 mM NaCl, 1 mM DTT, 2 mM MgCl₂, and 1 mM ATP. Protein was eluted isocratically. Fractions containing homogeneous SETX-HD were pooled and concentrated. Glycerol was adjusted to 10% (v/v), and samples were snap-frozen in liquid nitrogen and stored at -80 °C.

Thawed SETX-HD (17 μM final concentration) was mixed with 120 pmol annealed 5′-oh R-loop substrate (1.5 μM final concentration) in a total volume of 80 μL. After incubation on ice for 20 min, the sample was centrifuged to remove aggregates and loaded onto either a 3 mL Superdex 200 Increase 5/150 or 3 mL Superose 6 size-exclusion column connected to an ÄKTA pure Micro system (Cytiva). The column was equilibrated in 25 mM Tris-Cl, pH 7.5, 150 mM NaCl, 1 mM DTT, 1 mM ATP, 1.5 mM MgCl₂, and 0.5% (v/v) glycerol. Complexes were eluted isocratically, and complex-containing fractions were identified by monitoring the Cy3 label on the 5′-oh R-loop non-template strand. Because the amount of recovered material was limited, the concentration of the post-SEC sample used for grid preparation was not determined.

#### Purification of wild-type SETX-HD and the R1899A, R1904A, K2131A, K2129A/K2131A, R2136A, K2139A, R2168A, R2198A, and R2235A variants

A thawed insect-cell pellet corresponding to 200 mL of culture was resuspended in lysis buffer containing 50 mM Tris-Cl, pH 8, 500 mM NaCl, 10% (v/v) glycerol, 2 mM MgCl₂, 1 mM DTT, 0.4% (v/v) Triton X-100, and 10 mM imidazole, supplemented with protease inhibitors (0.66 μg/mL pepstatin, 5 μg/mL benzamidine, 4.75 μg/mL leupeptin, and 2 μg/mL aprotinin) and Benzonase (25 U per mL of lysate). The clarified lysate was applied to 250 μL Ni-NTA resin (Qiagen) pre-equilibrated in wash buffer (50 mM Tris-Cl, pH 8, 500 mM NaCl, 10 mM imidazole, 2 mM MgCl₂, and 1 mM DTT). The beads were subsequently washed extensively with the wash buffer. SETX-HD variants were eluted in 50 mM Tris-Cl, pH 8, 350 mM NaCl, 2 mM MgCl₂, 1 mM DTT, and 600 mM imidazole. The buffer was exchanged for storage buffer containing 25 mM Tris-Cl, pH 7.5, 300 mM NaCl, 2 mM MgCl₂, and 1 mM DTT. Proteins were concentrated, glycerol was adjusted to 10% (v/v), and samples were snap-frozen in liquid nitrogen and stored at -80 °C.

#### Purification of SETX-CT, SETX-CT^ΔCT-RBD^, and SETX-CT^Δ2527-2590^

A 5 g pellet of *E. coli* BL21 cells expressing SETX-CT variants was resuspended in lysis buffer containing 50 mM Tris-Cl, pH 8, 500 mM NaCl, 10 mM imidazole, and 1 mM DTT, supplemented with protease inhibitors (0.66 μg/mL pepstatin, 5 μg/mL benzamidine, 4.75 μg/mL leupeptin, and 2 μg/mL aprotinin). Cells were lysed by sonication, and the clarified lysate was applied to 250 μL Ni-NTA resin (Qiagen) pre-equilibrated in wash buffer (50 mM Tris-Cl, pH 8, 500 mM NaCl, 10 mM imidazole, 2 mM MgCl₂, and 1 mM DTT). The beads were subsequently washed extensively with the wash buffer. SETX-CT variants were eluted in 50 mM Tris-Cl, pH 8, 350 mM NaCl, 1 mM DTT, and 600 mM imidazole. SETX-CT- containing fractions were pooled, concentrated, and loaded onto a 24 mL Superdex 75 Increase 10/300 (Cytiva) size-exclusion column equilibrated in 25 mM Tris-Cl, pH 7.5, 200 mM NaCl, and 1 mM DTT. Protein was eluted isocratically. Fractions containing homogeneous SETX-CT were pooled and concentrated. Glycerol was adjusted to 10% (v/v), and samples were snap-frozen in liquid nitrogen and stored at -80 °C.

### Purification of RNAPII

The mammalian 12-subunit RNAPII complex was purified from porcine thymus. Chromatin-bound RNAPII was purified to homogeneity by sequential ion-exchange, immunoaffinity, ion-exchange, and size-exclusion chromatography using a procedure adapted from Bernecky *et al.*^76^

### Synthetic nucleic acid substrates

HPLC-purified oligonucleotides used to assemble fluorescent RNA-DNA substrates were purchased from Sigma; their sequences and fluorophore modifications are listed in Supplementary Table S2. ssRNA sequences were designed using INFO-RNA Sequence Designer^77–79^ to minimise predicted secondary structure. Duplex substrates were assembled at a 1:2 molar ratio of labelled strand to unlabelled complementary strand. Three-stranded substrates were assembled by combining the labelled strand with a fivefold molar excess of each unlabelled strand in annealing buffer (25 mM Tris-Cl, pH 7.5, 100 mM NaCl, and 3 mM MgCl₂). Samples were heated at 75 °C for 5 min and cooled to room temperature. Annealed substrates were purified by native polyacrylamide gel electrophoresis.

### ATPase assays

ATP hydrolysis was quantified using the PiColorLock kit (Abcam). Each 220 μL reaction contained 50 mM Tris-Cl, pH 7.5, 50 mM NaCl, 0.3 mg/mL BSA, 2.5 mM ATP, 0.5 mM MgCl₂, 1.2 μg MS2 phage ssRNA (Merck), and SETX-HD diluted in 50 mM Tris-Cl, pH 7.5, 300 mM NaCl, 1 mM DTT, and 0.01% NP-40. For the C-terminal-variant series in Fig. 3d, final protein concentrations were 1.875, 3.75, and 7.5 nM; for the nucleic-acid-channel mutant series in Supplementary Fig. 9d, final protein concentrations were 3.75, 7.5, and 15 nM. Reactions were initiated by incubation at 37 °C. At 0, 2.5, 5, and 10 min, 50 μL aliquots were transferred to the detection mixture, which had been prepared in a 96-well deep-well plate according to the manufacturer’s instructions. After 5 min, the colourimetric reaction was stopped by adding the supplied stabilising reagent, and absorbance at 650 nm was measured using a Tecan Infinite F500 plate reader. The conversion of absorbance to free-phosphate concentration (C_Pi_) was performed using a calibration curve generated with the supplied Pi standard. For each sample, the signal was corrected by subtracting the t = 0 value and the corresponding no-protein-control signal. C_Pi_ was plotted against time and fitted by linear regression; the slope represented the initial velocity (v_0_). v_0_ was then plotted against protein concentration and fitted by linear regression; the slope represented the apparent turnover number (*k*_cat,app_) at the fixed substrate concentrations used. Mean *k*_cat,app_ values from three independent experiments were compared between protein variants.

### Electrophoretic mobility shift assay (EMSA)

SETX-HD and its C-terminal deletion variants were analysed at 7.5, 15, 30, and 60 nM; SETX-HD and SETX-HD^Sen1prong^ were analysed at 7.5, 15, 30, and 60 nM; and isolated SETX-CT variants were analysed at 75, 150, 300, and 600 nM. Proteins were incubated with 10 nM fluorescently labelled nucleic acid substrate in buffer D (25 mM Tris-Cl, pH 7.5, 1 mM DTT, 0.1 mg/mL BSA, 5 mM MgCl₂, and 100 mM NaCl) for 20 min on ice. For RNA-containing substrates, RNasin Ribonuclease Inhibitor (Promega) was included at 2 U/μL. After addition of loading buffer (60% (v/v) glycerol and 0.005% Orange G), samples were resolved on 7.5% (w/v) native polyacrylamide gels in 0.5× TBE at 75 V for 1 h. Fluorescent species were imaged using a Typhoon FLA-9000 Starion scanner and quantified using MultiGauge (Fujifilm). The percentage of bound substrate was calculated from the intensity of the free substrate band as 100 × [1 - (I_sample_/I_control_)], in which I_control_ is the background-corrected intensity in the no-protein control. Binding curves were fitted by nonlinear regression in GraphPad Prism 10. Nonlinear fits were used to visualise concentration dependence; inferential comparisons used replicate-level values measured at the highest tested protein concentration.

### Helicase assays

SETX-HD variants at 35, 70, and 140 nM were incubated with 10 nM fluorescently labelled nucleic acid substrate in buffer H (25 mM Tris-Cl, pH 7.5, 1 mM DTT, 50 mM NaCl, 40 mM creatine phosphate, 55 μg/mL creatine kinase, 5 mM MgCl₂, and 2 mM ATP) for 30 min at 37 °C. For RNA-containing substrates, RNasin Ribonuclease Inhibitor (Promega) was included at 2 U/μL. Reactions were stopped by adding 1/10 volume of SDS/proteinase K solution (2.5% (w/v) SDS and 0.5 mg/mL proteinase K) and incubating for 3 min at 37 °C. After addition of loading buffer (60% (v/v) glycerol and 0.005% Orange G), samples were resolved on 10% (w/v) native polyacrylamide gels in 1× TBE at 80 V for 1 h. Fluorescent species were imaged using a Typhoon FLA-9000 Starion scanner and quantified using MultiGauge (Fujifilm). The percentage of unwound product was plotted against protein concentration and fitted by nonlinear regression in GraphPad Prism 10. Nonlinear fits were used to visualise concentration dependence; inferential comparisons used replicate-level values measured at the highest tested protein concentration.

SETX-CT rescue experiments (Supplementary Fig. 14f) were performed essentially as described above, except that SETX-HD^ΔCT^ (100 nM) was pre-incubated with SETX-CT (33, 100, or 300 nM) on ice for 5 min before initiation of the reaction. SETX-HD (100 nM) was included as a positive control.

### Biolayer interferometry (BLI)

Binding kinetics were measured by biolayer interferometry using an Octet RED96e instrument in Advanced Kinetics mode (ForteBio) at 25 °C with orbital shaking at 1,000 rpm. Streptavidin biosensors (SAX, ForteBio) were prehydrated for 10 min in BLI buffer (buffer D supplemented with 0.05% (v/v) Tween 20). Biotinylated ssDNA (pRMS739) and ssRNA (pRMS740) were loaded separately at 12 nM; sequences are listed in Supplementary Table S2. For ssRNA measurements, RNasin Ribonuclease Inhibitor (Promega) was included in BLI buffer at 0.5 U/μL. Association was recorded for 180 s using SETX-HD variants at 12.5, 25, and 50 nM and SETX-CT at 0.5, 1, and 2 μM, respectively. Dissociation was recorded for 120 s after transferring the biosensors to analyte-free BLI buffer. Sensorgrams report the change in optical thickness (nm) over time. Sensorgrams from three independent experiments at each concentration were fitted globally using the ‘Association then dissociation’ nonlinear-regression model in GraphPad Prism 10. Analyte concentrations (HotNM) were fixed to their nominal values, and association and dissociation rate constants were shared across datasets. Apparent equilibrium dissociation constants were calculated from the globally fitted kinetic rate constants as the ratio of the dissociation rate constant to the association rate constant. Goodness of fit was assessed from the coefficient of determination (R^2^).

### Pull-down assays using HEK293T cell lysate

Pull-down assays were performed as described previously^74^, with the following conditions. HEK293T cells were cultured in Dulbecco’s modified Eagle’s medium (DMEM; Sigma) supplemented with 10% (v/v) Tet-free-approved foetal bovine serum (Sigma), 100 U/mL penicillin, and 100 μg/mL streptomycin (Gibco) at 37 °C in a humidified atmosphere containing 5% CO₂. A cell pellet from one 10-cm dish was used per condition and resuspended in 1 mL NETN buffer (50 mM Tris-Cl, pH 8, 150 mM NaCl, 1 mM EDTA, 0.5% NP-40, and 0.1 mg/mL BSA) supplemented with Benzonase nuclease (2 U/μL) and cOmplete EDTA-free protease inhibitors. Lysates were incubated for 30 min at 4 °C, briefly sonicated, and clarified by centrifugation at 13,000 × g for 20 min at 4 °C. Each clarified lysate was incubated with 5 μg FLAG-tagged SETX-HD or SETX-HD^Sen1prong^ for 30 min at 4 °C, transferred to 25 μL ANTI-FLAG M2 Affinity Gel (Sigma) pre-equilibrated in NETN buffer, and incubated for a further 30 min at 4 °C. Beads were washed four times with four column volumes of NETN buffer. Bound proteins were eluted in 50 μL NETN buffer containing 0.25 mg/mL 3×FLAG peptide (Sigma). Input and eluate fractions were analysed by western blotting using antibodies against the FLAG epitope (Sigma, dilution 1:1000) and the RPB1 C-terminal domain (clone 8WG16, dilution 1:500), respectively.

#### *In vitro* transcription assays

*In vitro* transcription assays were performed as described previously^80^, with the following modifications. Promoter-independent ternary elongation complexes (ECs) were assembled by annealing fluorescently labelled RNA (DL2492) to template DNA and incubating 1 pmol of the resulting RNA-DNA hybrid with 0.5 pmol purified RNAPII for 10 min at 20 °C. A twofold molar excess of non-template (NT) strand was then added, followed by a further 10-min incubation at 20 °C. ECs were immobilised on Dynabeads MyOne Streptavidin T1 beads (Invitrogen) and washed sequentially with transcription buffer (TB; 20 mM Tris-HCl, pH 7.5, 100 mM NaCl, 8 mM MgCl₂, 10 μM ZnCl₂, 10% (v/v) glycerol, and 2 mM DTT) containing 0.1% (v/v) Triton X-100, TB containing 0.5 M NaCl, and TB alone. Transcription reactions (20 μL) were performed in TB with 400 nM SETX-HD or the indicated SETX-HD variant unless stated otherwise. For SETX-HD titration experiments, reactions contained 0, 400, or 800 nM SETX-HD. For the G-less cassette/G-rich tract template, DL3353 and DL3352 served as template and non-template strands, respectively, and transcription was initiated with a nucleotide mixture containing ATP, UTP, and CTP (1 mM each as final concentration). For the pause-inducing T-tract template, DL4607 and DL4604 served as template and non-template strands, respectively, and reactions contained 2 mM ATP, 1 mM GTP, 1 mM CTP, and 0.5 mM UTP. Reactions were incubated for 5 min at 37 °C and stopped with 1 μL of 0.5 M EDTA. After separation of beads and supernatant fractions, supernatant RNA was ethanol-precipitated and resuspended in 8 μL of loading buffer (1× Tris-borate-EDTA, 8 M urea). To measure RNAPII release, bead fractions were resuspended in 8 μL of loading buffer, heated at 95 °C for 5 min, and loaded separately from the supernatant fractions on 10% (w/v) denaturing polyacrylamide gels containing 8 M urea. To analyse the overall distribution of transcribing RNAPII on pause-inducing templates, beads and supernatant fractions were combined before heating and electrophoresis. Gels were imaged using a Typhoon scanner and quantified using ImageQuant TL (GE Healthcare).

### Statistical analyses

Statistical analyses were performed using GraphPad Prism 10 (version 10.6.1). Quantitative data are presented as mean ± s.d. from independent experiments, with the number of independent experiments specified in the corresponding figure legends. Normality was assessed using the Shapiro-Wilk test. Comparisons between two groups were performed using two-sided unpaired Student’s *t*-tests; Welch’s correction was applied to ATPase data to account for unequal variances. For EMSA binding curves and helicase unwinding curves, inferential comparisons used replicate-level values measured at the highest tested protein concentration; nonlinear fits were used to visualise concentration dependence. For BLI analyses, endpoint values from three independent experiments were compared as indicated in the figure legends. Statistical significance was defined as *P* < 0.05. *P* values are indicated as follows: ns, *P* > 0.05; *, *P* < 0.05; **, *P* < 0.01; ***, *P* < 0.001; and ****, *P* < 0.0001. The *P* values are indicated in the corresponding figure legends.

### Negative-stain electron microscopy of SETX-HD^ΔCT^

Peak fractions from SETX-HD^ΔCT^ heparin chromatography were diluted to 0.02 mg/mL in 50 mM Tris-Cl, pH 7.5, 300 mM NaCl, 1 mM DTT, and 0.01% NP-40. A 4 μL aliquot was applied to a freshly glow-discharged, carbon-coated, 300-mesh Cu grid (Quantifoil) and incubated for 45 s. The grid was washed once with a drop of Nano-W stain (Nanoprobes), blotted, and stained with a second fresh drop of Nano-W for 45 s. Excess stain was blotted away, and the remaining thin film was allowed to air-dry in a fume hood.

Micrographs were collected using a Thermo Fisher Scientific Talos F200C transmission electron microscope operated at 200 kV and equipped with a Thermo Fisher Scientific Falcon 3EC direct electron detector operated in integrating mode. A 100 μm objective aperture was used. Data were acquired in SerialEM^81^ at a pixel size of 1.598 Å pixel^-1^, with a defocus range of 1.5-3.0 μm, a total electron dose of 30 e^-^ Å^-2^, and an exposure time of 2.5 s. The dataset comprised 1,595 micrographs.

Micrographs were imported into cryoSPARC^82–83^ version 4.7.1, and CTF parameters were estimated using Patch CTF. Parameter outliers and micrographs with substantial contamination or imaging defects were removed, leaving 1,037 micrographs. Particles were selected using

Blob Picker and extracted in a 256-pixel box with twofold binning to 128 pixels, yielding 410,888 particles. Two rounds of 2D classification removed junk and other undesired particles. The remaining 69,359 particles were re-extracted in a 150-pixel box and subjected to six-class *ab initio* reconstruction. The class corresponding to SETX-HD^ΔCT^ (10,309 aligned particles) was refined using Non-uniform Refinement to generate the final map at a nominal resolution of 10.2 Å by gold-standard FSC.

### Cryo-EM sample preparation

LMNG (lauryl maltose neopentyl glycol) was added to 0.002% (v/v) immediately before grid preparation. A 3.5 μL aliquot of the sample was applied to a freshly glow-discharged Au-Flat R1.2/1.3, 300-mesh grid (Protochips) and vitrified in liquid ethane using a Thermo Fisher Scientific Vitrobot Mark IV at 4 °C and 100% relative humidity, with a 6 s blot time and 30 s wait time. Vitrified grids were mounted in autogrid cartridges for cryo-EM data collection.

### Cryo-EM data acquisition

Movies were collected using a Thermo Fisher Scientific Titan Krios transmission electron microscope operated at 300 kV, equipped with a Gatan K3 direct electron detector and a Gatan BioQuantum energy filter operated in zero-loss mode with a 10 eV slit. The K3 detector was operated in electron-counting and correlated-double-sampling (CDS) modes. Data were acquired in SerialEM^81^ at a pixel size of 0.5113 Å pixel^-1^ and a defocus range of 1.0-3.0 μm. Each movie comprised 49 frames acquired over 2.5 s with a total dose of 60 e^-^Å^-2^. Three datasets collected on three independently prepared samples comprised 9,842, 19,821, and 12,262 movies.

### Cryo-EM data processing

For concise labelling, SETX-HD^ATP-Mg2+^ denotes the ATP-compatible nucleotide-bound state obtained in the presence of ATP and Mg^2+^, and SETX-HD^R-loop^ denotes the 5′-oh R-loop-associated reconstruction. These are operational sample-state labels and do not imply unambiguous ligand identity or strand/duplex assignment.

Movies were imported into cryoSPARC^82–84^ version 4.7.1 and processed using Patch Motion Correction for alignment, motion correction, and dose weighting. CTF parameters were estimated using Patch CTF. Parameter outliers and micrographs with substantial contamination or imaging defects were removed, leaving three datasets of 7,762, 16,903, and 8,894 micrographs. For each dataset, 500 randomly selected micrographs were used to train a Micrograph Denoiser model after Patch Motion Correction; the resulting model was then applied to the corresponding full dataset. Particles were selected from denoised micrographs using Blob Picker and extracted in a 672-pixel box with eightfold binning to 84 pixels, yielding 625,131, 2,310,865, and 1,492,052 particles. Each dataset underwent two rounds of 2D classification, leaving 291,364, 482,049, and 487,289 particles, respectively. The three particle sets were combined for subsequent processing.

A multiclass *ab initio* reconstruction was used to generate an initial SETX-HD^ATP-Mg2+^ reference. The resulting volumes and a SETX-HD^R-loop^ reference from a separate dataset were used in iterative Heterogeneous Refinement. After each of three iterations, only particles assigned to the SETX-HD^ATP-Mg2+^ or SETX-HD^R-loop^ classes were retained. The 82,214 particles assigned to SETX-HD^ATP-Mg2+^ were re-extracted in a 400-pixel box with twofold binning to 200 pixels and subjected to NU Refinement and Local Refinement. Reference-based motion correction removed 4,811 particles. The remaining particles underwent NU Refinement, Local CTF Refinement, and Global CTF Refinement followed by post-processing. The final 77,403 particles yielded a consensus SETX-HD^ATP-Mg2+^ map at 3.83 Å global resolution (FSC = 0.143).

To improve the prong density, 283,799 SETX-HD^ATP-Mg2+^ particles from the first Heterogeneous Refinement iteration were re-extracted in a 400-pixel box with twofold binning to 200 pixels and refined using NU Refinement. Particles were imported into RELION 5.0^85^ and subjected to one round of masked, alignment-free 3D classification with Blush refinement^86^. The class with the strongest prong density contained 96,623 particles. This class was refined with masking and Blush refinement but without alignment, followed by post-processing. The focused SETX-HD^prong^ map reached a global resolution of 4.1 Å (FSC = 0.143).

To reconstruct the 5′-oh R-loop-associated state, 51,574 SETX-HD^R-loop^ particles were selected after four iterations of Heterogeneous Refinement, re-extracted in a 400-pixel box with twofold binning to 200 pixels, refined using NU Refinement, and imported into RELION 5.0^85^. Particles were subjected to 3D classification with Blush refinement, and the best class was refined further with Blush enabled. The resulting 37,195 particles were transferred back to cryoSPARC, re-extracted in a 600-pixel box with twofold binning to 300 pixels, and reconstructed using Homogeneous Reconstruction Only, followed by post-processing. The final SETX-HD^R-loop^ map reached a global resolution of 6.48 Å (FSC = 0.143).

A composite SETX-HD^ATP-Mg2+^ map was assembled from the consensus and prong-focused maps. A mask generated from the low-pass-filtered consensus map was divided into a prong segment and a segment encompassing the remainder of the volume. The SETX-HD^prong^ map was masked with the prong segment, whereas the consensus SETX-HD^ATP-Mg2+^ map was masked with the complementary segment. Contour levels were scaled to match, and the two masked maps were combined in ChimeraX^87^.

### AlphaFold 3 modelling

AlphaFold 3^88^ predictions were generated using the AlphaFold Server. Two classes of models were produced: (i) an initial SETX-HD^ATP-Mg2+^ model, used for fitting and model building in the corresponding cryo-EM map; and (ii) dsDNA, dsRNA, and RNA-DNA hybrid models, used to evaluate the placement of nucleic acid duplexes within the SETX-HD^R-loop^ map. Prediction-quality metrics and superpositions of the five highest-ranked models are shown in Supplementary Fig. 8. The top-ranked models used in these analyses were deposited in the ModelArchive^89^ database under accession codes ma-np8y0 (SETX-HD^ATP-Mg2+^), ma-5no17 (dsDNA), ma-k09z8 (dsRNA), and ma-l3hs8 (RNA-DNA hybrid).

### Model building of SETX-HD^ATP-Mg2+^

An initial model comprising SETX-HD residues 1699-2454, one ATP molecule, and one Mg^2+^ ion was predicted using AlphaFold 3^88^ and fitted as a rigid body into the composite map in ChimeraX. The prong and β-barrel subdomains were adjusted manually in ISOLDE^90^ with secondary-structure restraints enabled. The model was then imported into Coot 1.1.18^91^ for manual real-space refinement and extended to residues 1695-2465. Automated real-space refinement and geometry validation were performed in Phenix^92^. Side chains lacking sufficient density were removed, and residues lacking backbone support were omitted. Cycles of manual and automated refinement were repeated until model geometry and agreement with the experimental map were jointly optimised. The final coordinates were deposited in the Protein Data Bank under accession code 9TBG.

### Data visualisation

Atomic models, cryo-EM maps, and structural superpositions were visualised using UCSF ChimeraX^87^ or UCSF Chimera^93^. Electrostatic surface potential was calculated and visualised using default settings of the coulombic command in UCSF ChimeraX. Multiple sequence alignments were edited and visualised in Jalview^94^. Structure-guided multiple sequence alignments were generated, edited, and visualised in UCSF Chimera. Sequence logos were generated using WebLogo 3^95^. Biochemical data were plotted using GraphPad Prism 10 (version 10.6.1). Phylogenetic trees were visualised using TreeViewer^96^.

### Measurements of structural movements

Structural movements between SETX-HD and Sen1 models were quantified in UCSF ChimeraX. First, each pair of structures was superposed on the RecA1 subdomain, which served as the static reference, using Matchmaker. The structures were then superposed on the moving region of interest, either the RecA2 subdomain or the β-barrel-stalk-brace assembly, and the rotation relative to the RecA1-aligned reference was calculated using the measure rotation command.

### *In silico* evolutionary analysis of the C terminus of SETX-HD

SETX-HD residues 2100-2677 (UniProt accession Q7Z333-1) were used as the query for BLAST^97^ searches of the non-redundant protein sequence database (nr), restricted to Animalia (taxid: 33208). Searches used a maximum of 5,000 target sequences, a word size of 3, and low-complexity filtering. Hits were removed if query coverage was <60%, if the sequence was annotated as partial, low quality, hypothetical, or unknown, or if the C terminus was absent. When multiple entries or isoforms were available for one organism, only the longest sequence was retained. When multiple species from a single genus were available, only two were retained. The filtered set of 768 sequences was aligned using the EMBL-EBI implementation^98^ of Kalign 3^99^ with the following custom settings: terminal gap penalty 1; gap-extension penalty 2. The alignment was reduced to the section of the C terminus (human sequence 2468-end) in Jalview, and a phylogenetic tree was inferred using FastTree^100^. The full tree was visualised in TreeViewer^96^, rooted using fungal Sen1 sequences as the outgroup, and manually pruned to 51 representative sequences. Branch lengths were retained from the FastTree output.

### Use of large language models

ChatGPT (GPT-6 model, OpenAI) was used during manuscript preparation to assist with language editing and improvement of the clarity and consistency of the text. All scientific content, interpretations, and conclusions were determined by the authors, who critically reviewed and revised all LLM-assisted text and take full responsibility for the final manuscript.

## Data availability

All primary data are available in this manuscript, supplementary information, and source data. Structural data are available in PDB database under the accession number 9TBG and in EMDB database under accession numbers EMD-55770, EMD-55767, EMD-55768, EMD-55798, and EMD-55850. AlphaFold-generated models are available from ModelArchive under accession numbers ma-np8y0, ma-5no17, ma-k09z8, and ma-l3hs8.

## Supplementary data

Supplementary data are available online.

## Acknowledgements

We thank members of the laboratory for their support. We are grateful to Dalibor Blazek (CEITEC, Masaryk University), Lumir Krejci, Jakub Cibulka (Faculty of Medicine, Masaryk University), Olga Jasnovidova (Tallinn University of Technology), and Martin Pacesa (University of Zurich) for helpful discussions and critical reading of the manuscript. We thank Karel Skubnik and Jiri Novacek (Cryo-Electron Microscopy and Tomography Core Facility, CEITEC, Masaryk University), and David Buchta, Jan Petrovsky, David Zapletal (CEITEC, Masaryk University), and Martin Pacesa (University of Zurich) for advice and assistance with cryo-EM data analysis and structure determination. David Zapletal also assisted with purification of the SETX-HD^R-loop^ complex. We thank Veronika Klapstova and Nikola Noskova (CEITEC, Masaryk University) for assistance with RNAPII purification, and Josef Houser (Biomolecular Interactions and Crystallization Core Facility, CEITEC, Masaryk University) for assistance with biolayer interferometry.

## Author contributions

Experimentation: ML, KS, ZH, OP; Protein purification: ML, ZH; *In silico* modelling: ML; Structure determination: ML; Conceptualisation: OP, MS; Funding acquisition: OP, MS; Project administration: MS; Supervision: MS; Writing -original draft: ML, OP, MS; Writing -review and editing: ML, KS, ZH, OP, MS

