## Supplementary Material for "Human senataxin uses distinct peripheral elements to unwind R-loops and act on RNA polymerase II"

Supplementary Figure legends

Supplementary Figures: 1-20

Supplementary Tables: S1-S4

#### SUPPLEMENTARY FIGURE LEGENDS

**Supplementary Fig. 1 | Preparation of SETX-HD with a 5′-oh R-loop for cryo-EM. a,** Schematic purification workflow and Coomassie-stained SDS-PAGE analysis of purified SETX-HD. The indicated volumes of concentrated protein were loaded; M, molecular-mass marker. **b,** Oligonucleotide organisation of the three-stranded 5′-oh R-loop used for cryo-EM. The RNA strand is pRMS222, and the DNA strands are pRMS219 and pRMS217; the 5′ end of pRMS217 carries a Cy3 fluorophore. Oligonucleotide sequences are listed in Supplementary Table S2. **c,** Size-exclusion chromatography of SETX-HD, the 5′-oh R-loop, and the assembled complex. Peak fractions were analysed by SDS-PAGE and Cy3-detected native PAGE. Fraction D4 was used for downstream cryo-EM analysis.

**Supplementary Fig. 2 | Cryo-EM processing of nucleotide-bound and 5′-oh R-loop-associated SETX-HD datasets.** Representative denoised micrograph, particle-selection workflow, and final reconstructions. Three independently collected datasets were initially processed separately in cryoSPARC and then combined. Iterative heterogeneous refinement separated particles contributing to the nucleotide-bound SETX-HD consensus reconstruction, the focused prong reconstruction, and the 5′-oh R-loop-associated reconstruction. Selected particles were refined in cryoSPARC or transferred to RELION 5.0 for classification and refinement as indicated. The final particle sets contained 77,403 particles for the 3.83-Å nucleotide-bound consensus reconstruction, 96,623 particles for the 4.1-Å focused prong reconstruction, and 37,195 particles for the 6.48-Å 5′-oh R-loop-associated reconstruction. Representative two-dimensional class averages of the final particle sets are shown. For concise labelling, SETX-HD$^{ATP\text{-}Mg^{2+}}$ denotes the ATP-compatible nucleotide-bound state obtained in the presence of ATP and $Mg^{2+}$, and SETX-HD$^{R\text{-}loop}$ denotes the 5′-oh R-loop-associated reconstruction. These labels describe the respective sample states used for reconstruction.

**Supplementary Fig. 3 | Validation of the SETX-HD cryo-EM reconstructions. a,b,** Gold-standard Fourier shell correlation (FSC) curve (a) and particle-viewing distribution (b) for the 5′-oh R-loop-associated reconstruction at 6.48 Å. **c,d,** Gold-standard FSC curve (c) and particle-viewing distribution (d) for the nucleotide-bound SETX-HD consensus reconstruction at 3.83 Å. **e,f,** Gold-standard FSC curves (e) and angular distribution (f) for the focused prong reconstruction. **g,** Composite nucleotide-bound SETX-HD map coloured by local-resolution estimates calculated for the corresponding consensus and prong-focused reconstructions and shown in three views. Because the composite display combines local-resolution estimates from the consensus and prong-focused reconstructions, values close to the interface between the two map components, at the base of the prong subdomain, appear artificially low.

**Supplementary Fig. 4 | Structure-guided alignment of human SETX-HD and fungal Sen1 helicase domains.** Structure-guided sequence alignment of human SETX-HD (PDB 9TBG), *Saccharomyces cerevisiae* Sen1 (*Sc*Sen1; PDB 5MZN), and *Chaetomium thermophilum* Sen1 (*Ct*Sen1; PDB 8FTM), generated from structural superposition in UCSF Chimera. Residues matched by the structural alignment are outlined in black, unresolved regions are outlined in red, and grey bars above the alignment report local Cα root-mean-square deviations.[59,63]

**Supplementary Fig. 5 | Peripheral organisation and tracking-strand contact conservation across SETX-HD and fungal Sen1. a,b,** Superposition of nucleotide-bound SETX-HD (PDB 9TBG) with ADP-bound *Sc*Sen1 (PDB 5MZN; a) or RNA-bound *Ct*Sen1 (PDB 8FTM; b) after alignment on RecA1. The fungal structures are shown in grey, and the SETX β-barrel, stalk, and brace are coloured as indicated. Dashed axes indicate approximate outward displacements of 9° relative to *Sc*Sen1 and 12° relative to *Ct*Sen1. **c,** Three-way structural comparison of the tracking-strand interaction network. Thirty distinct RNA-contact positions reported for RNA-bound *Ct*Sen1 (PDB 8FTM) and *Sc*Sen1 (PDB 8RAN) were considered. Twenty corresponding positions are represented in the SETX-HD model (PDB

9TBG). Structural alignment was used to assign the corresponding residues in SETX-HD, *Ct*Sen1, and *Sc*Sen1 at each of these positions, including positions for which a tracking-strand contact was reported in only one of the fungal structures. Residue identities for all three proteins are labelled. Fifteen of the 20 positions are identical across all three proteins.[59,63,64]

**Supplementary Fig. 6 | Nucleotide density in the SETX-HD active site.** Close view of the nucleotide-binding cleft in the ATP-compatible nucleotide-bound SETX-HD map. The fitted ATP model is shown as sticks, the putative $Mg^{2+}$ ion as a green sphere, and the corresponding cryo-EM density as a cyan mesh. The density is compatible with the adenosine, ribose, and three phosphate groups of the fitted ATP-$Mg^{2+}$ model under the sample conditions.

**Supplementary Fig. 7 | Comparison of alternative duplex models with the 5′-oh R-loop-associated density. a-c,** AlphaFold 3 models of a 21-nt RNA-DNA hybrid (a), dsRNA (b), or dsDNA (c) fitted as rigid bodies into the duplex-associated density of the 5′-oh R-loop-associated SETX-HD map. Two views related by 90° are shown. Fits were evaluated in UCSF ChimeraX by simulating 6.5Å maps of the duplexes and calculating map-to-map correlation. The displayed correlation coefficients are approximately 0.86 for the RNA-DNA hybrid and 0.84 for both dsRNA and dsDNA. The similar correlations indicate that the reconstruction does not discriminate among the three duplex models.

**Supplementary Fig. 8 | AlphaFold 3 models used in the structural analyses. a-c,** Five independently seeded AlphaFold 3 predictions of the 21-nt dsDNA (a), dsRNA (b), and RNA-DNA hybrid (c) models used in Supplementary Fig. 7. **d,** Predictions of SETX-HD residues 1699-2454 with one ATP molecule and one $Mg^{2+}$ ion. For each target, the five highest-ranked predictions are superposed on the left, and the top-ranked model is coloured by pLDDT on the right. Top-ranked models were deposited in ModelArchive as ma-5no17 (dsDNA), ma-k09z8 (dsRNA), ma-l3hs8 (RNA-DNA hybrid), and ma-np8y0 (SETX-HD).

**Supplementary Fig. 9 | Basic residues in the candidate nucleic-acid channel support RNA-stimulated ATP turnover. a,** Electrostatic surface potential of nucleotide-bound SETX-HD with the additional density from the 5′-oh R-loop-associated map shown in yellow. Positions selected for alanine substitution are labelled. **b,** Multiple-sequence alignment of selected SETX/Sen1 proteins around the substituted residues. **c,** Coomassie-stained SDS-PAGE analysis of purified SETX-HD variants, with the purification workflow shown at left; M, molecular-mass marker. **d,** Apparent turnover numbers ($k_{cat,app}$) for RNA-stimulated ATP turnover. R2168A and its corresponding SETX-HD control were analysed in a separate experimental batch, as indicated. Data are mean ± s.d. from three independent experiments. After assessment of normality by the Shapiro-Wilk test, variants were compared with the corresponding SETX-HD control using two-sided Welch's *t*-tests. The unadjusted $P$ values for the comparisons indicated by asterisks are: batch 1, R1899A, $P < 0.001$; R1904A, $P < 0.001$; K2129A/K2131A, $P = 0.00206$; K2131A, $P = 0.00622$; R2136A, $P = 0.0113$; K2139A, $P < 0.0001$; R2198A, $P = 0.00391$; and R2235A, $P < 0.0001$; batch 2, R2168A, $P = 0.00920$.

**Supplementary Fig. 10 | ssRNA binding by SETX-HD channel variants. a,** Apparent equilibrium dissociation constants obtained by biolayer interferometry (BLI) for SETX-HD and the indicated alanine-substitution variants. **b-k,** BLI sensorgrams for SETX-HD (b), R1899A (c), R1904A (d), K2131A (e), K2129A/K2131A (f), R2136A (g), K2139A (h), R2168A (i), R2198A (j), and R2235A (k). Biotinylated 41-nt ssRNA was immobilised on streptavidin biosensors, and proteins were analysed at 12.5, 25, and 50 nM. Association was monitored for 180 s and dissociation for 120 s. Sensorgrams were obtained from three independent experiments at each concentration; global fits to an association-then-dissociation model are shown, with $R^2$ values indicated. **l,** Control sensorgrams obtained without immobilised ssRNA.

**Supplementary Fig. 11 | Selective divergence at two *Ct*Sen1 tracking-strand contact positions. a,** Structural comparison of nucleotide-bound SETX-HD (PDB 9TBG; cyan) with RNA-bound *Ct*Sen1 (PDB 8FTM; grey) at the position corresponding to *Ct*Sen1 R1427, which contacts the tracking-strand backbone and has been proposed to function as a ratchet pawl. The corresponding SETX P2048 and *Sc*Sen1 R1444 positions are labelled. **b,** Comparison at the position corresponding to *Ct*Sen1 H1556, which contributes a polar contact within the 3′ tracking-strand grip. The corresponding SETX G2160 and *Sc*Sen1 H1574 positions are labelled. **c,d,** Structure-guided local alignments around SETX P2048 (c) and G2160 (d) for SETX-HD, *Sc*Sen1, and *Ct*Sen1. **e,f,** Multiple-sequence alignments of the regions surrounding P2048 (e) and G2160 (f) in selected animal SETX proteins and fungal Sen1 orthologues. Arrows mark the positions corresponding to the *Ct*Sen1 tracking-strand contacts. **g,h,** Sequence logos generated from the alignments in e and f, respectively.[59,63]

**Supplementary Fig. 12 | Conservation of the jawed-vertebrate-associated C-terminal basic motif in selected SETX/Sen1 sequences. a,** Multiple-sequence alignment of the SETX/Sen1 C-terminal region. Human SETX residues 2527 and 2560 delimit the segment removed in SETX-HD$^{\Delta2527\text{-}2560}$, and the basic motif is indicated. Residues are shaded according to percentage identity. The motif is readily recognisable across the displayed jawed-vertebrate SETX sequences and is not recognisable in the displayed lamprey, lancelet, or fungal Sen1 sequences. **b,** Sequence logo generated from the alignment in a. The sampled alignment supports the motif as a jawed-vertebrate-associated feature.

**Supplementary Fig. 13 | Phylogenetic distribution of the SETX C-terminal basic motif.** Phylogenetic tree inferred from aligned C-terminal SETX/Sen1 sequences. Branches and labels coloured magenta identify displayed sequences in which the basic motif defined in Supplementary Fig. 12 was not recognisable; motif-containing displayed sequences are shown in black. The scale bar indicates estimated amino-acid substitutions per site. Colouring reports motif-scoring status across the sampled sequences and summarises the observed phylogenetic distribution of the motif.

**Supplementary Fig. 14 | Biochemical characterisation of SETX-HD C-terminal deletion variants. a,** Coomassie-stained SDS-PAGE analysis of purified SETX-HD, SETX-HD$^{\Delta CT\text{-}RBD}$ (deletion of residues 2527-2560), SETX-HD$^{\Delta CT,}$ and catalytically inactive SETX-HD$^{\Delta CT,D2181A/E2182A,}$ with the purification workflow shown below. **b-d,** Representative native PAGE assays of 5′-oh R-loop unwinding by increasing concentrations of SETX-HD (b), SETX-HD$^{\Delta CT}$ (c), and SETX-HD$^{\Delta CT\text{-}RBD}$ (d) in the presence or absence of ATP. **e,** Quantification of b-d. **f,** Test of whether purified SETX-CT added *in trans* rescues 5′-oh R-loop unwinding by SETX-HD$^{\Delta CT}$. **g,** Quantification of f. **h-j,** EMSAs comparing binding of SETX-HD (h), SETX-HD$^{\Delta CT}$ (i), and SETX-HD$^{\Delta CT\text{-}RBD}$ (j) to the 5′-oh R-loop, ssRNA, and ssDNA. **k,l,** Quantification of 5′-oh R-loop binding (k) and ssRNA and ssDNA binding (l). Data are mean ± s.d. from three independent experiments. Values at the highest protein concentrations (140 nM in e and 60 nM in k,l) were compared using two-sided unpaired Student's *t*-tests without adjustment for multiple comparisons. The *P* values for the indicated comparisons are: **e,** SETX-HD versus SETX-HD$^{\Delta CT\text{-}RBD,}$ $P < 0.0001$; SETX-HD versus SETX-HD$^{\Delta CT,}$ $P < 0.001$; and SETX-HD$^{\Delta CT}$ versus SETX-HD$^{\Delta CT\text{-}RBD,}$ $P = 0.0149$; **k,** SETX-HD versus SETX-HD$^{\Delta CT\text{-}RBD,}$ $P = 0.00577$; l, for ssRNA binding, SETX-HD versus SETX-HD$^{\Delta CT,}$ $P = 0.0245$, and SETX-HD versus SETX-HD$^{\Delta CT\text{-}RBD,}$ $P = 0.0239$. Non-significant comparisons are not marked.

**Supplementary Fig. 15 | Negative-stain analysis of SETX-HD$^{\Delta CT}$. a,** Representative negative-stain micrograph and particle-processing workflow for SETX-HD$^{\Delta CT}$. Particles were selected and classified in cryoSPARC, and the indicated class was refined from 10,309 particles to a map with a nominal resolution of 10.2 Å by gold-standard FSC. Representative two-dimensional class averages are shown. **b,c,** Gold-standard FSC curve (b) and particle-viewing distribution (c) for the SETX-HD$^{\Delta CT}$ reconstruction. **d,** Negative-stain density map of SETX-HD$^{\Delta CT}$ shown in two views related by 90°. **e,**

Composite ATP-compatible nucleotide-bound SETX-HD cryo-EM map low-pass filtered to 15 Å and shown in corresponding views for comparison. The maps are compared at the level of their overall envelopes.

**Supplementary Fig. 16 | Nucleic-acid binding properties of the isolated SETX C-terminal extension. a,** Coomassie-stained SDS-PAGE analysis of SETX-CT (residues 2455-2677), SETX-CT$^{\Delta CT\text{-}RBD}$ (deletion of residues 2527-2560), and SETX-CT$^{\Delta 2527\text{-}2590}$, with the purification workflow shown at left. **b-d,** EMSAs of SETX-CT with 61-nt (b), 41-nt (c), or 15-nt (d) ssRNA; **e,** quantification of EMSAs shown in b-d. **f,g,** EMSAs of SETX-CT with 61-nt ssDNA (f) or the 5′-oh R-loop (g); **h,** quantification comparing 61-nt ssDNA, 61-nt ssRNA, and the 5′-oh R-loop. **i,j,** EMSAs of SETX-CT$^{\Delta CT\text{-}RBD}$ (i) and SETX-CT$^{\Delta 2527\text{-}2590}$ (j) with 61-nt ssRNA; **k,** quantification together with SETX-CT. **l-q,** BLI sensorgrams for SETX-HD (l,o), SETX-HD$^{\Delta CT\text{-}RBD}$ (m,p), and SETX-CT (n,q) binding to 41-nt ssRNA (l-n) or ssDNA (o-q). **r-t,** Normalised RNA-versus-DNA sensorgrams for SETX-HD (r), SETX-HD$^{\Delta CT\text{-}RBD}$ (s), and SETX-CT (t). **u,** Comparison of SETX-HD and SETX-HD$^{\Delta CT\text{-}RBD}$ binding to ssDNA. **v,** BLI controls for buffer exposure to ssRNA- or ssDNA-loaded sensors and protein exposure to sensors without immobilised nucleic acid. EMSA data in e, h, and k are mean ± s.d. from three independent experiments. Values at 600 nM were compared using two-sided unpaired Student's *t*-tests without adjustment for multiple comparisons. The $P$ values for the indicated comparisons are: e, 61-nt versus 41-nt ssRNA, $P = 0.0293$; h, ssDNA versus ssRNA, $P < 0.0001$; ssDNA versus 5′-oh R-loop, $P < 0.001$; ssRNA versus 5′-oh R-loop, $P = 0.0247$; **k,** SETX-CT versus SETX-CT$^{\Delta CT\text{-}RBD}$, $P = 0.00177$, and SETX-CT versus SETX-CT$^{\Delta 2527\text{-}2590}$, $P < 0.001$. BLI measurements were performed in three independent experiments. The indicated endpoint values in r-u at the final recorded dissociation timepoint (299.803 s) were compared using two-sided unpaired Student's *t*-tests without adjustment for multiple comparisons. The $P$ values are: **r,** SETX-HD on ssRNA versus ssDNA, $P < 0.0001$; **s,** SETX-HD$^{\Delta CT\text{-}RBD}$ on ssRNA versus ssDNA, $P < 0.0001$; **t,** SETX-CT on ssRNA versus ssDNA, $P = 0.0328$; **u,** SETX-HD versus SETX-HD$^{\Delta CT\text{-}RBD}$ on ssDNA, $P = 0.0255$. Non-significant comparisons are not marked.

**Supplementary Fig. 17 | SETX-HD redistributes endpoint transcripts at a defined T-rich RNAPII pause. a,** Sequence of the immobilised elongation-complex substrate, showing the RNA-annealing region and transcript positions classified as upstream, $T_3$-$T_4$ pause, downstream, and run-off. **b,** Representative denaturing PAGE analysis of combined bead and supernatant (B+S) RNA after incubation with 0, 400, or 800 nM SETX-HD. **c,** Quantification of upstream, $T_3$-$T_4$ pause, downstream, and run-off products in b. **d,e,** Quantification of run-off (d) and shorter upstream (e) products for buffer control, SETX-HD, SETX-HD$^{\Delta CT\text{-}RBD}$ (deletion of residues 2527-2560), and SETX-HD$^{\Delta CT}$. Data are mean ± s.d. from three independent experiments. Statistical comparisons used two-sided unpaired Student's *t*-tests without adjustment for multiple comparisons. The $P$ values for comparisons in c, relative to 0 nM SETX-HD at 400 and 800 nM, respectively, are: upstream, $P = 0.0348$ and $P < 0.001$; $T_3$-$T_4$ pause, $P = 0.0326$ and $P = 0.00242$; downstream, $P = 0.0109$ and $P < 0.001$; run-off, $P = 0.254$ and $P = 0.371$ (both marked ns). In d, buffer versus SETX-HD, $P = 0.0448$, and SETX-HD versus SETX-HD$^{\Delta CT\text{-}RBD}$, $P = 0.0255$. In e, buffer versus SETX-HD, $P < 0.001$. These assays report endpoint RNA populations across the indicated transcript classes.

**Supplementary Fig. 18 | Sequence variation within the SETX/Sen1 prong.** Multiple-sequence alignment of the region encompassing the prong in selected animal SETX proteins, *Sc*Sen1, and *Ct*Sen1. The prong boundaries are indicated above the alignment blocks. Residues are shaded according to percentage identity. Lineage-specific insertions within the prong are visible in fungal Sen1 and selected non-vertebrate sequences.

**Supplementary Fig. 19 | Biochemical characterisation of the SETX-HD$^{Sen1prong}$ chimera. a,** Solubility analysis of SETX-HD$^{\Delta prong}$, which lacks SETX residues 2061-2147. Coomassie-stained SDS-

PAGE and anti-His immunoblotting show the protein in total lysate, soluble supernatant, and pellet fractions; M, molecular-mass marker. **b,** Coomassie-stained SDS-PAGE of purified SETX-HD and SETX-HD$^{Sen1prong,}$ with the purification workflow shown at left. **c,d,** Representative EMSAs of increasing concentrations of SETX-HD (c) or SETX-HD$^{Sen1prong}$ (d) with the 5′-oh R-loop. **e,** Quantification of c and d; data are mean ± s.d. from three independent experiments. **f,g,** Quantification of shorter upstream (f) and run-off (g) endpoint RNA products in the T-rich pause assay comparing buffer control, SETX-HD, and SETX-HD$^{Sen1prong}$. Data are mean ± s.d. from three independent experiments. Statistical comparisons in f and g were performed against the buffer control using two-sided unpaired Student's *t*-tests without adjustment for multiple comparisons. The *P* values are: f, buffer versus SETX-HD, $P = 0.0156$, and buffer versus SETX-HD$^{Sen1prong,}$ $P = 0.996$ (ns); g, buffer versus SETX-HD, $P = 0.312$ (ns), and buffer versus SETX-HD$^{Sen1prong,}$ $P = 0.456$ (ns). Here, ns denotes not significant.

**Supplementary Fig. 20 | Pairwise RecA2 rotations among SETX-HD and fungal Sen1 structures. a-e,** Pairwise superpositions after alignment on RecA1: ATP-compatible nucleotide-bound SETX-HD (PDB 9TBG) and ADP·$BeF_3$-bound *Sc*Sen1 (PDB 8RAO; a); ADP·$BeF_3$-bound *Sc*Sen1 and ADP·$SO_4^{2-}$-bound *Ct*Sen1 (PDB 8FTM; b); ADP·$SO_4^{2-}$-bound *Ct*Sen1 and ADP-bound *Sc*Sen1 (PDB 5MZN; c); ADP-bound *Sc*Sen1 and nucleotide-free *Sc*Sen1 (PDB 8RAN; d); and nucleotide-free *Sc*Sen1 and ATP-compatible nucleotide-bound SETX-HD (e). Approximate RecA2 rotations of 10°, 21°, 18°, 17°, and 16°, respectively, are indicated. **f,** Schematic summary of these cross-orthologue comparisons arranged according to their nominal ligand states. Black dashed lines indicate rotation around the rotation axis and grey dashed lines indicate changes in rotation-axis tilt; approximate values are indicated in the panel. Measurements were performed in UCSF ChimeraX. The schematic summarises cross-orthologue structural comparisons arranged by nominal ligand state; the ordering is comparative rather than chronological and does not represent a directly observed catalytic sequence.[59,63,64]

**Supplementary Table S1: Cryo-EM and negative-stain EM data collection and refinement statistics.**

| | Data collection | | | |
|---|---|---|---|---|
| Sample | SETX-HD$^{ATP\text{-}Mg2+}$ | SETX-HD$^{R\text{-}loop}$ | SETX-HD$^{\Delta CT}$ (negative staining) | |
| EMDB accession number | EMD-55770* | EMD-55798 | EMD-55850 | |
| PDB accession number | 9TBG | - | - | |
| Microscope | Thermo Fisher Scientific Titan Krios | | Thermo Fisher Scientific Talos F200C | |
| Voltage (kV) | 300 | | 200 | |
| Detector | Gatan K3 (counting, CDS); Gatan BioQuantum energy filter, 10 eV slit | | Thermo Fisher Scientific Falcon 3EC (integrating mode) | |
| Symmetry | C1 | | C1 | |
| Electron dose ($e^-$/Å$^2$) | 60 | | 30 | |
| Defocus range (µm) | 1.0-3.0 | | 1.5-3.0 | |
| Pixel size (Å) | 0.5113 | | 1.598 | |
| Total collected data | 41925 movies | | 1595 micrographs | |
| Initial particles | 4428048 | | 410888 | |
| Final particles | 149662** | 37195 | 10309 | |
| | Refinement | | | |
| Masked map resolution 0.143/0.5 (Å) | 4.0/4.4*** | 6.5/7.7 | 10.2/16.8 | |
| Unmasked map resolution 0.143 (Å) | 4.6*** | 8.4 | 13.2 | |
| Map-to-model FSC 0.143/0.5 (Å) | 3.7/4.1 | - | - | |
| Map sharpening *B*-factor (Å$^2$) | **** | -50 | Unsharpened | |
| | R.m.s deviations | | | |
| Bond lengths (Å) | 0.002 | - | - | - |
| Bond angles (°) | 0.551 | - | - | - |
| | Validation | | | |
| MolProbity score | 1.51 | - | - | - |
| Clashscore | 8.01 | - | - | - |
| Poor rotamers (%) | 0.18 | - | - | - |
| | Ramachandran plot | | | |
| Favored (%) | 97.66 | - | - | - |
| Allowed (%) | 2.34 | - | - | - |
| Outliers (%) | 0 | - | - | - |

**Note:** SETX-HD$^{ATP\text{-}Mg2+}$ and SETX-HD$^{R\text{-}loop}$ are operational sample-state labels as defined in Supplementary Fig. 2.

*This accession code corresponds to the composite map. The codes for the consensus and focused maps are EMD-55767 and EMD-55768, respectively.

**These particles consist of two sets: 77403 particles for the consensus map and 96623 particles for the focused map. There is a 24364 particle overlap between the two sets.

*** For the composite map, the reported resolution values follow the EMDB recommendation for composite-map reporting and are the means of the corresponding FSC-derived resolutions of the constituent consensus and prong-focused reconstructions; they do not represent FSC values calculated directly from composite half-maps. The individual values are as follows: consensus map, masked 3.8/4.2 Å and unmasked 4.4 Å; focused prong map, masked 4.1/4.6 Å and unmasked 4.7 Å.

****The composite map was not sharpened globally by the same *B*-factor. The consensus map was sharpened globally with a *B*-factor of –176 Å$^2$ and also locally according to the local resolution using the Local filtering job in CryoSPARC. The focused map was sharpened globally with a *B*-factor of –102 Å$^2$.

**Supplementary Table S2: List of oligonucleotides used in this work.**

| Name | Sequence 5′-3′ | Use |
|---|---|---|
| pRMS103 | CCCTTCTCTTGCGTGATCGTGGCCGCAGCCGG ACAGTCTTGCGAAATCGAAACTC | Forward primer for site-directed mutagenesis to introduce D2181A/E2182A mutation. |
| pRMS104 | GAGTTTCGATTTCGCAAGACTGTCCGGCTGCG GCCACGATCACGCAAGAGAAGGG | Reverse primer for site-directed mutagenesis to introduce D2181A/E2182A mutation. |
| pRMS114 | ACCTCCTGCCTCTCTGTGC | Sequencing of SETX, starts at AA 1721 |
| pRMS115 | GGACTTCAACGAGGACCAAA | Sequencing of SETX, starts at AA 1923 |
| pRMS116 | CTGCTGCACCTTGTCTACCA | Sequencing of SETX, starts at AA 2152 |
| pRMS117 | AGTCGACACCGTTGATGCTT | Sequencing of SETX, starts at AA 2371 |
| pRMS217 | GACGCTGCCGAATTCTACCAGTGCCTTGCTAG GACATCTTTGCCCACCTGCAGGTTCACCC | 61-mer DNA oligo fluorescently labelled on 5′ end with Cy3 used to create R-loop for EMSAs and helicase assays. |
| pRMS219 | GGGTGAACCTGCAGGTGGGCGGCTGCTCATC GTAGGTTAGTTGGTAGAATTCGGCAGCGTC | 61-mer DNA oligo used to create R-loop substrates for EMSAs and helicase assays. |
| pRMS222 | UAAGAGCAAGAUGUUCUAUAAAAGAUGUCC UAGCAAGGCAC | 41-mer RNA oligo used to create a 5′-overhang R-loop for EMSAs and helicase assays. |
| pRMS323 | TTATCCACTTCCAATGTTATTATTAGTGATGG TGATGGTGATG | Reverse primer for generating a 1674-2677 amino acid fragment of SETX for cloning into 438-A plasmid. |
| pRMS590 | TTTAAGAAGGAGATATAGTTCATGGTCCTGAA CACCTTCG | Forward primer for generating a 1674-2677 amino acid fragment of SETX for cloning into 2Bc-T plasmid. |
| pRMS591 | GGATTGGAAGTAGAGGTTCTCGAGCAACTTA CGCTTCTTTG | Reverse primer for generating a 1674-2677 amino acid fragment of SETX for cloning into 2Bc-T plasmid. |
| pRMS592 | TACTTCCAATCCAATCGATGGTCCTGAACACC TTCG | Forward primer for generating a 1674-2677 amino acid fragment of SETX for cloning into 438-A plasmid. |
| pRMS678 | UUGUCUUCUCAUAAAUUUAAUCCCCGUACG CUUAUACUCCUUUAACUACCCAGUCUUCCCG | 61-mer RNA oligo fluorescently labelled on 5′ end with Cy3 used as a single-stranded RNA for EMSAs. |
| pRMS739 | TTGTCTTCTCATAAATTTAATCCCCGTACGCTT ATACTCCT | 41-mer 5′-biotin-tagged DNA primer for BLI assay |
| pRMS740 | UUGUCUUCUCAUAAAUUUAAUCCCCGUACG CUUAUACUCCU | 41-mer 5′-biotin-tagged RNA primer for BLI assay |
| DL2492 | UGCAUUUCGACCAGGC | 5′-FAM-labelled RNA used to assemble immobilised RNAPII elongation complexes. |

| **Name** | **Sequence 5′-3′** | **Use** |
|---|---|---|
| DL3352 | CTAGAGGAAACAAACTATAGGAAACGACCAGGCCCTCAACATCTCTCACCCATCTCCACACGGGGGTTACCCGGCCTGCA | Non-template DNA strand for the G-less cassette/G-rich tract transcription template. |
| DL3353 | GGCCGGGTAACCCCCGTGTGGAGATGGGTGAGAGATGTTGAGGGCCTGGTCGTTTCCTATAGTTTGTTTCCT | Template DNA strand for the G-less cassette/G-rich tract transcription template. |
| DL4604 | CTAGAGGAAACAAACTATAGGAAACGACCAGGCCCTCAACATCTCTCACCCATCTCCACACTTTTTTTTTGGACCCGGCC | Non-template DNA strand for the pause-inducing T-tract transcription template. |
| DL4607 | GGCCGGGTCCAAAAAAAAAGTGTGGAGATGGGTGAGAGATGTTGAGGGCCTGGTCGTTTCCTATAGTTTGTTTCCT | Template DNA strand for the pause-inducing T-tract transcription template. |
| pRMS754 | AGCCACATCATCTGCTGCACC | Non-phosphorylated forward primer for generating the SETX prong-deleted backbone used to construct SETX-HD$^{Sen1prong}$. |
| pRMS755 | GGAGAACTTCAGGACCTCAGA | Non-phosphorylated reverse primer for generating the SETX prong-deleted backbone used to construct SETX-HDS$^{en1prong}$. |
| pRMS756 | AGCCACATCATCTGCTGCACC | 5′-phosphorylated forward primer for generating an internal deletion within SETX-HD (Δ2061-2147; SETX-HDΔprong). |
| pRMS757 | GGAGAACTTCAGGACCTCAGA | 5′-phosphorylated reverse primer for generating an internal deletion within SETX-HD (Δ2061-2147; SETX-HDΔprong). |
| pRMS758 | TTAGAAGAACTTGTGGATAAAAG | 5′-phosphorylated forward primer for amplifying the *S. cerevisiae* Sen1 prong insert (residues 1461-1554) used to generate SETX-HD$^{Sen1prong}$. |
| pRMS759 | AACCGCCAAGATATGGGCCTG | 5′-phosphorylated reverse primer for amplifying the *S. cerevisiae* Sen1 prong insert (residues 1461-1554) used to generate SETX-HD$^{Sen1prong}$. |
| pRMS804 | UUUUUUGGCAUAGUU | 15-mer RNA primer fluorescently labelled on 5′ end with Cy3 used as a single-stranded RNA for EMSAs assays. |
| pRMS833 | TACTTCCAATCCAATGCATGCCATTCGGAGAATCTAAG | Forward primer for generating a 1674-2677 amino acid fragment of SETX for cloning into 2L-T plasmid. |
| pRMS834 | TTATCCACTTCCAATGTTATTAATGGTGATGGTGATGGTGATG | Reverse primer for generating a 1674-2677 amino acid fragment of SETX for cloning into 2L-T plasmid. |
| pRMS835 | TACTTCCAATCCAATCGATGGACTACAAAGACGATGA | Forward primer for generating a FLAG-tagged 1674-2677 amino acid fragment of SETX for cloning into 438-A plasmid. |

| Name | Sequence 5′-3′ | Use |
|---|---|---|
| pRMS795 | TACTTCCAATCCAATGCATACAGGCACGACGC TGTCAAG | Forward primer for generating SETX C terminus (2455-2677) in 2B-T plasmid. |
| pRMS840 | TTATCCACTTCCAATGTTATTACAGCAACTTG CGCTTCTTGGA | Reverse primer for generating SETX C-terminus (2455-2677) in 2B-T plasmid. |
| pRMS893 | GGATTGGAAGTAGAGGTTCTCGGGCTTGAGC TTGAGGAT | Reverse primer for generating SETX-HD$^{\Delta CT}$ (residues 1674-2467) for cloning into 2Bc-T plasmid. |
| pRMS937 | UAAGAGCAAGAUGUUCUAUAAAAGAUGUCC UAGCAAGGCAC | 41-mer RNA primer fluorescently labelled on 5′ end with FITC used as single-stranded substrate for EMSAs assays. |
| pRMS948 | GTCCTTGCTGGTGACGGTCAA | 5′-phosphorylated reverse primer for generating Δ2527-2560 in SETX-HD (residues 1674-2677) and SETX-CT (residues 2455-2677), and Δ2527-2590 in SETX-CT. |
| pRMS949 | TCATCTCCACAGCACCCTGGT | 5′-phosphorylated forward primer for generating Δ2527-2560 in SETX-HD (residues 1674-2677) and SETX-CT (residues 2455-2677). |
| pRMS950 | GCTGCTGTAGTCGCTGCCCTC | 5′-phosphorylated forward primer for generating Δ2527-2590 in SETX-CT (residues 2455-2677). |
| pRMS951 | GCCATGTCTCTGTTGGGATCTGCTAACCAGCT GGCTAGGGCCGTC | Forward primer for site-directed mutagenesis to introduce R1899A mutation. |
| pRMS952 | GACGGCCCTAGCCAGCTGGTTAGCAGATCCC AACAGAGACATGGC | Reverse primer for site-directed mutagenesis to introduce R1899A mutation. |
| pRMS953 | TCTCGCAACCAGCTGGCTGCTGCCGTCTTGAA CCCCAATCCTATG | Forward primer for site-directed mutagenesis to introduce R1904A mutation. |
| pRMS954 | CATAGGATTGGGGTTCAAGACGGCAGCAGCC AGCTGGTTGCGAGA | Reverse primer for site-directed mutagenesis to introduce R1904A mutation. |
| pRMS955 | CAAGAGCTGGCCTCCAAGATCGCAGAGGTCC AGGGACGCCCTCAG | Forward primer for site-directed mutagenesis to introduce K2131A mutation. |
| pRMS956 | CTGAGGGCGTCCCTGGACCTCTGCGATCTTGG AGGCCAGCTCTTG | Reverse primer for site-directed mutagenesis to introduce K2131A mutation. |
| pRMS957 | AAGATCAAAGAGGTCCAGGGAGCCCCTCAGA AAACCCAGTCTATC | Forward primer for site-directed mutagenesis to introduce R2136A mutation. |
| pRMS958 | GATAGACTGGGTTTTCTGAGGGGCTCCCTGGA CCTCTTTGATCTT | Reverse primer for site-directed mutagenesis to introduce R2136A mutation. |
| pRMS959 | CTCCTGCTGGAATCCGCATTCGCTGGACAAGG CGGAGTCCCCTTC | Forward primer for site-directed mutagenesis to introduce R2168A mutation. |
| pRMS960 | GAAGGGGACTCCGCCTTGTCCAGCGAATGCG GATTCCAGCAGGAG | Reverse primer for site-directed mutagenesis to introduce R2168A mutation. |

| Name | Sequence 5′-3′ | Use |
|---|---|---|
| pRMS961 | ACTCTGACCCCACTGATCCACGCGTGCAACAAGCTCATCCTCGTG | Forward primer for site-directed mutagenesis to introduce R2198A mutation. |
| pRMS962 | CACGAGGATGAGCTTGTTGCACGCGTGGATCAGTGGGGTCAGAGT | Reverse primer for site-directed mutagenesis to introduce R2198A mutation. |
| pRMS965 | GAAAGGCAAGAGCTGGCCTCCGCGATCGCAGAGGTCCAGGGACGCCCTCAG | Forward primer for site-directed mutagenesis to introduce K2129A/K2131A substitutions. |
| pRMS966 | CTGAGGGCGTCCCTGGACCTCTGCGATTGCGGAGGCCAGCTCTTGCCTTTC | Reverse primer for site-directed mutagenesis to introduce K2129A/K2131A substitutions. |
| pRMS967 | GAGGTCCAGGGACGCCCTCAGGCAACCCAGTCTATCATTATCCTC | Forward primer for site-directed mutagenesis to introduce K2139A mutation. |
| pRMS968 | GAGGATAATGATAGACTGGGTTGCCTGAGGGCGTCCCTGGACCTC | Reverse primer for site-directed mutagenesis to introduce K2139A mutation. |
| pRMS969 | TCCATGATGGCCAGGTTCTGCGCCCTGCTCGAAGAAAACGTCGAA | Forward primer for site-directed mutagenesis to introduce R2235A mutation. |
| pRMS970 | TTCGACGTTTTCTTCGAGCAGGGCGCAGAACCTGGCCATCATGGA | Reverse primer for site-directed mutagenesis to introduce R2235A mutation. |

**Supplementary Table S3. List of plasmids used in *in vitro* work**

| Name | Addgene # | Plasmid | Tags / Fusion Proteins | Bacterial Resistance(s) |
|---|---|---|---|---|
| 2B-T | 29666 | pET His6 TEV LIC cloning vector | His6-TEV (N-terminal on backbone) | Ampicillin |
| 2Bc-T | 37236 | pET His6 LIC cloning vector | His6-TEV (C-terminal on backbone) | Ampicillin |
| 2L-T | 29715 | pET Flag TEV LIC cloning vector | FLAG-TEV (N-terminal on backbone) | Ampicillin |
| 438-A | 55218 | pFastBac cloning vector with BioBrick PolyPromoter LIC Subcloning | No tag | Ampicillin |

**Supplementary Table S4. List of recombinant plasmids used in *in vitro* work**

| Name | Backbone | ORF(s) | Boundaries | Source |
|---|---|---|---|---|
| pMS5-1 | 438-A | FLAG-SETX-HD-$(His)_6$ | 1674-2677 | this study |
| pMS5-2 | 438-A | SETX-HD$^{\Delta CT}$-$(His)_6$ | 1674-2467 | this study |
| pMS5-3 | 438-A | SETX-HD$^{\Delta CT,D2181A/E2182A}$-$(His)_6$ | 1674-2467 | this study |
| pMS5-4 | 438-A | FLAG-SETX-HD$^{\Delta CT\text{-}RBD}$-$(His)_6$ | 1674-2677$^{\Delta 2527\text{-}2560}$ | this study |
| pMS5-5 | 2B-T | $(His)_6$-SETX-CT | 2455-2677 | this study |
| pMS5-6 | 2B-T | $(His)_6$-SETX-CT$^{\Delta CT\text{-}RBD}$ | 2455-2677$^{\Delta 2527\text{-}2560}$ | this study |
| pMS5-7 | 2B-T | $(His)_6$-SETX-CT$^{\Delta 2527\text{-}2590}$ | 2455-2677$^{\Delta 2527\text{-}2590}$ | this study |
| pMS5-8 | 438-A | FLAG-SETX-HD$^{Sen1prong}$-$(His)_6$ | 1674-2677; SETX 2061-2147 replaced by *Sc*Sen1 1461-1554 | this study |
| pMS5-9 | 438-A | FLAG-SETX-HD$^{R1899A}$-$(His)_6$ | 1674-2677 | this study |
| pMS5-10 | 438-A | FLAG-SETX-HD$^{R1904A}$-$(His)_6$ | 1674-2677 | this study |
| pMS5-11 | 438-A | FLAG-SETX-HD$^{K2129A/K2131A}$-$(His)_6$ | 1674-2677 | this study |
| pMS5-12 | 438-A | FLAG-SETX-HD$^{K2131A}$-$(His)_6$ | 1674-2677 | this study |
| pMS5-13 | 438-A | FLAG-SETX-HD$^{R2136A}$-$(His)_6$ | 1674-2677 | this study |
| pMS5-14 | 438-A | FLAG-SETX-HD$^{K2139A}$-$(His)_6$ | 1674-2677 | this study |
| pMS5-15 | 438-A | FLAG-SETX-HD$^{R2168A}$-$(His)_6$ | 1674-2677 | this study |
| pMS5-16 | 438-A | FLAG-SETX-HD$^{R2198A}$-$(His)_6$ | 1674-2677 | this study |
| pMS5-17 | 438-A | FLAG-SETX-HD$^{R2235A}$-$(His)_6$ | 1674-2677 | this study |
| pMS5-18 | 438-A | FLAG-SETX-HD$^{\Delta prong}$-$(His)_6$ | 1674-2677$^{\Delta 2061\text{-}2147}$ | this study |

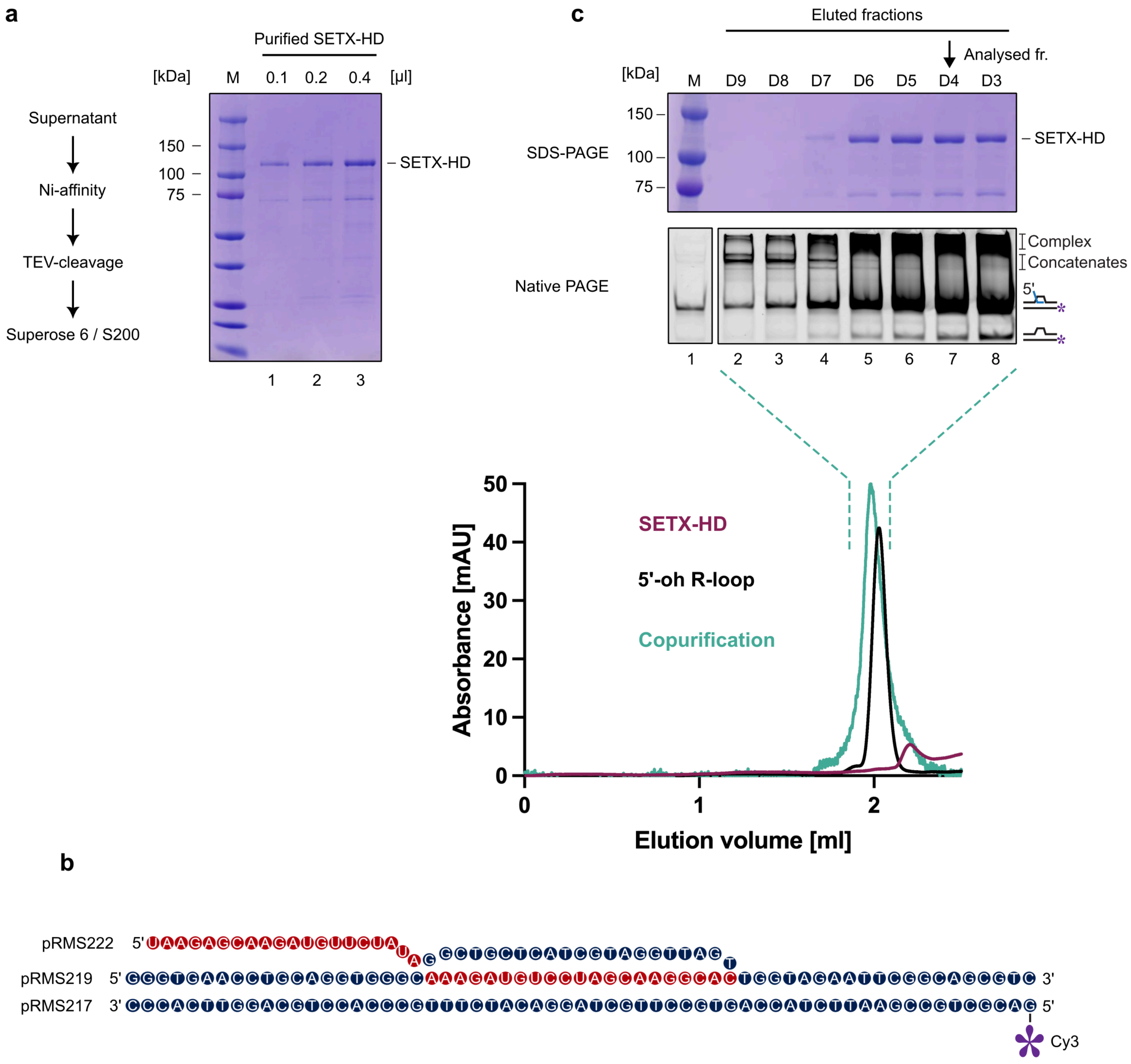
a
Purified SETX-HD
[kDa]
M
0.1
0.2
0.4
[µl]
Supernatant
Ni-affinity
TEV-cleavage
Superose 6 / S200
150
100
75
SETX-HD
1
2
3
c
Eluted fractions
Analysed fr.
[kDa]
M
D9
D8
D7
D6
D5
D4
D3
SDS-PAGE
150
100
75
SETX-HD
Native PAGE
Complex
Concatenates
5'
1
2
3
4
5
6
7
8
50
40
30
20
10
0
Absorbance [mAU]
SETX-HD
5'-oh R-loop
Copurification
0
1
2
Elution volume [ml]
b
pRMS222 5' UAAGAGCAAGAUGUUCUAUAG GCTGCTCATCGTAGGTTAGT
pRMS219 5' GGGTGAACCTGCAGGTGGGCAAAGAUGUCCUAGCAAGGCACTGGTAGAATTCGGCAGCGTC 3'
pRMS217 3' CCCACTTTGGACGTCCACCCGTTTCTACAGGATCGTTCCGTGACCATCTTAAGCCGTCGCAG 5'
Cy3

Supplementary figure 2

Lanicek *et al.*

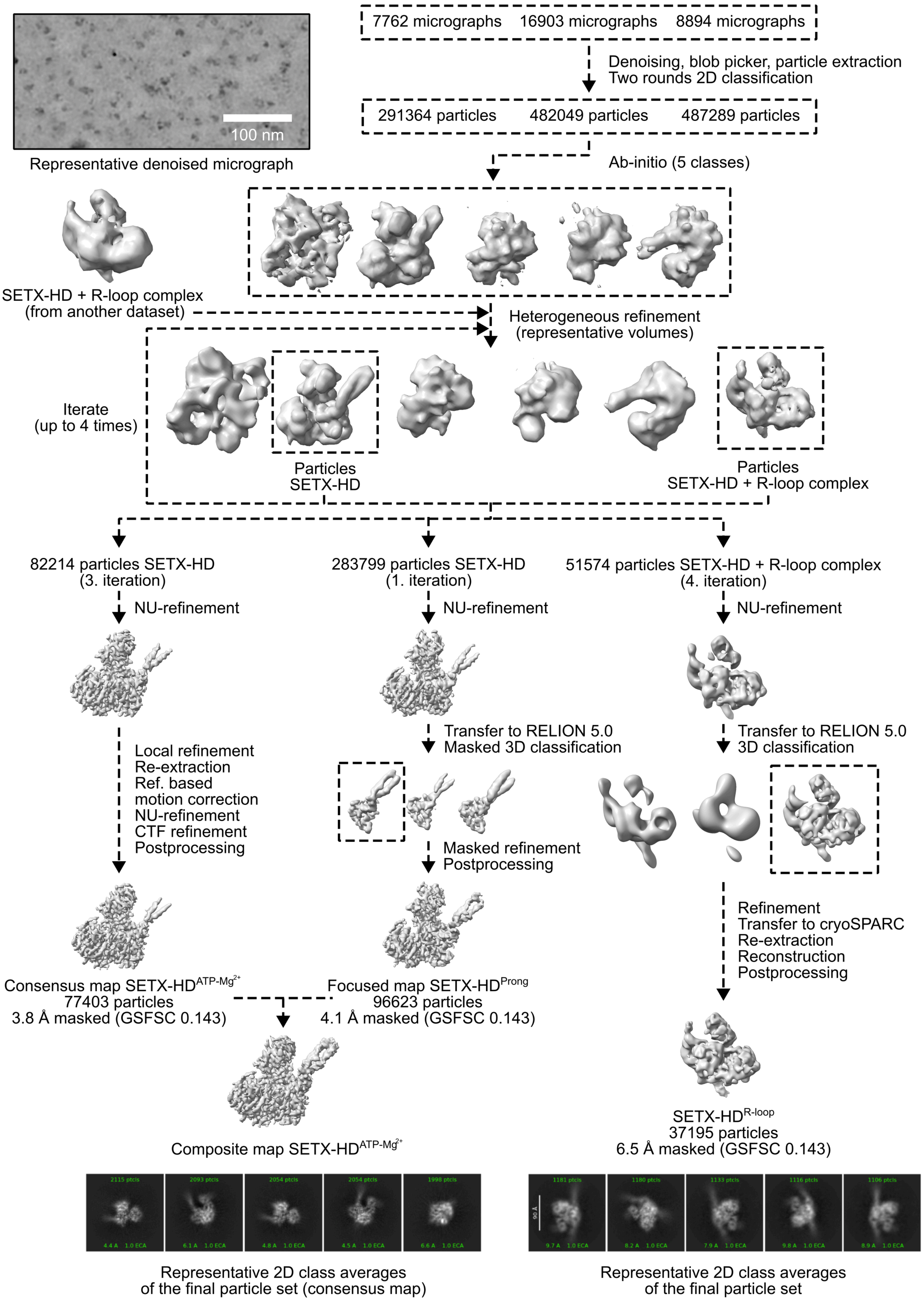

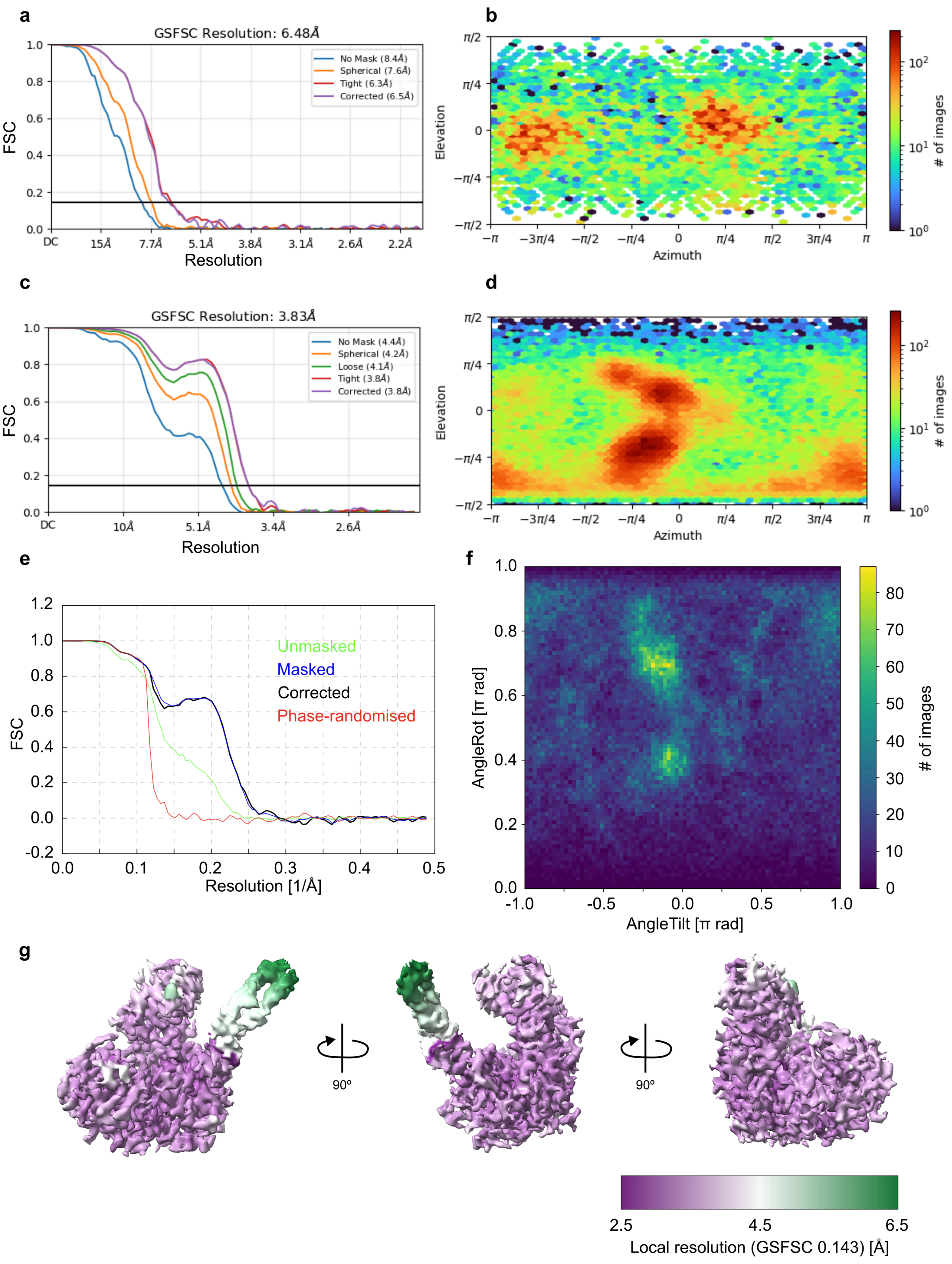
a
GSFSC Resolution: 6.48Å
No Mask (8.4Å)
Spherical (7.6Å)
Tight (6.3Å)
Corrected (6.5Å)
FSC
Resolution
DC
15Å
7.7Å
5.1Å
3.8Å
3.1Å
2.6Å
2.2Å
b
Elevation
Azimuth
### of images
c
GSFSC Resolution: 3.83Å
No Mask (4.4Å)
Spherical (4.2Å)
Loose (4.1Å)
Tight (3.8Å)
Corrected (3.8Å)
FSC
Resolution
DC
10Å
5.1Å
3.4Å
2.6Å
d
Elevation
Azimuth
### of images
e
Unmasked
Masked
Corrected
Phase-randomised
FSC
Resolution [1/Å]
f
AngleRot [π rad]
AngleTilt [π rad]
### of images
g
90º
90º
2.5
4.5
6.5
Local resolution (GSFSC 0.143) [Å]

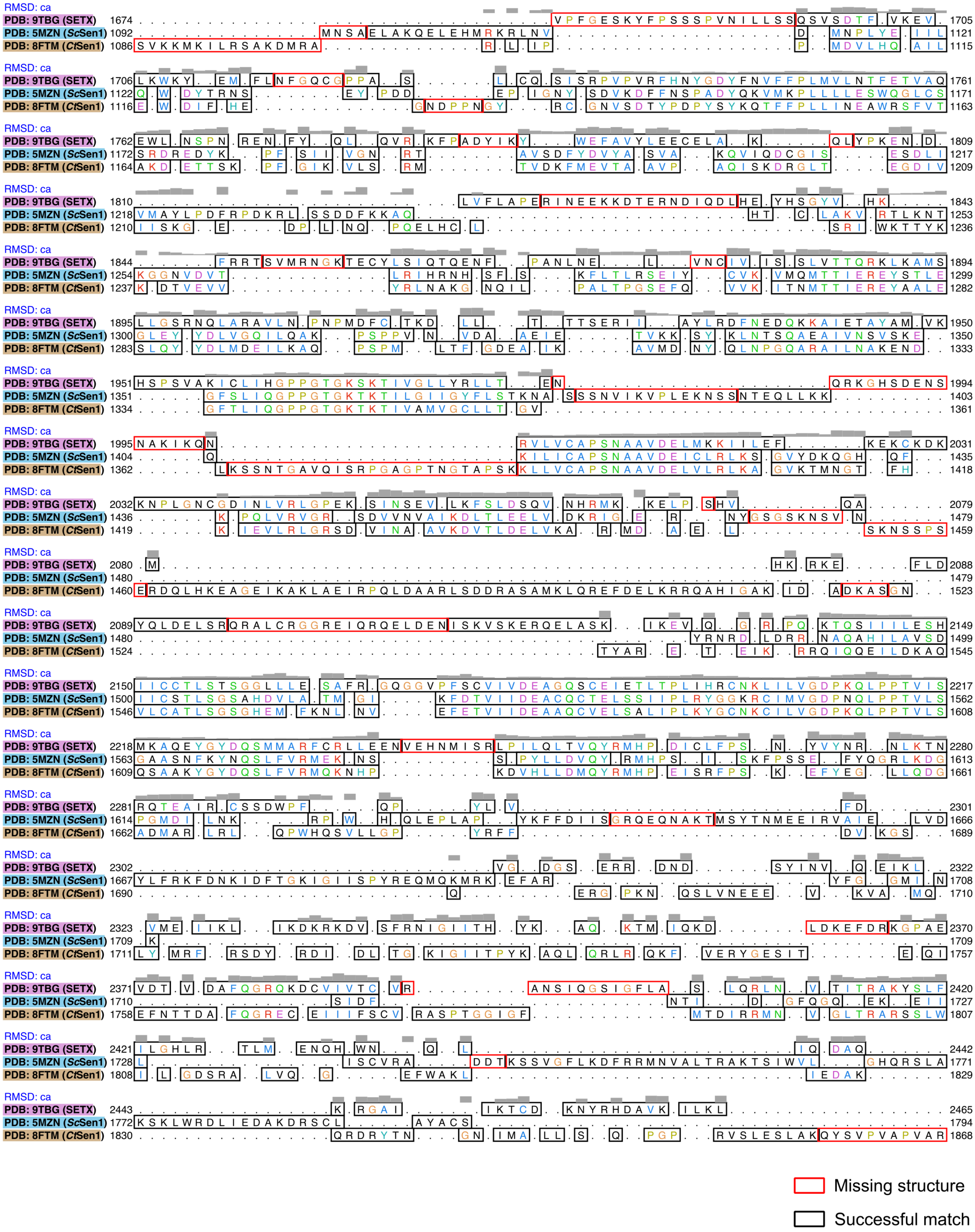
RMSD: ca
PDB: 9TBG (SETX)
PDB: 5MZN (ScSen1)
PDB: 8FTM (CtSen1)
Missing structure
Successful match

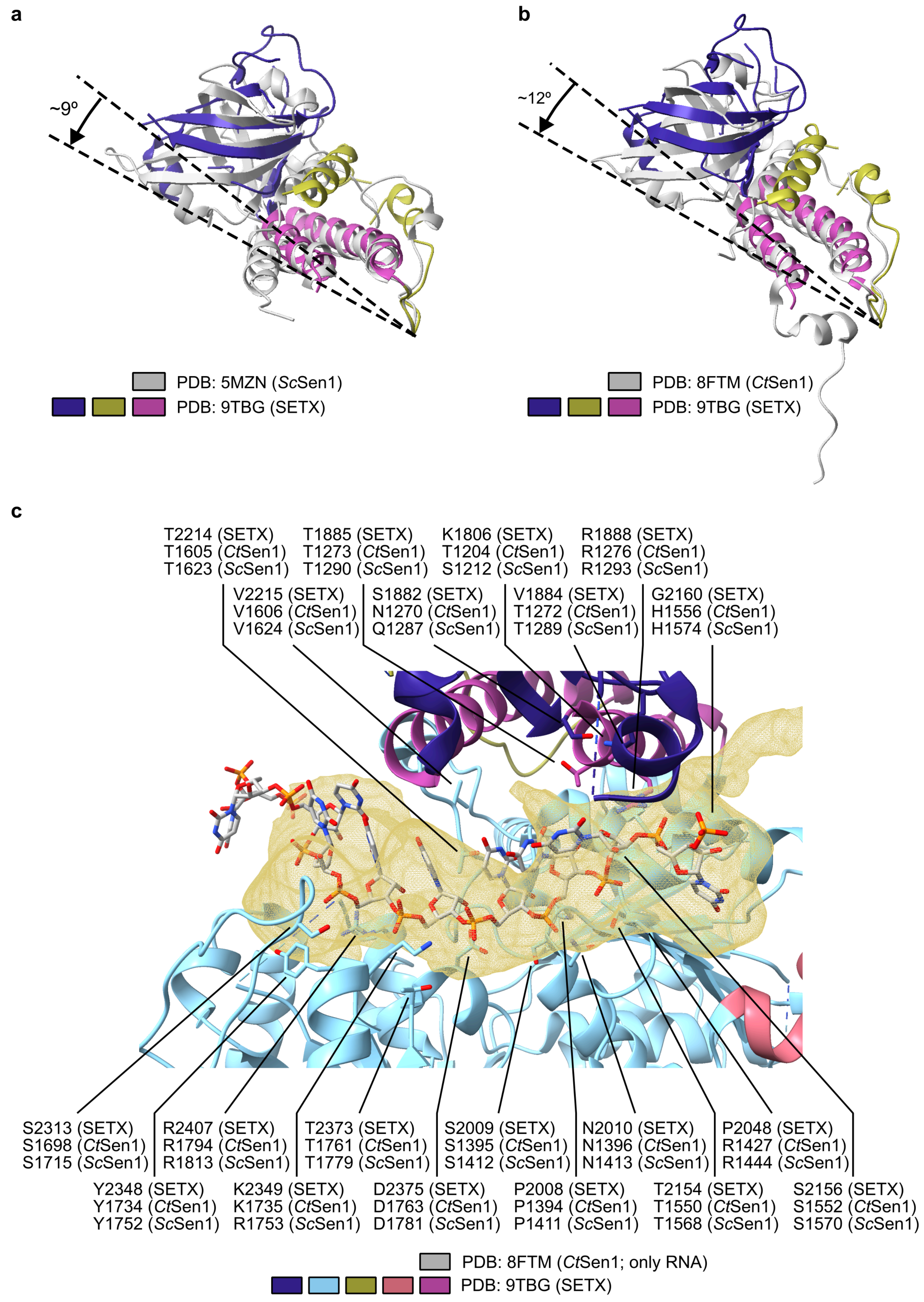
a
~9º
PDB: 5MZN (ScSen1)
PDB: 9TBG (SETX)
b
~12º
PDB: 8FTM (CtSen1)
PDB: 9TBG (SETX)
c
T2214 (SETX) T1605 (CtSen1) T1623 (ScSen1)
T1885 (SETX) T1273 (CtSen1) T1290 (ScSen1)
K1806 (SETX) T1204 (CtSen1) S1212 (ScSen1)
R1888 (SETX) R1276 (CtSen1) R1293 (ScSen1)
V2215 (SETX) V1606 (CtSen1) V1624 (ScSen1)
S1882 (SETX) N1270 (CtSen1) Q1287 (ScSen1)
V1884 (SETX) T1272 (CtSen1) T1289 (ScSen1)
G2160 (SETX) H1556 (CtSen1) H1574 (ScSen1)
S2313 (SETX) S1698 (CtSen1) S1715 (ScSen1)
R2407 (SETX) R1794 (CtSen1) R1813 (ScSen1)
T2373 (SETX) T1761 (CtSen1) T1779 (ScSen1)
S2009 (SETX) S1395 (CtSen1) S1412 (ScSen1)
N2010 (SETX) N1396 (CtSen1) N1413 (ScSen1)
P2048 (SETX) R1427 (CtSen1) R1444 (ScSen1)
Y2348 (SETX) Y1734 (CtSen1) Y1752 (ScSen1)
K2349 (SETX) K1735 (CtSen1) R1753 (ScSen1)
D2375 (SETX) D1763 (CtSen1) D1781 (ScSen1)
P2008 (SETX) P1394 (CtSen1) P1411 (ScSen1)
T2154 (SETX) T1550 (CtSen1) T1568 (ScSen1)
S2156 (SETX) S1552 (CtSen1) S1570 (ScSen1)
PDB: 8FTM (CtSen1; only RNA)
PDB: 9TBG (SETX)

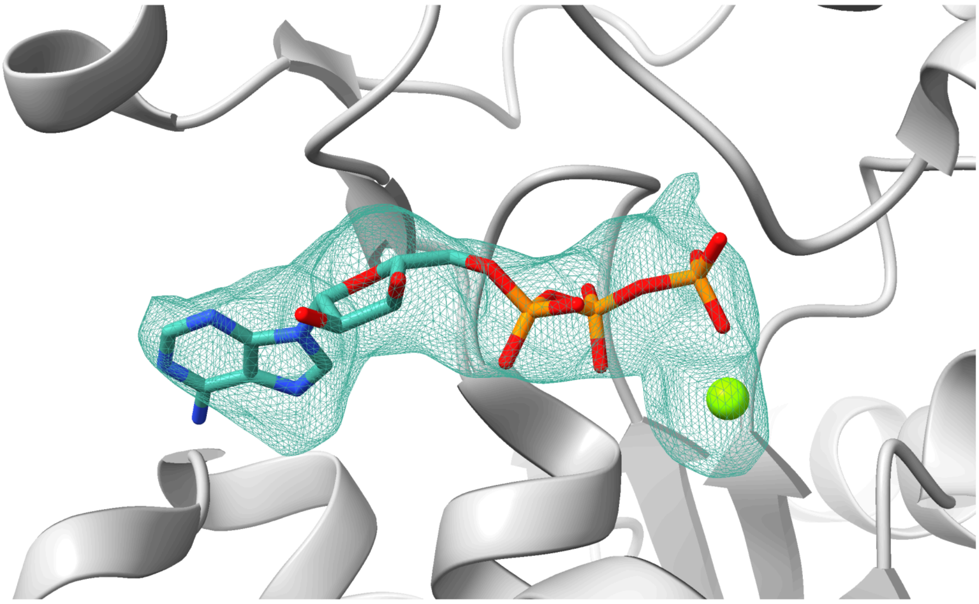

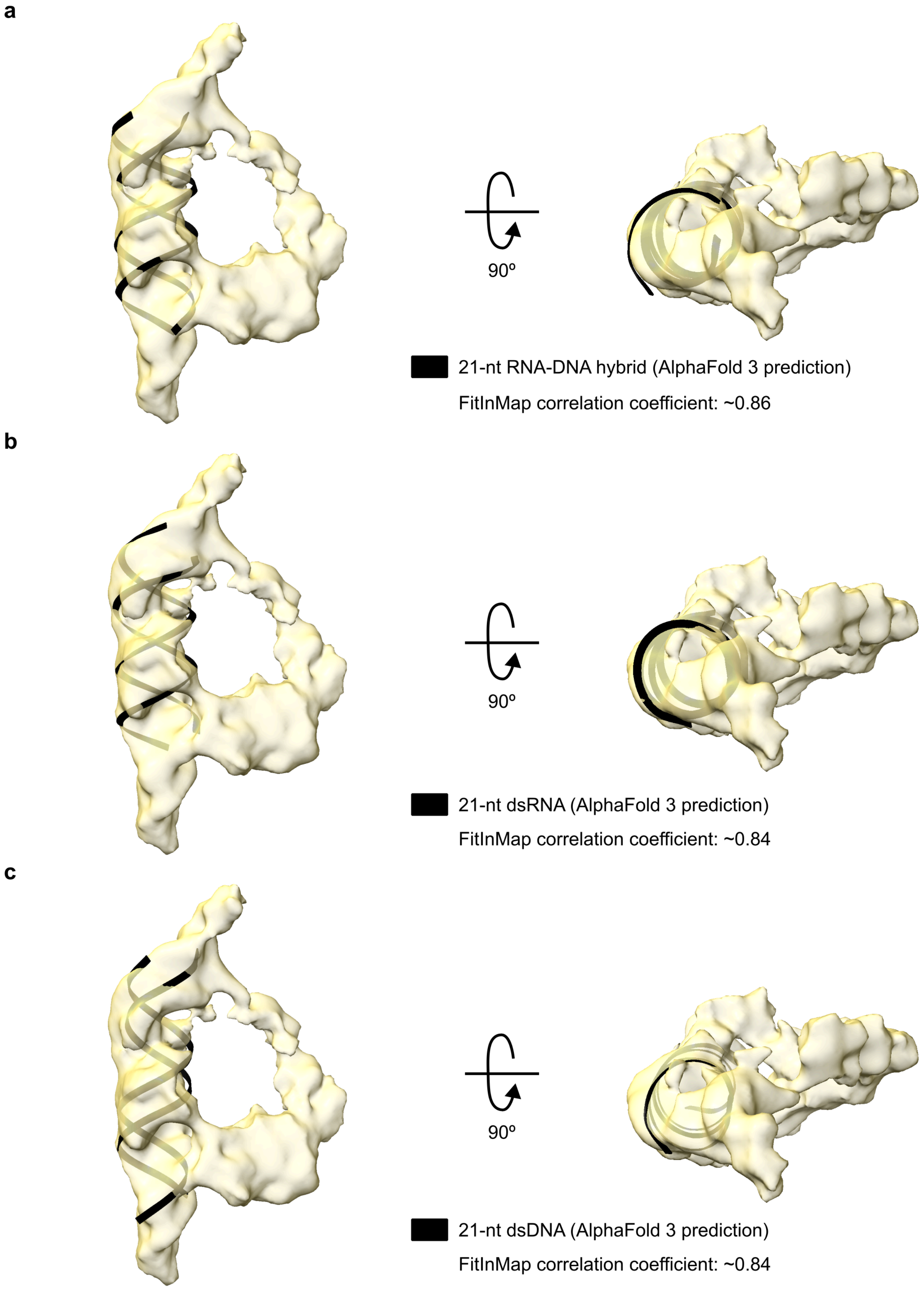
a
90º
21-nt RNA-DNA hybrid (AlphaFold 3 prediction)
FitInMap correlation coefficient: ~0.86
b
90º
21-nt dsRNA (AlphaFold 3 prediction)
FitInMap correlation coefficient: ~0.84
c
90º
21-nt dsDNA (AlphaFold 3 prediction)
FitInMap correlation coefficient: ~0.84

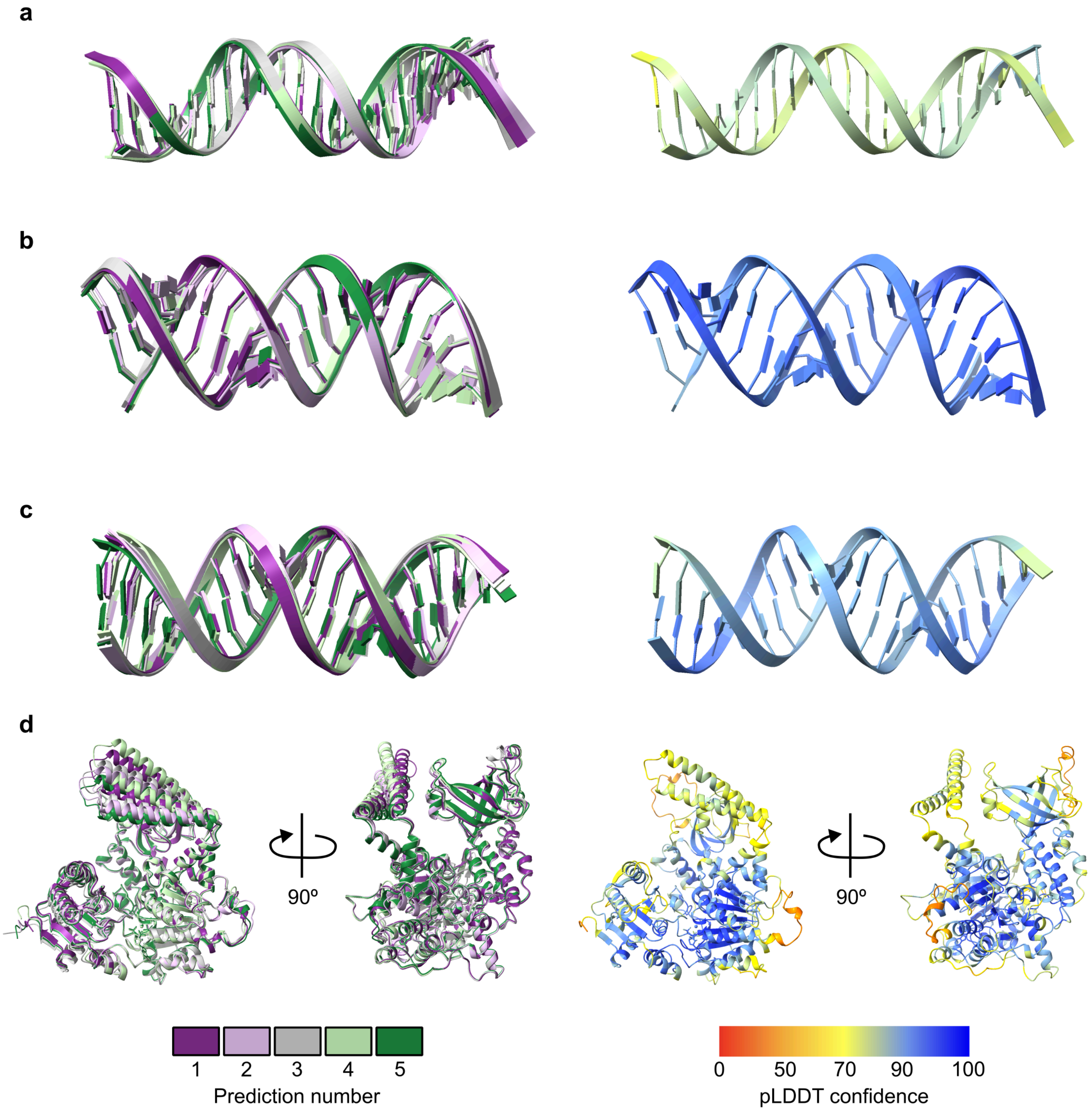
a
b
c
d
90º
90º
1
2
3
4
5
Prediction number
0
50
70
90
100
pLDDT confidence

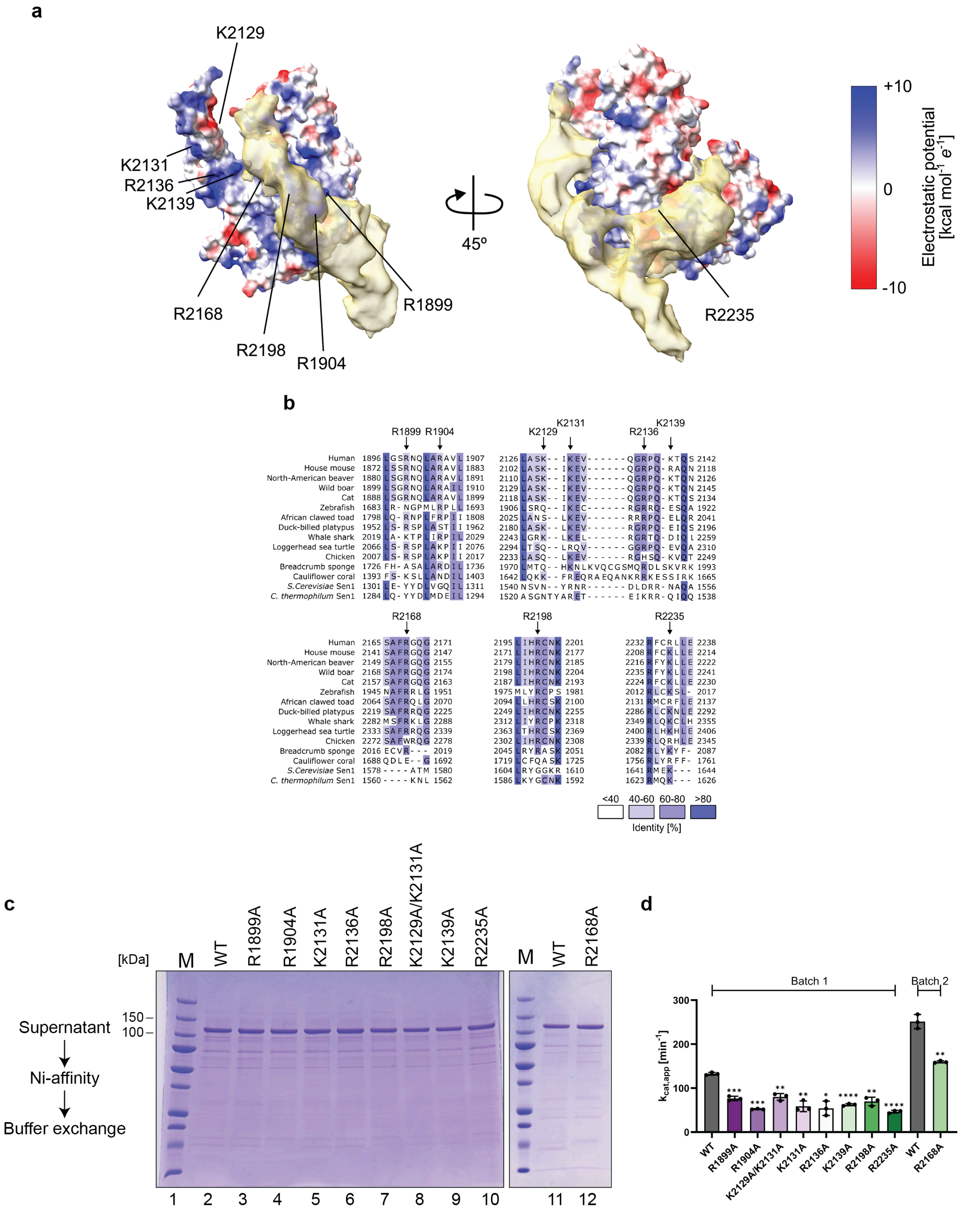
a
K2129
K2131
R2136
K2139
R2168
R2198
R1904
R1899
45º
R2235
+10
0
-10
Electrostatic potential [kcal mol-1 e-1]
b
R1899 R1904
K2129 K2131 R2136 K2139
Human 1896 LGSRNQLARAVL 1907 2126 LASK--IKEV------QGRPQ-KTQS 2142
House mouse 1872 LSSRNQLARAVL 1883 2102 LASK--IKEV------QGRPQ-RAQN 2118
North-American beaver 1880 LSGRNQLARAVL 1891 2110 LASK--IKEV------QGRPQ-KTQN 2126
Wild boar 1899 LSGRNQLARAIL 1910 2129 LASK--IKEV------QGRPQ-KTQN 2145
Cat 1888 LSGRNQLARAVL 1899 2118 LASK--IKEV------QGRPQ-KTQS 2134
Zebrafish 1683 LR-NGPMLRPLL 1693 1906 LSRQ--IKEC------RGRRQ-ESQA 1922
African clawed toad 1798 LQ-RNPLFRPII 1808 2025 LANS--LKEV------RRRPQ-ELQR 2041
Duck-billed platypus 1952 LS-RSPLASTII 1962 2180 LASK--LKEV------RGRPQ-EIQS 2196
Whale shark 2019 LA-KTPLIRPIL 2029 2243 LGRK--LKEL------RGRTQ-DIQL 2259
Loggerhead sea turtle 2066 LS-RSPLAKPII 2076 2294 LTSQ--LRQV------GGRPQ-EVQA 2310
Chicken 2007 LS-RSPLAKPII 2017 2233 LASQ--LKEV------RGHSQ-KVQT 2249
Breadcrumb sponge 1726 FH-ASALARDIL 1736 1970 LMTQ--HKNLKVQCGSMQRDLSKVRK 1993
Cauliflower coral 1393 FS-KSLLANDIL 1403 1642 LQKK--FREQRAEQANKRRKESSIRK 1665
S.Cerevisiae Sen1 1301 LE-YYDLVGQIL 1311 1540 NSVN--YRNR------DLDRR-NAQA 1556
C. thermophilum Sen1 1284 LQ-YYDLMDEIL 1294 1520 ASGNTYARET------EIKRR-QIQQ 1538
R2168 R2198 R2235
Human 2165 SAFRGQG 2171 2195 LIHRCNK 2201 2232 RFCRLLE 2238
House mouse 2141 SAFRGQG 2147 2171 LIHRCNK 2177 2208 RFCKLLE 2214
North-American beaver 2149 SAFRGQG 2155 2179 LIHRCNK 2185 2216 RFYKLLE 2222
Wild boar 2168 SAFRGQG 2174 2198 LIHRCNK 2204 2235 RFYKLLE 2241
Cat 2157 SAFRGQG 2163 2187 LIHRCNK 2193 2224 RFCKLLE 2230
Zebrafish 1945 NAFRRLG 1951 1975 MLYRCPS 1981 2012 RLCKSL- 2017
African clawed toad 2064 SAFRQLG 2070 2094 LLHRCSK 2100 2131 RMCRFLE 2137
Duck-billed platypus 2219 SAFRRQG 2225 2249 LIHRCNK 2255 2286 RLCKNLE 2292
Whale shark 2282 MSFRKLG 2288 2312 LIYRCPK 2318 2349 RLQKCLH 2355
Loggerhead sea turtle 2333 SAFRRQG 2339 2363 LTHRCSK 2369 2400 RLHKHLE 2406
Chicken 2272 SAFWRQG 2278 2302 LIHRCNK 2308 2339 RLQRHLE 2345
Breadcrumb sponge 2016 ECVR--- 2019 2045 LRYRASK 2051 2082 RLYKYF- 2087
Cauliflower coral 1688 QDLE--G 1692 1719 LCFQASK 1725 1756 RLYRFF- 1761
S.Cerevisiae Sen1 1578 ----ATM 1580 1604 LRYGGKR 1610 1641 RMEK--- 1644
C. thermophilum Sen1 1560 ----KNL 1562 1586 LKYGCNK 1592 1623 RMQK--- 1626
<40 40-60 60-80 >80
Identity [%]
c
[kDa]
150 –
100 –
M WT R1899A R1904A K2131A R2136A R2198A K2129A/K2131A K2139A R2235A
M WT R2168A
Supernatant
Ni-affinity
Buffer exchange
1 2 3 4 5 6 7 8 9 10
11 12
d
Batch 1
Batch 2
300
200
100
0
kcat,app [min-1]
***
***
**
**
*
****
**
****
**
WT
R1899A
R1904A
K2129A/K2131A
K2131A
R2136A
K2139A
R2198A
R2235A
WT
R2168A

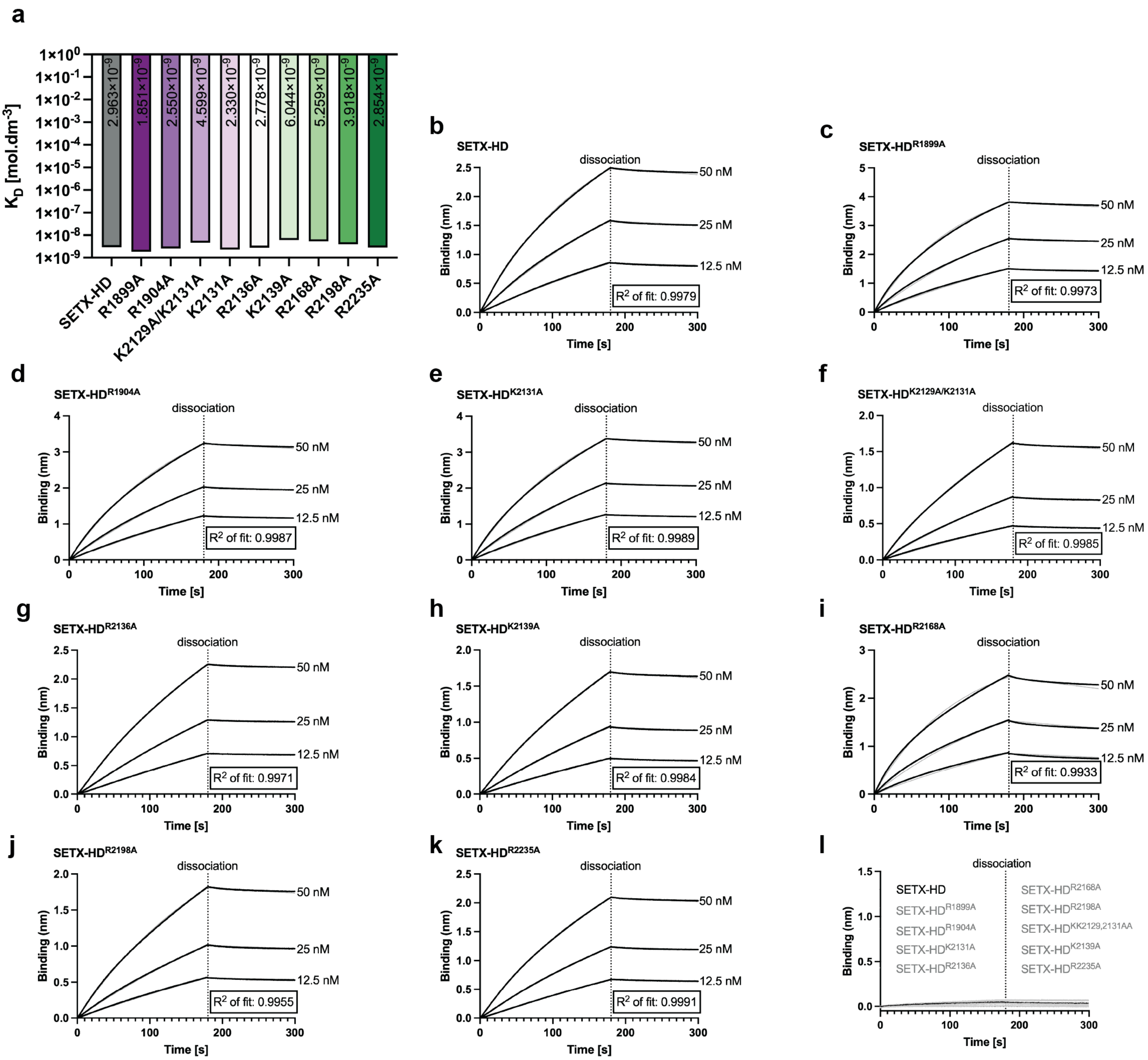
a
K_D [mol.dm-3]
2.963×10-9
1.851×10-9
2.550×10-9
4.599×10-9
2.330×10-9
2.778×10-9
6.044×10-9
5.259×10-9
3.918×10-9
2.854×10-9
SETX-HD
R1899A
R1904A
K2129A/K2131A
K2131A
R2136A
K2139A
R2168A
R2198A
R2235A
b
SETX-HD
R2 of fit: 0.9979
c
SETX-HDR1899A
R2 of fit: 0.9973
d
SETX-HDR1904A
R2 of fit: 0.9987
e
SETX-HDK2131A
R2 of fit: 0.9989
f
SETX-HDK2129A/K2131A
R2 of fit: 0.9985
g
SETX-HDR2136A
R2 of fit: 0.9971
h
SETX-HDK2139A
R2 of fit: 0.9984
i
SETX-HDR2168A
R2 of fit: 0.9933
j
SETX-HDR2198A
R2 of fit: 0.9955
k
SETX-HDR2235A
R2 of fit: 0.9991
l
SETX-HD
SETX-HDR1899A
SETX-HDR1904A
SETX-HDK2131A
SETX-HDR2136A
SETX-HDR2168A
SETX-HDR2198A
SETX-HDKK2129,2131AA
SETX-HDK2139A
SETX-HDR2235A
dissociation
50 nM
25 nM
12.5 nM
Binding (nm)
Time [s]

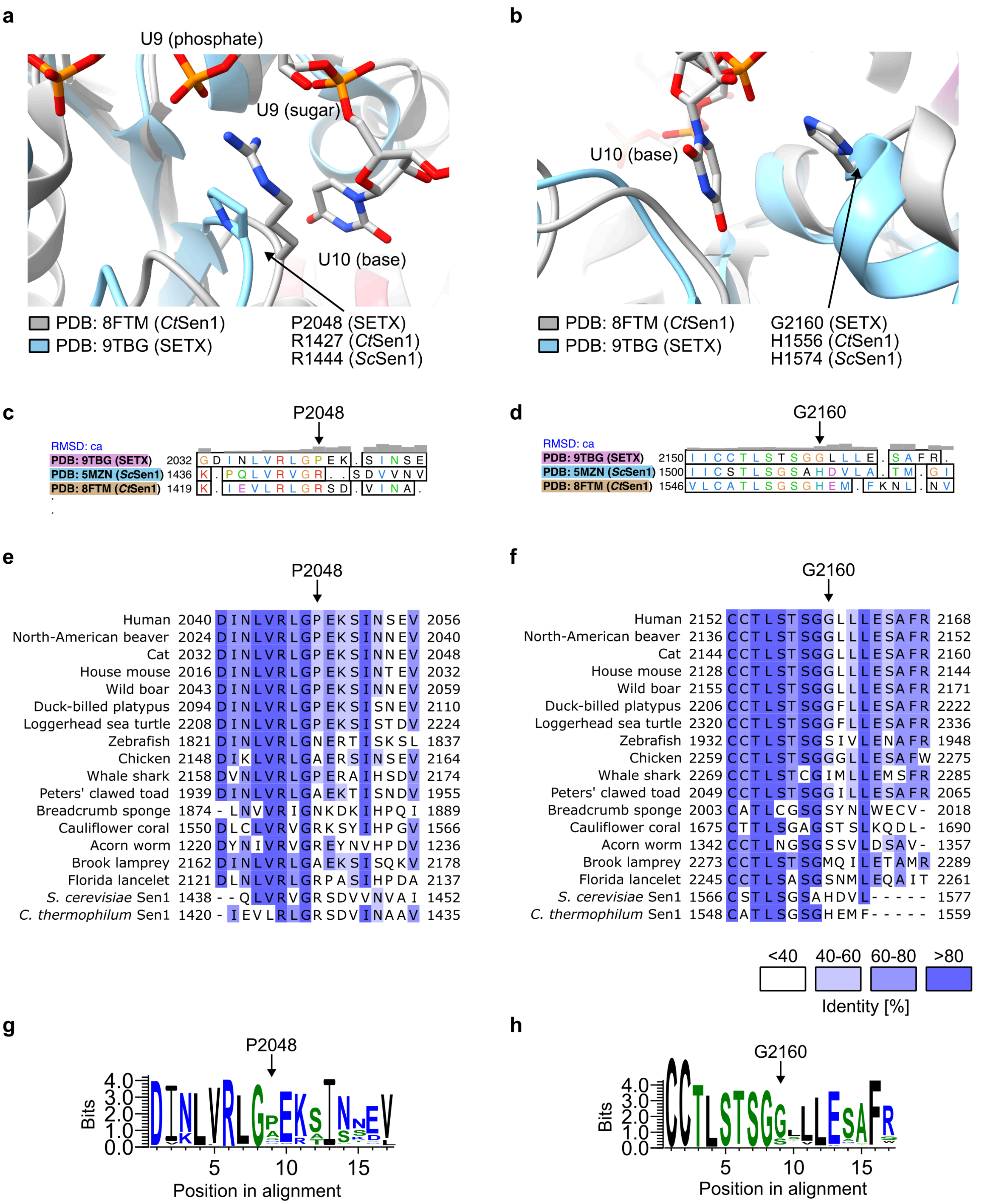
a
U9 (phosphate)
U9 (sugar)
U10 (base)
PDB: 8FTM (CtSen1)
PDB: 9TBG (SETX)
P2048 (SETX)
R1427 (CtSen1)
R1444 (ScSen1)
b
U10 (base)
PDB: 8FTM (CtSen1)
PDB: 9TBG (SETX)
G2160 (SETX)
H1556 (CtSen1)
H1574 (ScSen1)
c
P2048
RMSD: ca
PDB: 9TBG (SETX) 2032 GDINLVRLGPEK.SINSE
PDB: 5MZN (ScSen1) 1436 K.PQLVRVGR..SDVVNV
PDB: 8FTM (CtSen1) 1419 K.IEVLRLGRSD.VINA.
d
G2160
RMSD: ca
PDB: 9TBG (SETX) 2150 IICCTLSTSGGLLLE.SAFR.
PDB: 5MZN (ScSen1) 1500 IICSTLSGSAHDVLA.TM.GI
PDB: 8FTM (CtSen1) 1546 VLCATLSGSGHEM.FKNL.NV
e
P2048
Human 2040 DINLVRLGPEKSINSEV 2056
North-American beaver 2024 DINLVRLGPEKSINNEV 2040
Cat 2032 DINLVRLGPEKSINNEV 2048
House mouse 2016 DINLVRLGPEKSINTEV 2032
Wild boar 2043 DINLVRLGPEKSINNEV 2059
Duck-billed platypus 2094 DINLVRLGPEKSISNEV 2110
Loggerhead sea turtle 2208 DINLVRLGPEKSISTDV 2224
Zebrafish 1821 DINLVRLGNERTISKSL 1837
Chicken 2148 DIKLVRLGAERSINSEV 2164
Whale shark 2158 DVNLVRLGPERAIHSDV 2174
Peters' clawed toad 1939 DINLVRLGAEKTISNDV 1955
Breadcrumb sponge 1874 -LNVVRIGNKDKIHPQI 1889
Cauliflower coral 1550 DLCLVRVGRKSYIHPGV 1566
Acorn worm 1220 DYNIVRVGREYNVHPDV 1236
Brook lamprey 2162 DINLVRLGAEKSISQKV 2178
Florida lancelet 2121 DLNLVRLGRPASIHPDA 2137
S. cerevisiae Sen1 1438 --QLVRVGRSDVVNVAI 1452
C. thermophilum Sen1 1420 -IEVLRLGRSDVINAAV 1435
f
G2160
Human 2152 CCTLSTSGGLLLESAFR 2168
North-American beaver 2136 CCTLSTSGGLLLESAFR 2152
Cat 2144 CCTLSTSGGLLLESAFR 2160
House mouse 2128 CCTLSTSGGLLLESAFR 2144
Wild boar 2155 CCTLSTSGGLLLESAFR 2171
Duck-billed platypus 2206 CCTLSTSGGFLLESAFR 2222
Loggerhead sea turtle 2320 CCTLSTSGGFLLESAFR 2336
Zebrafish 1932 CCTLSTSGSIVLENAFR 1948
Chicken 2259 CCTLSTSGGGLLESAFW 2275
Whale shark 2269 CCTLSTCGIMLLEMSFR 2285
Peters' clawed toad 2049 CCTLSTSGGILLESAFR 2065
Breadcrumb sponge 2003 CATLCGSGSYNLWECV- 2018
Cauliflower coral 1675 CTTLSGAGSTSLKQDL- 1690
Acorn worm 1342 CCTLNGSGSSVLDSAV- 1357
Brook lamprey 2273 CCTLSTSGMQILETAMR 2289
Florida lancelet 2245 CCTLSASGSNMLEQAIT 2261
S. cerevisiae Sen1 1566 CSTLSGSAHDVL----- 1577
C. thermophilum Sen1 1548 CATLSGSGHEMF----- 1559
<40
40-60
60-80
>80
Identity [%]
g
P2048
Bits
Position in alignment
h
G2160
Bits
Position in alignment

**a**

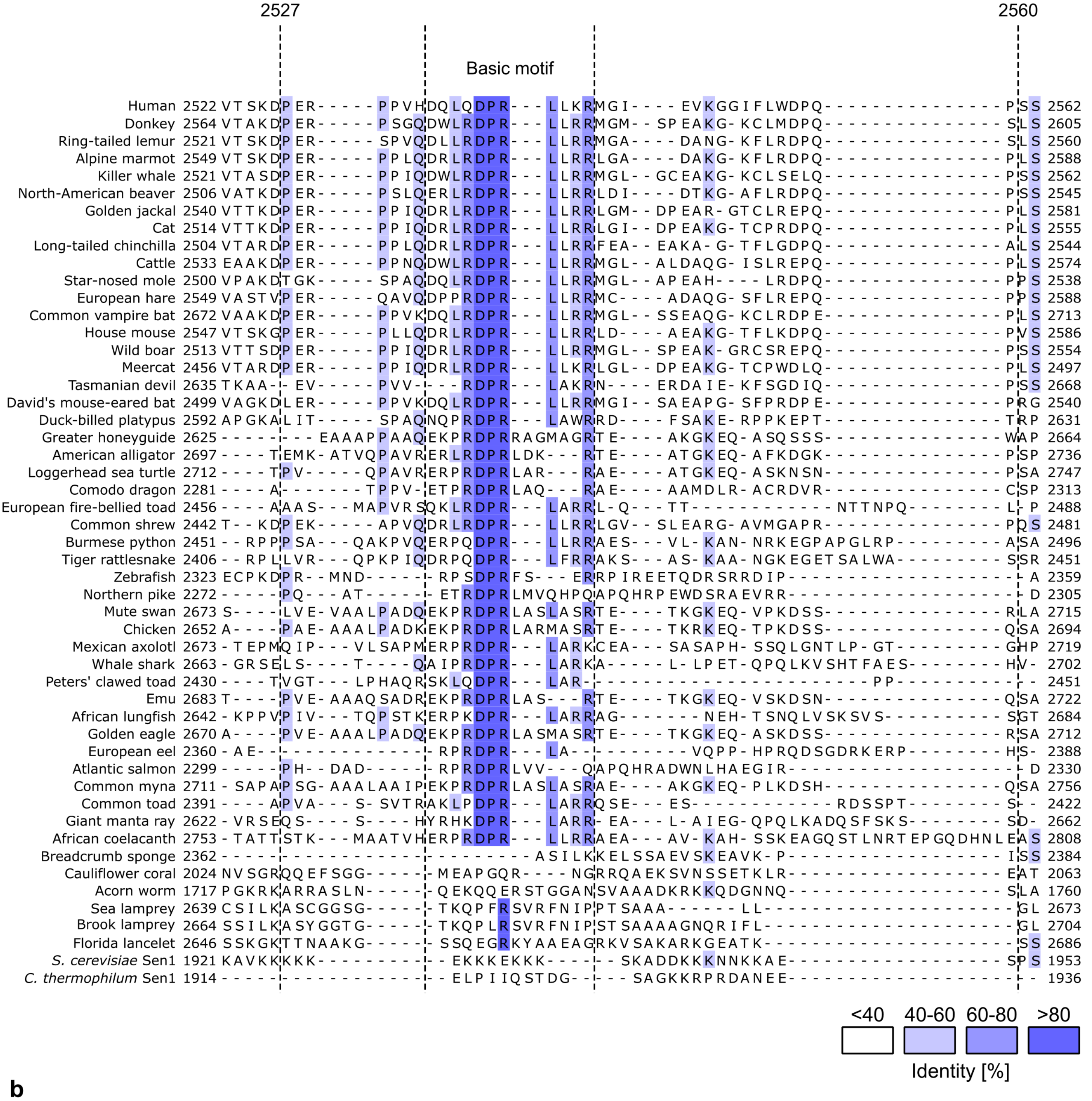
2527
2560
Basic motif
<40
40-60
60-80
>80
Identity [%]

**b**

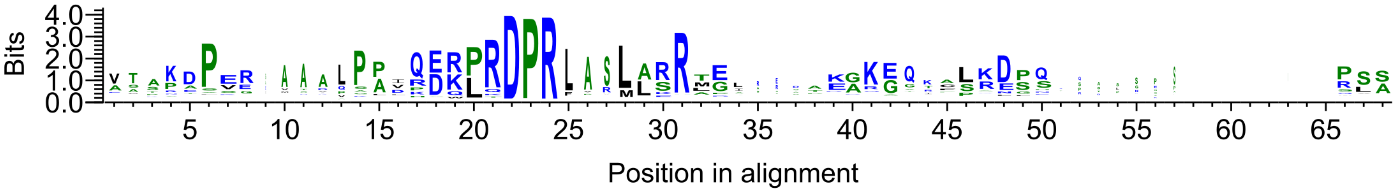
Bits
Position in alignment

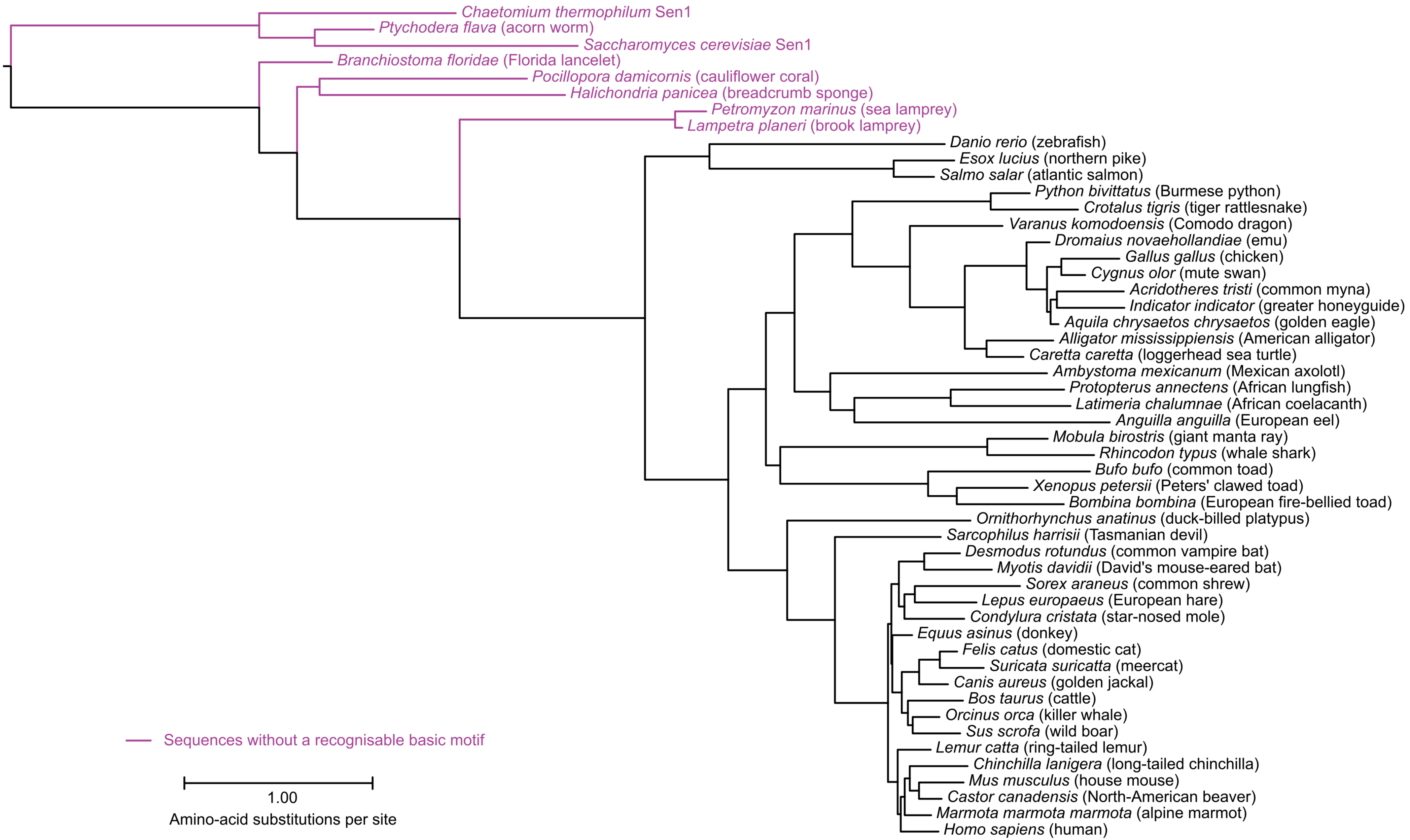
Chaetomium thermophilum Sen1
Ptychodera flava (acorn worm)
Saccharomyces cerevisiae Sen1
Branchiostoma floridae (Florida lancelet)
Pocillopora damicornis (cauliflower coral)
Halichondria panicea (breadcrumb sponge)
Petromyzon marinus (sea lamprey)
Lampetra planeri (brook lamprey)
Danio rerio (zebrafish)
Esox lucius (northern pike)
Salmo salar (atlantic salmon)
Python bivittatus (Burmese python)
Crotalus tigris (tiger rattlesnake)
Varanus komodoensis (Comodo dragon)
Dromaius novaehollandiae (emu)
Gallus gallus (chicken)
Cygnus olor (mute swan)
Acridotheres tristi (common myna)
Indicator indicator (greater honeyguide)
Aquila chrysaetos chrysaetos (golden eagle)
Alligator mississippiensis (American alligator)
Caretta caretta (loggerhead sea turtle)
Ambystoma mexicanum (Mexican axolotl)
Protopterus annectens (African lungfish)
Latimeria chalumnae (African coelacanth)
Anguilla anguilla (European eel)
Mobula birostris (giant manta ray)
Rhincodon typus (whale shark)
Bufo bufo (common toad)
Xenopus petersii (Peters' clawed toad)
Bombina bombina (European fire-bellied toad)
Ornithorhynchus anatinus (duck-billed platypus)
Sarcophilus harrisii (Tasmanian devil)
Desmodus rotundus (common vampire bat)
Myotis davidii (David's mouse-eared bat)
Sorex araneus (common shrew)
Lepus europaeus (European hare)
Condylura cristata (star-nosed mole)
Equus asinus (donkey)
Felis catus (domestic cat)
Suricata suricatta (meercat)
Canis aureus (golden jackal)
Bos taurus (cattle)
Orcinus orca (killer whale)
Sus scrofa (wild boar)
Lemur catta (ring-tailed lemur)
Chinchilla lanigera (long-tailed chinchilla)
Mus musculus (house mouse)
Castor canadensis (North-American beaver)
Marmota marmota marmota (alpine marmot)
Homo sapiens (human)
Sequences without a recognisable basic motif
1.00
Amino-acid substitutions per site

Supplementary figure 14 Lanicek *et al.*

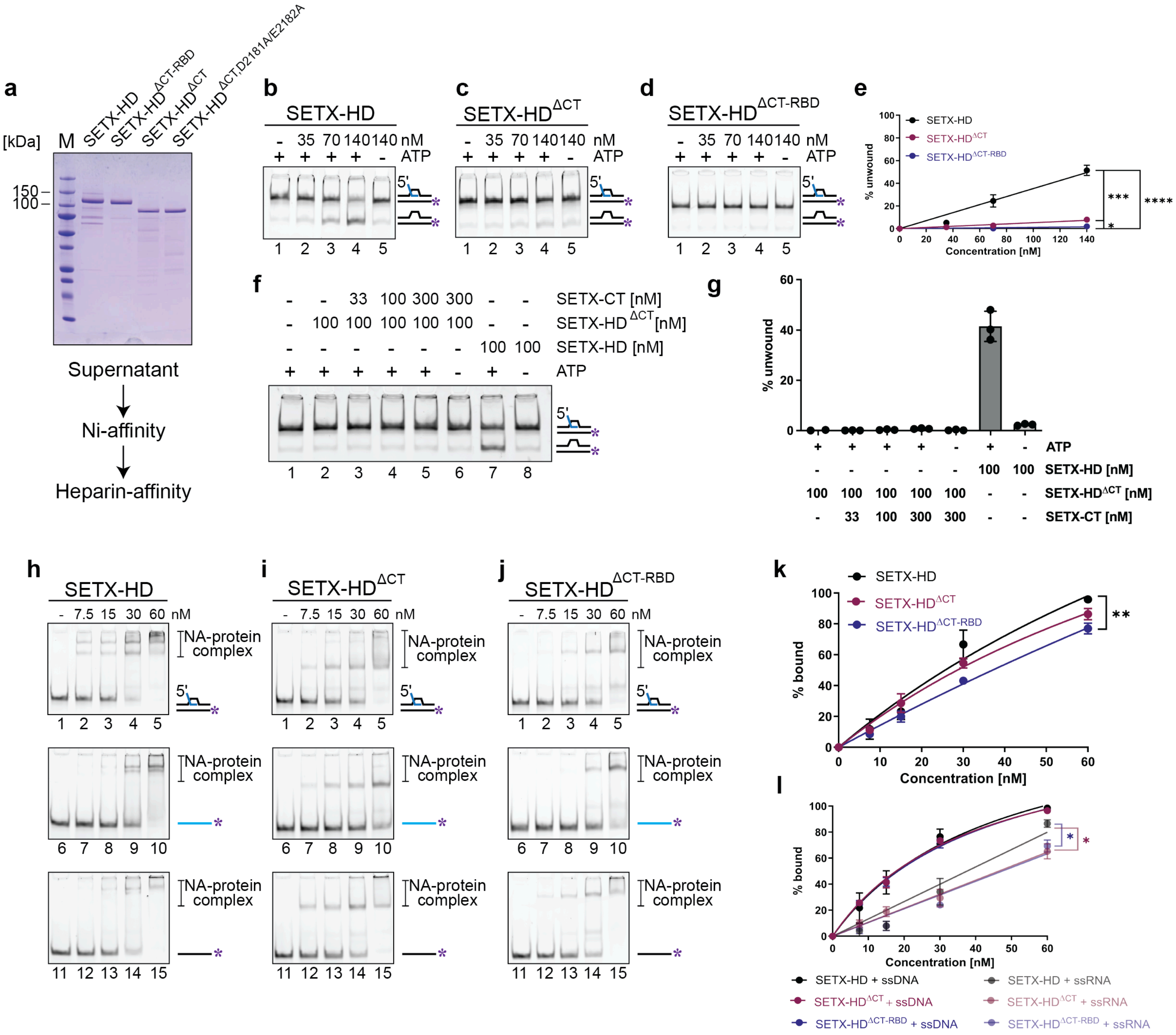

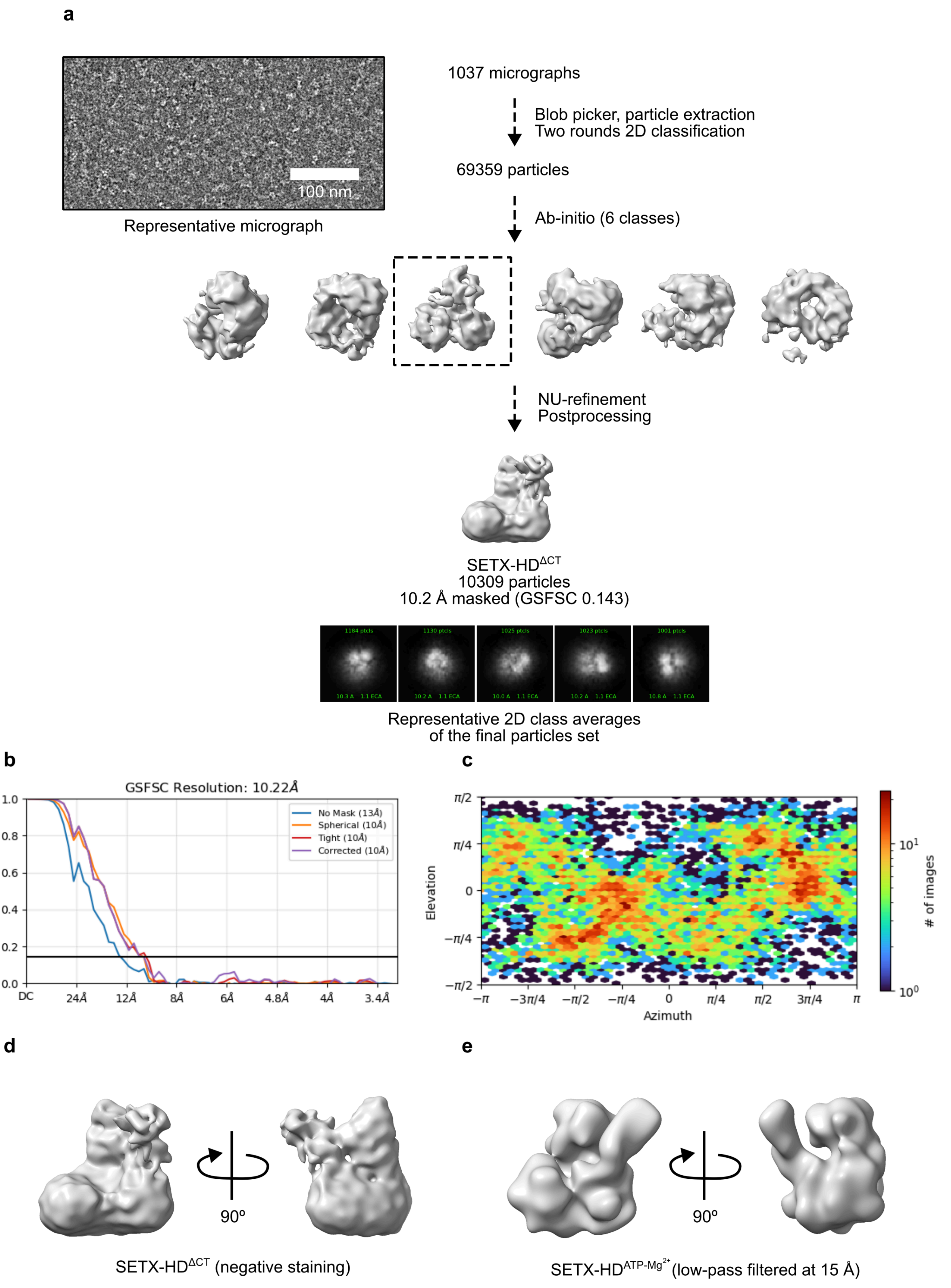
a
100 nm
Representative micrograph
1037 micrographs
Blob picker, particle extraction
Two rounds 2D classification
69359 particles
Ab-initio (6 classes)
NU-refinement
Postprocessing
SETX-HD$^{\Delta CT}$
10309 particles
10.2 Å masked (GSFSC 0.143)
Representative 2D class averages
of the final particles set
b
GSFSC Resolution: 10.22Å
No Mask (13Å)
Spherical (10Å)
Tight (10Å)
Corrected (10Å)
1.0
0.8
0.6
0.4
0.2
0.0
DC
24Å
12Å
8Å
6Å
4.8Å
4Å
3.4Å
c
Elevation
π/2
π/4
0
−π/4
−π/2
Azimuth
−π
−3π/4
−π/2
−π/4
0
π/4
π/2
3π/4
π
### of images
$10^1$
$10^0$
d
90º
SETX-HD$^{\Delta CT}$ (negative staining)
e
90º
SETX-HD$^{ATP\text{-}Mg^{2+}}$ (low-pass filtered at 15 Å)

Supplementary figure 16 Lanicek *et al.*

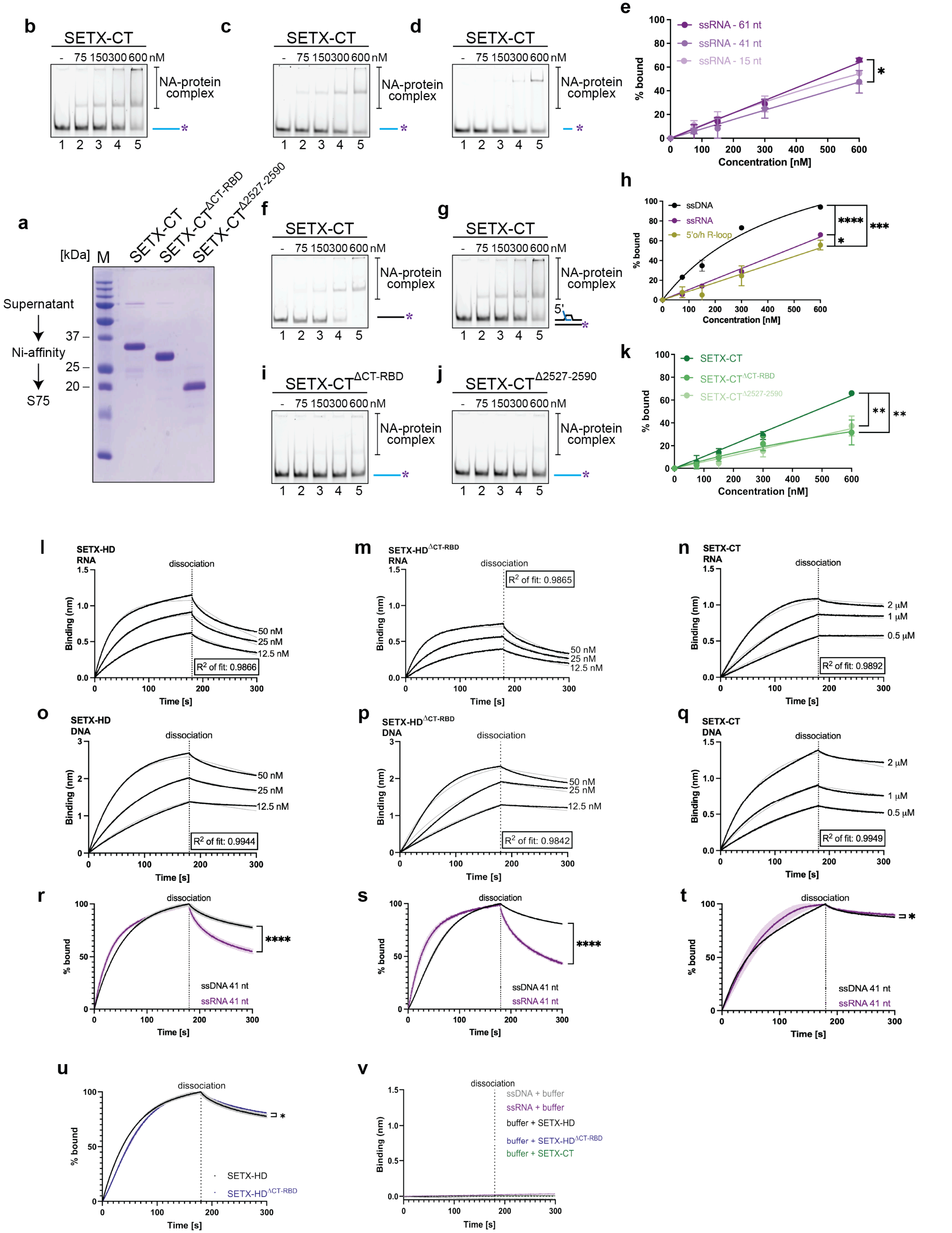

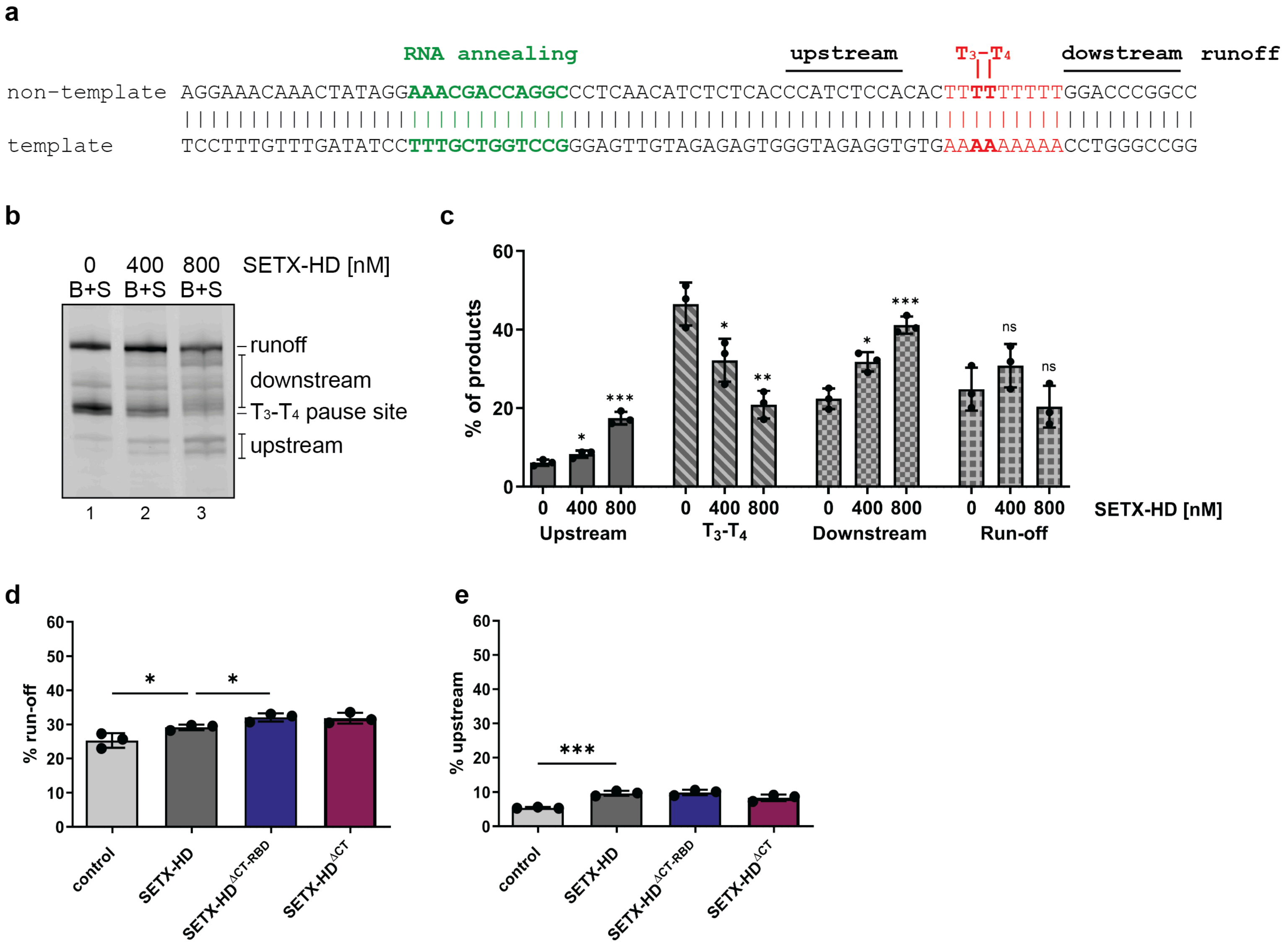
a
RNA annealing
upstream
T3–T4
dowstream
runoff
non-template AGGAAACAAACTATAGGAAACGACCAGGCCCTCAACATCTCTCACCCATCTCCACACTTTTTTTTTGGACCCGGCC
template TCCTTTGTTTGATATCCTTTGCTGGTCCGGGAGTTGTAGAGAGTGGGTAGAGGTGTGAAAAAAAAACCTGGGCCGG
b
0 400 800 SETX-HD [nM]
B+S B+S B+S
runoff
downstream
T3-T4 pause site
upstream
1 2 3
c
% of products
0 400 800 0 400 800 0 400 800 0 400 800 SETX-HD [nM]
Upstream
T3-T4
Downstream
Run-off
ns
d
% run-off
control
SETX-HD
SETX-HDΔCT-RBD
SETX-HDΔCT
e
% upstream
control
SETX-HD
SETX-HDΔCT-RBD
SETX-HDΔCT

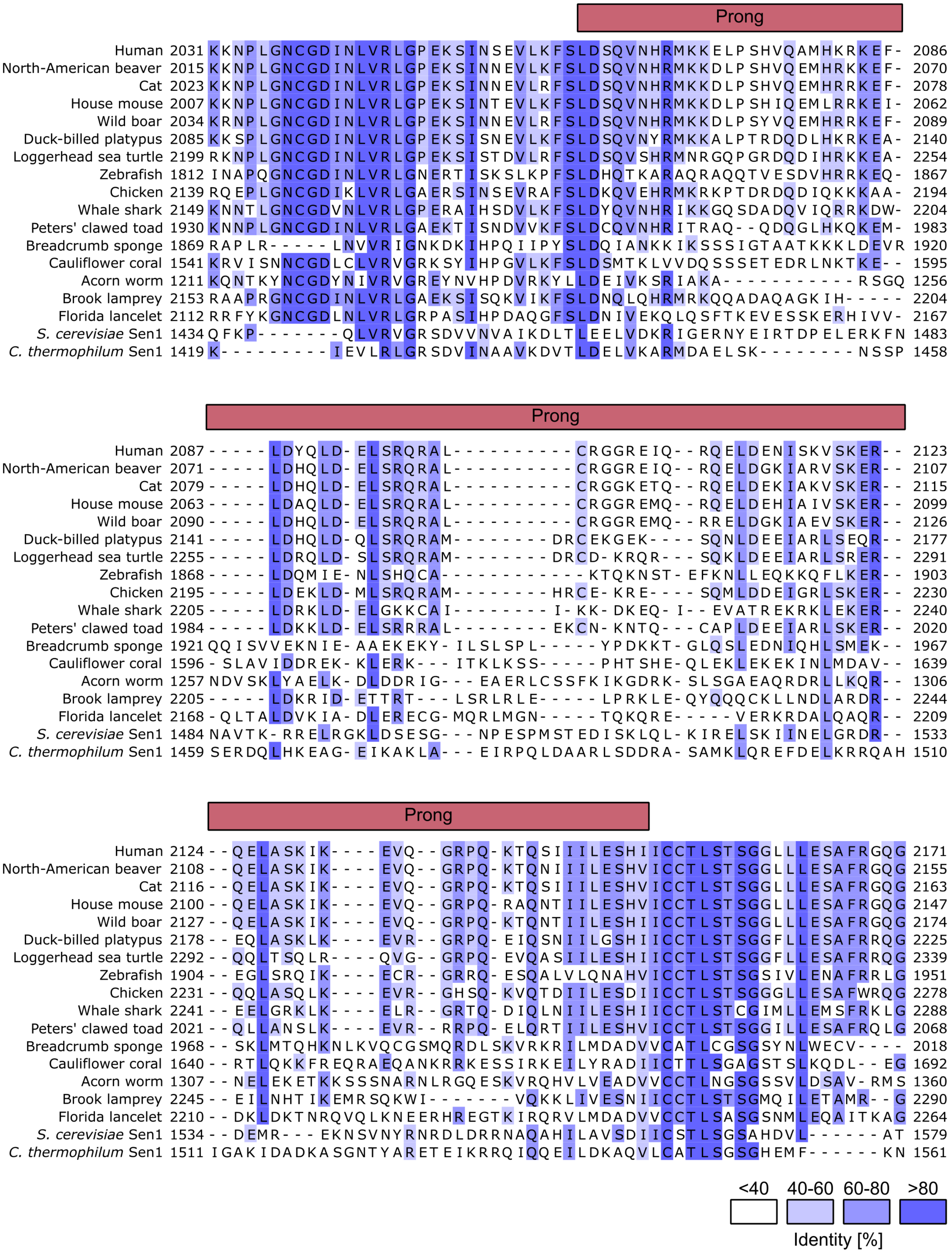
Prong
Human 2031 KKNPLGNCGDINLVRLGPEKSINSEVLKFSLDSQVNHRMKKELPSHVQAMHKRKEF- 2086
North-American beaver 2015 KKNPLGNCGDINLVRLGPEKSINNEVLKFSLDSQVNHRMKKDLPSHVQEMHRKKEF- 2070
Cat 2023 KKNPLGNCGDINLVRLGPEKSINNEVLRFSLDSQVNHRMKKDLPSHVQEMHRRKEF- 2078
House mouse 2007 KKNPLGNCGDINLVRLGPEKSINTEVLKFSLDSQVNHRMKKDLPSHIQEMLRRKEI- 2062
Wild boar 2034 KRNPLGNCGDINLVRLGPEKSINNEVLRFSLDSQVNHRMKKDLPSYVQEMHRRKEF- 2089
Duck-billed platypus 2085 KKSPLGNCGDINLVRLGPEKSISNEVLKFSLDSQVNYRMKKALPTRDQDLHKRKEA- 2140
Loggerhead sea turtle 2199 RKNPLGNCGDINLVRLGPEKSISTDVLRFSLDSQVSHRMNRGQPGRDQDIHRKKEA- 2254
Zebrafish 1812 INAPQGNCGDINLVRLGNERTISKSLKPFSLDHQTKARAQRAQQTVESDVHRRKEQ- 1867
Chicken 2139 RQEPLGNCGDIKLVRLGAERSINSEVRAFSLDKQVEHRMKRKPTDRDQDIQKKKAA- 2194
Whale shark 2149 KNNTLGNCGDVNLVRLGPERAIHSDVLKFSLDYQVNHRIKKGQSDADQVIQRRKDW- 2204
Peters' clawed toad 1930 KNNPLGNCGDINLVRLGAEKTISNDVVKFSLDCQVNHRITRAQ--QDQGLHKQKEM- 1983
Breadcrumb sponge 1869 RAPLR-----LNVVRIGNKDKIHPQIIPYSLDQIANKKIKSSSIGTAATKKKLDEVR 1920
Cauliflower coral 1541 KRVISNNCGDLCLVRVGRKSYIHPGVLKFSLDSMTKLVVDQSSSETEDRLNKTKE-- 1595
Acorn worm 1211 KQNTKYNCGDYNIVRVGREYNVHPDVRKYLLDEIVKSRIAKA-----------RSGQ 1256
Brook lamprey 2153 RAAPRGNCGDINLVRLGAEKSISQKVIKFSLDNQLQHRMRKQQADAQAGKIH----- 2204
Florida lancelet 2112 RRFYKGNCGDLNLVRLGRPASIHPDAQGFSLDNIVEKQLQSFTKEVESSKERHIVV- 2167
*S. cerevisiae* Sen1 1434 QFKP------QLVRVGRSDVVNVAIKDLTLEELVDKRIGERNYEIRTDPELERKFN 1483
*C. thermophilum* Sen1 1419 K---------IEVLRLGRSDVINAAVKDVTLDELVKARMDAELSK--------NSSP 1458
Prong
Human 2087 -----LDYQLD-ELSRQRAL----------CRGGREIQ--RQELDENISKVSKER-- 2123
North-American beaver 2071 -----LDHQLD-ELSRQRAL----------CRGGREIQ--RQELDGKIARVSKER-- 2107
Cat 2079 -----LDHQLD-ELSRQRAL----------CRGGKETQ--RQELDEKIAKVSKER-- 2115
House mouse 2063 -----LDAQLD-ELSRQRAL----------CRGGREMQ--RQELDEHIAIVSKER-- 2099
Wild boar 2090 -----LDHQLD-ELSRQRAL----------CRGGREMQ--RRELDGKIAEVSKER-- 2126
Duck-billed platypus 2141 -----LDHQLD-QLSRQRAM--------DRCEKGEK----SQNLDEEIARLSEQR-- 2177
Loggerhead sea turtle 2255 -----LDRQLD-SLSRQRAM--------DRCD-KRQR---SQKLDEEIARLSRER-- 2291
Zebrafish 1868 -----LDQMIE-NLSHQCA------------KTQKNST-EFKNLLEQKKQFLKER-- 1903
Chicken 2195 -----LDEKLD-MLSRQRAM--------HRCE-KRE----SQMLDDEIGRLSKER-- 2230
Whale shark 2205 -----LDRKLD-ELGKKCAI--------I-KK-DKEQ-I--EVATREKRKLEKER-- 2240
Peters' clawed toad 1984 -----LDKKLD-ELSRRRAL--------EKCN-KNTQ---CAPLDEEIARLSKER-- 2020
Breadcrumb sponge 1921 QQISVVEKNIE-AAEKEKY-ILSLSPL-----YPDKKT-GLQSLEDNIQHLSMEK-- 1967
Cauliflower coral 1596 -SLAVIDDREK-KLERK---ITKLKSS-----PHTSHE-QLEKLEKEKINLMDAV-- 1639
Acorn worm 1257 NDVSKLYAELK-DLDDRIG---EAERLCSSFKIKGDRK-SLSGAEAQRDRLLKQR-- 1306
Brook lamprey 2205 -----LDKRID-ETTRT---LSRLRLE-----LPRKLE-QYQQQCKLLNDLARDR-- 2244
Florida lancelet 2168 -QLTALDVKIA-DLERECG-MQRLMGN-----TQKQRE-----VERKRDALQAQR-- 2209
*S. cerevisiae* Sen1 1484 NAVTK-RRELRGKLDSESG---NPESPMSTEDISKLQL-KIRELSKIINELGRDR-- 1533
*C. thermophilum* Sen1 1459 SERDQLHKEAG-EIKAKLA---EIRPQLDAARLSDDRA-SAMKLQREFDELKRRQAH 1510
Prong
Human 2124 --QELASKIK----EVQ--GRPQ-KTQSIIILESHIICCTLSTSGGLLLESAFRGQG 2171
North-American beaver 2108 --QELASKIK----EVQ--GRPQ-KTQNIIILESHVICCTLSTSGGLLLESAFRGQG 2155
Cat 2116 --QELASKIK----EVQ--GRPQ-KTQSIIILESHVICCTLSTSGGLLLESAFRGQG 2163
House mouse 2100 --QELASKIK----EVQ--GRPQ-RAQNTIILESHVICCTLSTSGGLLLESAFRGQG 2147
Wild boar 2127 --QELASKIK----EVQ--GRPQ-KTQNTIILESHIICCTLSTSGGLLLESAFRGQG 2174
Duck-billed platypus 2178 --EQLASKLK----EVR--GRPQ-EIQSNIILGSHIICCTLSTSGGFLLESAFRRQG 2225
Loggerhead sea turtle 2292 --QQLTSQLR----QVG--GRPQ-EVQASIILESHIICCTLSTSGGFLLESAFRRQG 2339
Zebrafish 1904 --EGLSRQIK----ECR--GRRQ-ESQALVLQNAHVICCTLSTSGSIVLENAFRRLG 1951
Chicken 2231 --QQLASQLK----EVR--GHSQ-KVQTDIILESDIICCTLSTSGGGLLESAFWRQG 2278
Whale shark 2241 --EELGRKLK----ELR--GRTQ-DIQLNIILESHIICCTLSTCGIMLLEMSFRKLG 2288
Peters' clawed toad 2021 --QLLANSLK----EVR--RRPQ-ELQRTIILESHVICCTLSTSGGILLESAFRQLG 2068
Breadcrumb sponge 1968 --SKLMTQHKNLKVQCGSMQRDLSKVRKRILMDADVVCATLCGSGSYNLWECV---- 2018
Cauliflower coral 1640 --RTLQKKFREQRAEQANKRRKESSIRKEILYRADIICTTLSGAGSTSLKQDL--EG 1692
Acorn worm 1307 --NELEKETKKSSSNARNLRGQESKVRQHVLVEADVVCCTLNGSGSSVLDSAV-RMS 1360
Brook lamprey 2245 --EILNHTIKEMRSQKWI------VQKKLIVESNIICCTLSTSGMQILETAMR--G 2290
Florida lancelet 2210 --DKLDKTNRQVQLKNEERHREGTKIRQRVLMDADVVCCTLSASGSNMLEQAITKAG 2264
*S. cerevisiae* Sen1 1534 --DEMR---EKNSVNYRNRDLDRRNAQAHILAVSDIICSTLSGSAHDVL------AT 1579
*C. thermophilum* Sen1 1511 IGAKIDADKASGNTYARETEIKRRQIQQEILDKAQVLCATLSGSGHEMF------KN 1561
<40 40-60 60-80 >80
Identity [%]

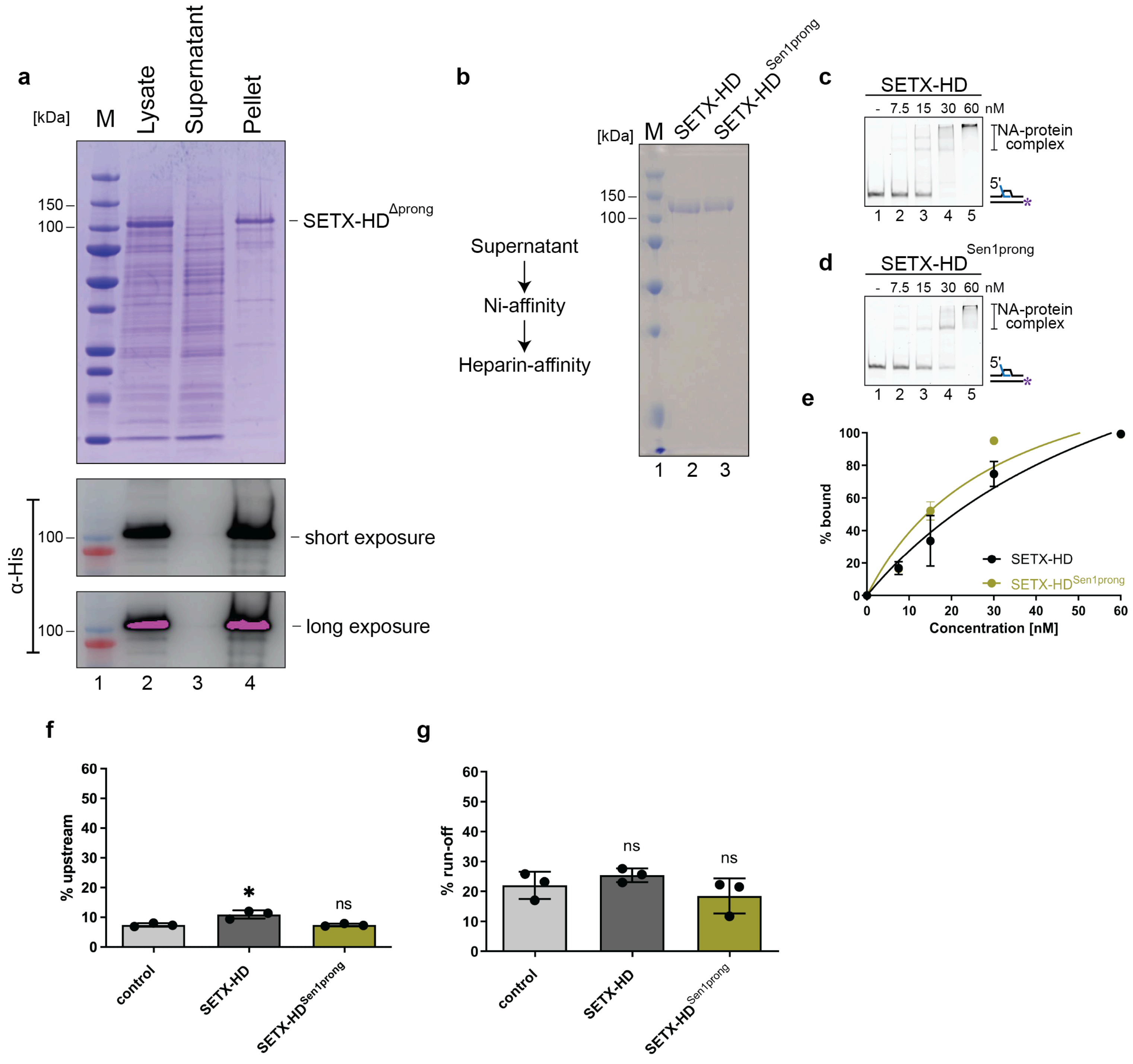
a
[kDa]
M
Lysate
Supernatant
Pellet
150
100
SETX-HD^Δprong
α-His
100
short exposure
100
long exposure
1 2 3 4
b
Supernatant
Ni-affinity
Heparin-affinity
[kDa]
M
SETX-HD
SETX-HD^Sen1prong
150
100
1 2 3
c
SETX-HD
- 7.5 15 30 60 nM
NA-protein complex
5'
1 2 3 4 5
d
SETX-HD^Sen1prong
- 7.5 15 30 60 nM
NA-protein complex
5'
1 2 3 4 5
e
% bound
Concentration [nM]
SETX-HD
SETX-HD^Sen1prong
f
% upstream
*
ns
control
SETX-HD
SETX-HD^Sen1prong
g
% run-off
ns
ns
control
SETX-HD
SETX-HD^Sen1prong

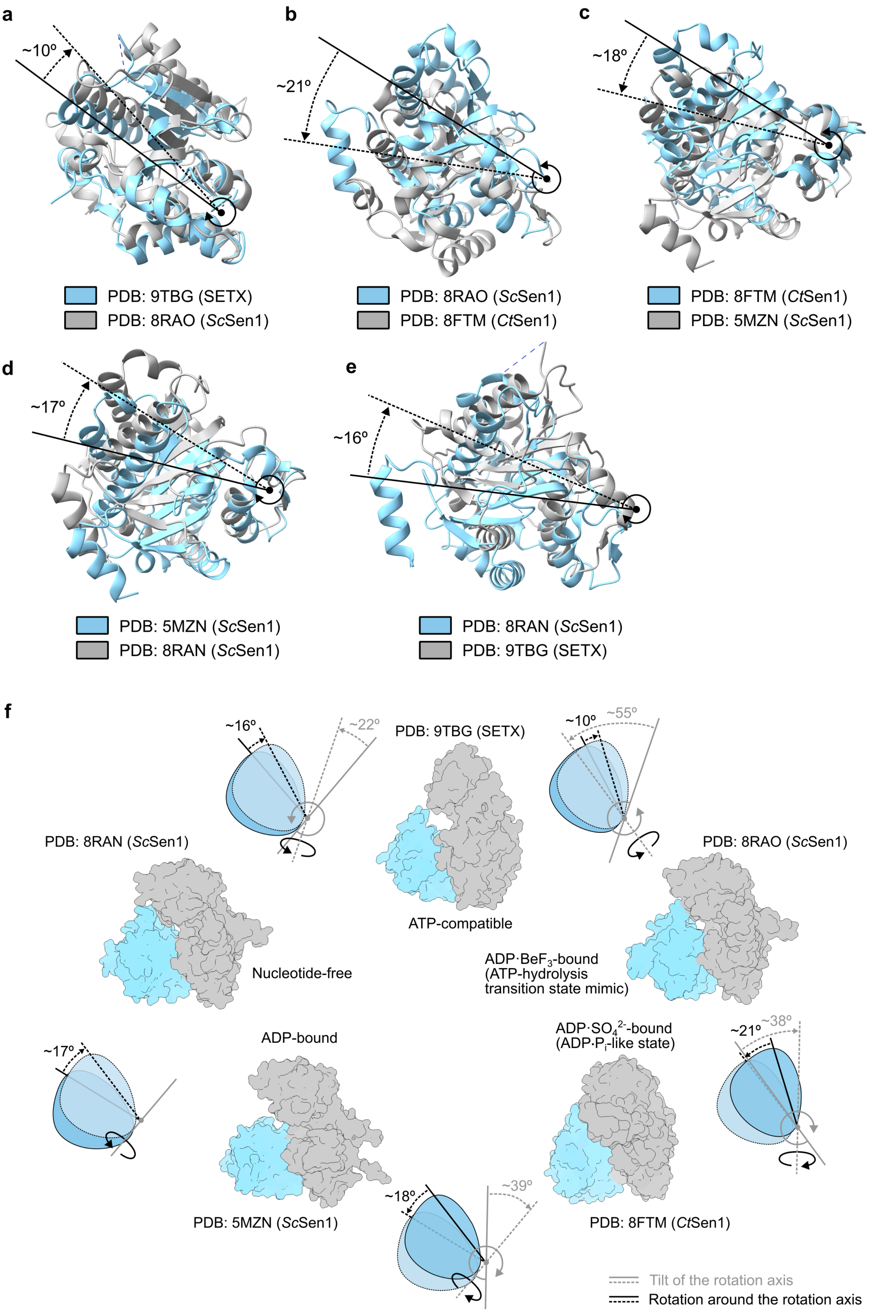
a
~10º
PDB: 9TBG (SETX)
PDB: 8RAO (ScSen1)
b
~21º
PDB: 8RAO (ScSen1)
PDB: 8FTM (CtSen1)
c
~18º
PDB: 8FTM (CtSen1)
PDB: 5MZN (ScSen1)
d
~17º
PDB: 5MZN (ScSen1)
PDB: 8RAN (ScSen1)
e
~16º
PDB: 8RAN (ScSen1)
PDB: 9TBG (SETX)
f
~16º
~22º
PDB: 9TBG (SETX)
~10º
~55º
PDB: 8RAN (ScSen1)
PDB: 8RAO (ScSen1)
ATP-compatible
Nucleotide-free
ADP·BeF3-bound
(ATP-hydrolysis
transition state mimic)
ADP-bound
ADP·SO4 2--bound
(ADP·Pi-like state)
~17º
~21º
~38º
~18º
~39º
PDB: 5MZN (ScSen1)
PDB: 8FTM (CtSen1)
Tilt of the rotation axis
Rotation around the rotation axis
